# Open-source Photobleacher for Fluorescent Imaging of Large Pigment-Rich Tissues

**DOI:** 10.1101/2025.02.24.639965

**Authors:** Tatsuya C. Murakami, Nick Belenko, Griffin Dennis, Cuidong Wang, Maria Esterlita Siantoputri, Yurie Maeda, Christina Pressl, Nathaniel Heintz

## Abstract

Fluorescent imaging enables visualization of the specific molecules of interest with high contrast, and the use of multiple fluorophores in a single tissue sample allows visualization of complex relationships between biological molecules, cell types, and anatomy. The utility of fluorescent imaging in human tissue has been limited by endogenous pigments that can block the light path or emit an autofluorescence, thereby interfering with the specific imaging of target molecules. Although photobleachers have been developed to quench endogenous pigments, the lack of customizability limits their utility for a broad range of applications. Here, we present a high luminous-intensity photobleacher that is based on rigorous simulations of illumination patterns using the laws of radiation, along with the framework to maximize bleaching efficiency. This open-source project is designed to help researchers customize and scale according to the tissue types and the research goals. The photobleacher is applicable to both thin tissue slices and large-volume cleared tissue samples to enable serial three-dimensional imaging of postmortem human brain using multiplexed antibody or oligonucleotide probes.

**SIGNIFICANCE STATEMENT:** Photobleaching is an effective technique for quenching endogenous pigments, enabling multiplexed fluorescent imaging of pigment-rich tissues, such as postmortem human samples. While many photobleaching strategies have been proposed, there is no standard guidance on how to design and use a photobleacher. This study introduces a general strategy for designing an effective, scalable, and customizable photobleacher, and proposes a workflow for properly treating tissues with the photobleacher. The technique enables high-contrast molecular visualization in tissues of various sizes, including large volumetric cleared tissues. Our framework will accelerate the quantitative understanding of human molecular anatomy and is applicable to diverse biological fields, including medical diagnostics.

## INTRODUCTION

Advances in fluorescence imaging have shed light on many biological phenomena that were previously invisible. Despite these advances, serious problems arising from endogenous pigments such as heme and autofluorescence from lipofuscin pose significant problems for multiplexed fluorescence imaging of human tissue samples. This has limited the utility of fluorescent imaging in human tissue despite its importance in studies of molecular neuroanatomy and pathogenesis.

In human nervous tissues, pigments include lipofuscin, a byproduct of lipid peroxidation that accumulates with age and exhibits broad-spectrum fluorescence (Mann et al., 1978; Gray and Woulfe, 2005). Other key metabolic molecules, such as NADH, flavins, and hemoglobin and its breakdown products, also produce autofluorescent signals that complicate imaging (Duong and Han, 2013). Various strategies have been proposed to mitigate the issue of autofluorescence, with quenching agents being widely employed for this purpose. Among the chemicals used are copper sulfate, ammonium chloride, Sudan Black B, and sodium borohydride. While these chemical treatments can enhance the specificity of fluorescence imaging, the chemical modification of the tissue can obscure true signals or compromise tissue integrity (Duong and Han, 2013). Furthermore, their use requires optimization to achieve the best signal-to-noise ratio, but this is often challenging because optimal conditions vary depending on the size and type of tissue. Given the heterogeneous shape of human tissues and the growing demand for fluorescent imaging of multiple types of human organs, a quenching technique that does not require extensive optimization for its application for each tissue type would enable more efficient and reproducible human molecular neuroanatomy.

To circumvent the limitations of chemical quenching of autofluorescence, photobleaching is increasingly gaining attention. Few adverse effects of photobleaching have been reported with respect to signal-to-noise ratio in fluorescence imaging (Sun and Chakrabartty, 2016; Pigoli et al., 2019; Tsuneoka et al., 2022). To exploit the advantages of photobleaching, several research groups have proposed photobleaching devices (hereafter photobleachers), as summarized in **Table S1**. Although fluorescent tubes were initially used as a light source for photobleaching (Neumann and Gabel, 2002), arrayed light-emitting diodes (LEDs) are now predominantly employed due to their lower heat generation. Current photobleaching setups are either based on repurposing a generic desk lamp or floodlight (Duong and Han, 2013; Sun and Chakrabartty, 2016; Pigoli et al., 2019) or are fully custom-made (Ku et al., 2020; Tsuneoka et al., 2022; Park et al., 2024). Only one pre-assembled photobleacher is commercially available (Tsuneoka et al., 2022).

The designs of these photobleachers were often heuristically determined, and the processes used to reach these designs have not been fully disclosed. Thus, while the density of the LEDs and the distance between the biological specimen and the LEDs are important parameters that control the efficiency of bleaching, the rationale behind the choice of the configuration in previous studies is not discussed. As a result, there is limited accumulated knowledge on how to choose and assemble light sources, as well as how to position the samples to be bleached. Moreover, the published photobleachers were designed to satisfy specific applications, requiring redesign of the device without standard instructions if new applications are planned.

For example, the commercially available photobleacher, TiYO (Nepagene, Japan), is excellent for photobleaching small, thin tissue slices but is not appropriate for large, volumetric cleared tissue. Therefore, we see an urgent need for a fully open-source photobleacher, along with clear guidance on how to design, assemble, and properly use it.

In this study, we describe an open-source photobleacher that can rapidly quench the autofluorescence of fixed tissues using arrayed high-power LEDs. The design of the photobleacher is based on optical simulations of critical parameters and is optimized for both thin tissue slices and cleared large volumetric tissues. To enable fabrication of the photobleacher, we have included detailed assembly instructions in this manuscript (**Supporting text**). We designed the photobleacher to be customizable, allowing users to flexibly adjust the device’s capacity. Using lipofuscin-rich human nervous tissue, we demonstrated the applicability of the photobleacher for two most common molecular staining techniques: fluorescent in situ hybridization and fluorescent immunohistochemistry. The staining results indicate little to no impact on the signals while successfully suppressing autofluorescence. We explored the impact of the immersion medium during photobleaching, providing the users with concrete guidance on how to utilize the photobleacher. We also demonstrated the device’s applicability in 3D staining using a piece of human brain and a blood-rich mouse organ, proving that the photobleaching process can be easily incorporated into many common molecular staining techniques.

## RESULTS

### Simulation of the total illuminance from multiple light sources

Photobleaching is an irreversible process in which fluorophores, upon repeated absorption of excitation photons, undergo chemical modifications that result in a permanent loss of their fluorescence. One pathway leading to photobleaching involves intersystem crossing from the excited singlet state to the excited triplet state of the fluorophore. The triplet state has significantly longer lifetimes than the singlet state (Widengren and Rigler, 1996). This extended lifetime, coupled with the high chemical reactivity of the triplet state, increases the probability of interactions with surrounding molecules, particularly molecular oxygen, and can direct chemical modifications of the fluorophore, ultimately resulting in the destruction of its fluorescent properties. Generally speaking, the reception of more photons will increase the probability of the loss of the fluorophore (Deschenes and Vanden Bout, 2002).

If the object to be bleached is small, ranging from ∼100 µm to a few millimeters in size with limited thickness (which is comparable to the field of view of fluorescent microscopy) laser-based photobleaching is a common option (Lubeck et al., 2014; Chen et al., 2015). For larger tissues, the use of laser is not feasible due to the long duration required to cover wider areas. In larger tissue sample, the use of a spatially divergent light source such as an LED is preferred. If a single light source is used, there are only two parameters that control how fast the photobleaching can proceed: the intensity of the light source and the distance to the specimen. In this case, the best photobleaching strategy is to use a stronger light source and place the specimen close to the light source. The limitation of this approach is that one cannot photobleach multiple specimens at a time, and the size of the specimen must be similar to the size of the light source. For this reason, many published photobleachers exploit the use of multiple light sources (**Table S1**), but establishing the best photobleaching strategy is more complex as the alignment of the light sources can affect the brightness at a given position. The simple strategy of placing the strong light sources as dense as possible will generate excessive heat and destroy the specimen or device itself. These considerations motivated us to simulate how the alignment of light sources affects the brightness at each position with the aim of finding optimal parameters that can achieve evenly distributed and strong illumination in a benchtop device.

There are two ways to express optical metrics: radiometric quantities and photometric quantities. Radiometric quantities measure the total electromagnetic energy across all wavelengths, while photometric quantities are weighted by the human eye’s sensitivity to visible light; the two are mutually convertible provided that the wavelength properties of the light source are known. In this manuscript, we will use photometric terminology, such as luminous intensity and illuminance, for simulation and explanatory purposes. We note that all conclusions would remain the same if radiometric quantities were used.

We calculated the illuminance at the observation point with the position **p**, given an arrayed illumination setup composed of light sources with identical optical properties, based on the fundamental principles of optics (**Fig. 1A)**. The illuminance at the observation point from a single light source follows the inverse-square law, which can be expressed as

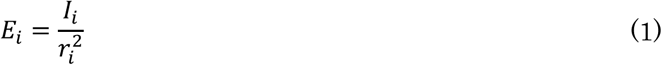

**Figure 1.**
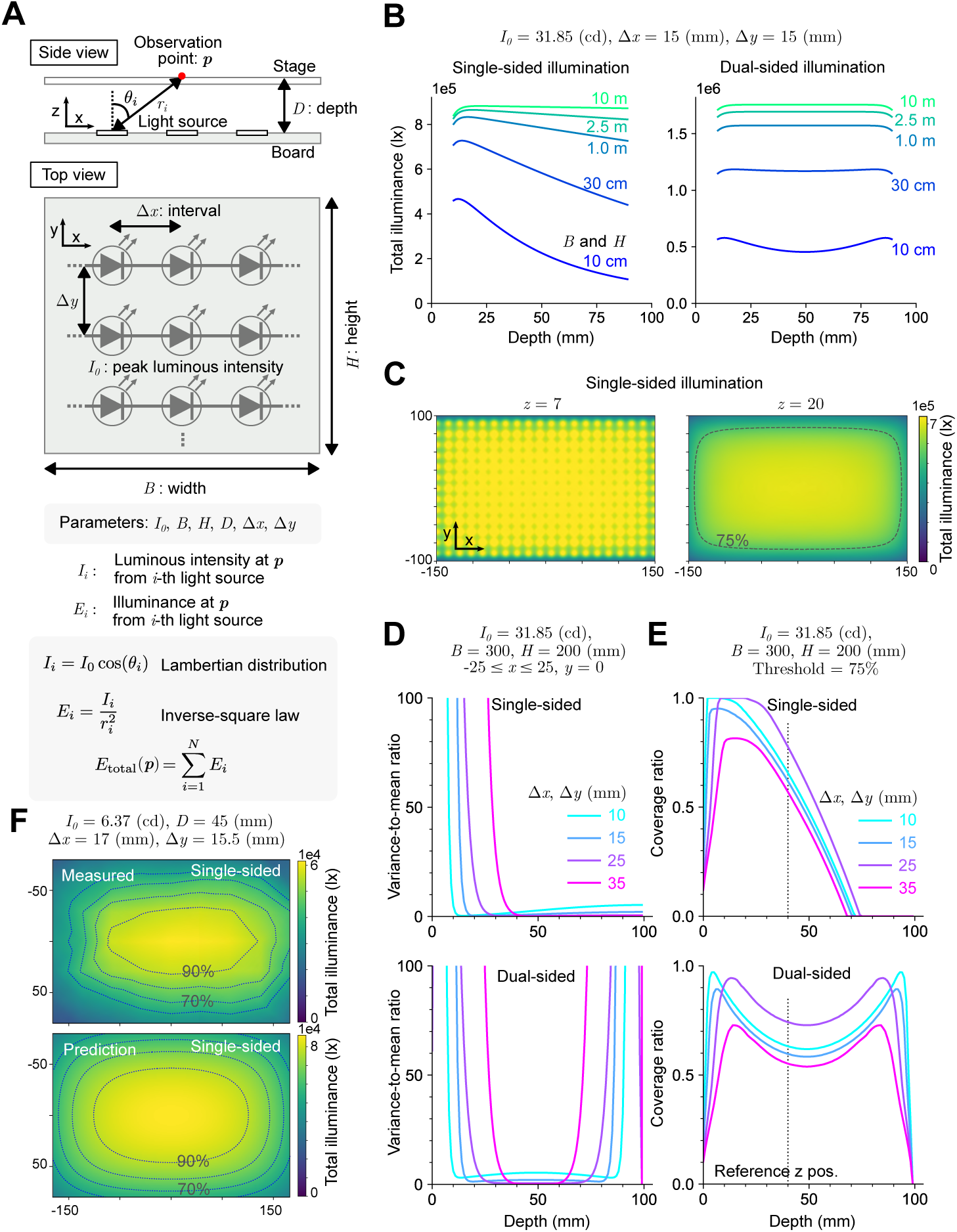
Simulation of total illuminance from multiple light sources. (**A**) Schematic representation of the parameters used in the simulation. The optical model and laws applied in the simulation are shown at the bottom. (**B**) The effect of board size on the illuminance decay along the depth. Left: single-sided illumination; Right: dual-sided illumination. (**C**) Heatmap visualization of illuminance at specified depths. The 75% contour of peak illuminance is highlighted in a gray dot line on the right. (**D**) Variance-to-mean ratio of illuminance along the x between -25 mm and 25 mm at y = 0 mm, plotted as a function of depth. (**E**) Coverage ratio of the area where illuminance exceeds 75% of the center value relative to the total board size. (**F**) The comparison of the measured illuminance (top) and the prediction (bottom). The 90%, 80%, 70% and 60% contours of peak illuminance is highlighted in a blue dot lines. See also **Figures S1**.

where the *E*_*i*_ represents the illuminance (in lux, lx) from the *i*-th light source, *I*_*i*_ denotes the luminous intensity (in candela, cd), and *r*_*i*_ is the distance (in meters, m) between the observation point and the light source. We note that the *E*_*i*_, *I*_*i*_, and *r*_*i*_ are all dependent on *p*. The total illuminance at the observation point from all light sources is given by

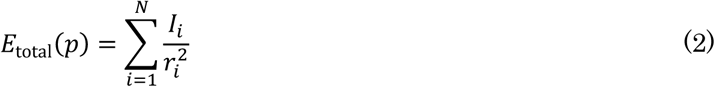

where *N* is the total number of light sources. We assumed a layout where the light sources are aligned at constant intervals, with *Δx* as the interval in one direction and *Δy* in the orthogonal direction, all located on the same plane as shown in **Fig. 1A**. The light sources were positioned on a rectangular board with height *H* and width *B*. The shortest distance between the observation point and the board was defined as depth *D*. We denoted the x-, y-, and z-axes as corresponding to the width, height, and depth directions, respectively, with the center of the board set at *x* = 0, *y* = 0, and *z* = 0. The observation point is constrained within the range −*B*/2 ≤ *x* ≤ *B*/2, −*H*/2 ≤ *y* ≤ *H*/2 and *z* = *D*. Luminous intensity is typically influenced by the angle at which the light source is viewed. To account for this angular dependency in our simulation, we assumed a Lambertian distribution, as given by

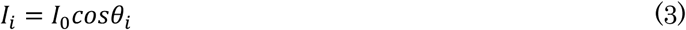

where *I*_0_represents the peak luminous intensity of the light source, and *θ*_*i*_ is the angle between the observation point and the light source. Many common light emitters, such as LEDs, follow this Lambertian distribution. From the geometry, the *cosθ*_*i*_ is given by

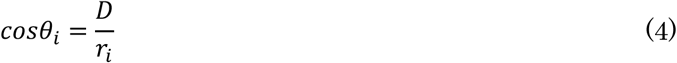

From the (1) ∼ (4), we get

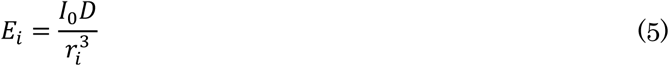

And

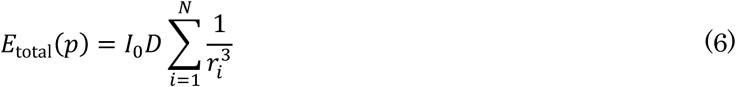

The aim of this simulation is to determine the optimal parameters—*I*_0_, *B*, *H*, *D*, *Δx* and *Δy* —that maximize the total illuminance and retain even illumination over the sample stage, while ensuring that the parameters remain within realistic physical scales. In practical scenarios, biological samples can extend up to several centimeters, and it is often necessary to bleach multiple samples simultaneously. The illumination thus has to cover the relatively large area of the sample stage. Additionally, when the photobleacher is used for large volumetric objects, such as a cleared human organ, uniform illumination in the z-direction, as well as in the x- and y-directions, is needed.

From equation (6), it is obvious that the total illuminance is proportional to the peak luminous intensity *I*_0_ of the light sources but not affecting the pattern of the illuminance on the sample stage. We thus simulate how *B*, *H*, *D*, *Δx* and *Δy* affect the total illuminance. Note that the distance *r*_*i*_ can be easily calculated using basic geometry from the provided parameters. For instance, when the coordinates of the observation point are *x* = 0, *y* = 0, and *z* = *D*, the right-hand side of equation (6) can be written as

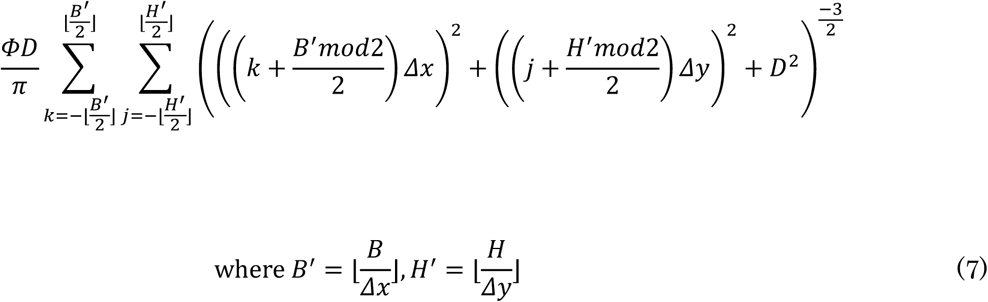

It is conceivable that smaller values of *Δx* or *Δy* will contribute more to the total illuminance, but it is not immediately clear how changing *Δx* or *Δy* affects the total illuminance from equation (7). By fixing *H* at 200 mm, *B* at 300 mm, and *D* at 20 mm, we calculated the total illuminance at the center of the sample stage while varying *Δx* and *Δy* (**Fig. S1A**). The size of *Δx* or *Δy* is almost inversely proportional to the total illuminance. This result can be intuitively understood by considering that halving *Δx* (or *Δy*) effectively doubles the number of light sources within the same area, which is essentially the same as doubling the luminous intensity of the light source. If there is a large difference between *Δx* and *Δy*, the stripe-pattern “hot spot” becomes more visible (**Fig. S1B**) where the areas gain more photons than other areas, resulting in uneven photobleaching. For this reason, we only considered the case where *Δx* equals *Δy* in the following simulations. Next, we examined the decay of the total illuminance as the depth increased (**Fig. S1C**). The results indicated an approximately linear decay of the total illuminance with depth. One simple solution to address this uneven illumination in the z-direction is to increase *Δx* and *Δy* (**Fig. S1C**). However, this strategy is not ideal, as increasing *Δx* and *Δy* can significantly reduce the total illuminance, as discussed earlier. We decided to explore an alternative solution to prevent the decay of illuminance in the z-direction.

It is known that if the size of the board is sufficiently large compared to the depth, the board can be approximated as an infinitely large Lambertian plane where the illuminance becomes constant across depth ranges. By fixing the observation point at *x* = 0, *y* = 0 and varying the depth of the sample stage from 10 mm to 90 mm, we calculated the minimum board size required to make illuminance independent of depth (**Fig. 1B**). We found that when both the height and width are equal to 10,000 mm (10 m), the illuminance becomes nearly constant in the depth range between 10 mm and 90 mm. However, this solution is not practical because a 10-meter-wide board is too large for a typical laboratory environment. An alternative approach is to place an additional light source to compensate for the decay of illuminance in the z- direction. Since the decay is approximately linear to the depth, the loss can be easily compensated by adding another set of light sources positioned to illuminate the specimen from behind. We found that a board size of 300 mm for both *H* and *B* is sufficient to achieve uniform illuminance in the z-direction when we place another array of light sources (**Fig. 1B**). This trend holds true for both *Δx* and *Δy* values of 15 mm and 30 mm (**Fig. S1D**). We considered the situation with the board size of 200 mm for *H* and 300 mm for *B*, with the 100 mm interval between two arrays when dual-illumination was used.

If the sample stage is positioned too close to the light sources, hot spots appear with patterns that reflect the arrangement of the light sources (**Fig. 1C**). These hot spots also appear in simulations of a published photobleacher (Tsuneoka et al., 2022) (**Fig. S1E**). Keeping the specimen further away from the light sources can eliminate the hot spots, but it is unclear at what depth the illuminance becomes uniform. To address this question, we calculated the variance of the illuminance at various depths (**Fig. 1D**). There is a certain depth at which the hot spots disappear.

This depth depends on the interval between light sources, with shorter intervals being more effective at diminishing the hot spot effect. When *Δx* is 15 mm, the illumination becomes almost uniform at a depth of 20 mm.

Another consideration is the decay of illuminance at the edges (**Fig. 1C**). Generally, the position at *x* = 0 and *y* = 0 has the highest illuminance, and the illuminance drops sharply as the position moves closer to the edges of the sample stage. To simulate how large an area is uniformly illuminated, we set a threshold of 75% of the peak illuminance at a given depth and investigated the how large area is beyond this threshold criteria. We found that the coverage ratio above this threshold is more than 0.6 at a depth of 40 mm with a *Δx* of 15 mm (**Fig. 1E**). The illuminated area extends approximately 240 mm in width and 160 mm in height (**Fig. 1C**), which is large enough to accommodate multiple biological specimens.

To verify the simulation, we assembled an array of LEDs and measured the total illuminance on a plane at 45 mm from the array. The illuminance patterns were similar between the measured and predicted values, demonstrating the utility of the simulation (**Fig. 1F**). There was a discrepancy in the absolute values of the illuminance, with measured values being approximately 30% lower than the predicted values. This discrepancy is likely due to the fact that the theoretical maximum power listed in an LED datasheet is often not achievable in practice. The internal resistance of the LED can lead to energy losses in the form of heat, which results in a decrease in brightness. Regardless of the limited capability to predict the absolute illuminance, the simulation is very useful to overview the evenness of the illuminance in a given space.

### Design of an open-source photobleacher

Considering the simulation results mentioned above, we developed an open-source photobleacher. We selected LEDs as a light source because of their low cost and high energy conversion efficiency. A detailed explanation of how we selected our design is provided in the **Supporting text**. The total size of the board is 297.6 mm in width and 197.6 mm in height, with the distance between the facing boards set at 110 mm (**Fig. 2A**). This size was chosen to ensure the device can easily fit into a standard laboratory environment. To accommodate the demand for a larger photobleacher, we incorporated customizability, allowing users to increase the capacity of the device. We designed a scalable printed circuit board (PCB) on which the LEDs are mounted. The modular PCB design allows users to extend the board size easily by simply connecting multiple boards, either horizontally or vertically (**Fig. 2B**). The intervals between the LEDs are set at 12.5 mm. After running the device continuously for over 4 hours with the use of cooling fans, we found that the temperature at the sample stage never exceeded 32°C, ensuring that the temperature remained stable at this spacing.

**Figure 2.**
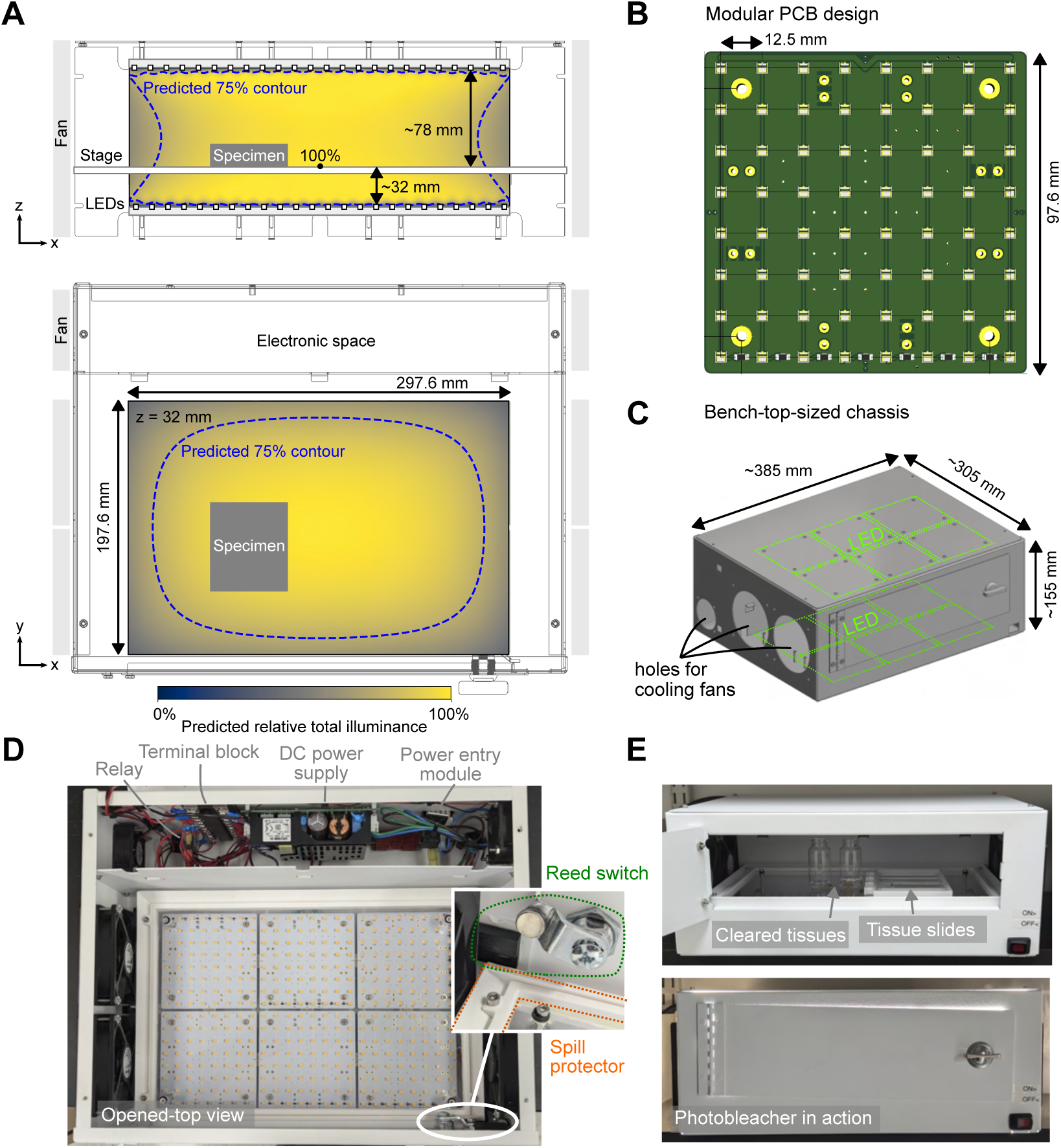
Design of the open-source photobleacher. (**A**) Orthographic sketches of the photobleacher, showing the chassis wireframes along with simulation results in a heatmap. Top: side view; Bottom: top view. (**B**) Diagram of the modular printed circuit board (PCB) design, which is scalable by connecting boards both vertically and horizontally. (**C**) Illustration of the bench-top- sized chassis. (**D**) Overview of the completed photobleacher from the top. (**E**) Photobleacher in operation with cleared tissues and tissue slides.

We conducted a simulation of the illuminance using the dimensions of our photobleacher, confirming that the illuminance is uniform in most areas of the chamber (**Fig. 2A**). The chassis was designed to be as compact as possible, with no dimension exceeding 40 cm, while still accommodating the power modules (**Fig. 2C and 2D**). For cooling, we decided to use four cooling fans for the illumination chamber and two smaller fans for the electronic space. The cooling system is based on simple air cooling without heat sinks, and we positioned the fans on both sides of the device to maximize airflow. We implemented a reed switch that automatically turns off the device when the door is accidentally opened while the device is active. We also included spill protection to prevent any solution from dripping onto the LEDs, which helps protect the photobleacher from short-circuiting (**Fig. 2D**). Thanks to the large size of the illumination chamber, users can place a biological specimen with clearing solution in a glass vial. The device is designed to support both bleaching of large volumetric tissues and tissue slides (**Fig. 2E**). This project is fully open-source and completely customizable to meet users’ needs, such as board size, light colors, and illumination power. We have included assembly instructions, which extensively describe the step-by-step building process, as well as the wiring diagram in this manuscript, so that researchers without prior experience can replicate the device (**Fig. S2**). The latest version is available on GitHub (https://github.com/NBelenko/Photobleacher).

### Efficiency of photobleaching

We used human brain slices to characterize the rate of photobleaching. Using dried fresh-frozen human tissue slices in air, we performed photobleaching over 72 hours and measured the intensity of lipofuscin autofluorescence with confocal microscopy. The results indicate that the majority of the autofluorescence was largely quenched after 72 hours (**Fig. S3A**). Excitation with longer visible wavelengths (555 nm and 639 nm) showed almost negligible autofluorescence, while excitation with a shorter visible wavelength (488 nm) yielded only a minor level of autofluorescence. Excitation in the ultraviolet (405 nm) was unaffected by photobleaching, presumably due to the lack of UV irradiation in our LED light source. This observation is consistent with a previous report (Tsuneoka et al., 2022). We did not explore the quenching of UV autofluorescence further, as UV excitation is often used for counterstaining (e.g., DAPI), and the signal intensity can be easily adjusted to mask autofluorescence by increasing the concentration of the staining molecules. We confirmed a decrease in autofluorescence with increasing irradiation time in either dried-fresh frozen slices or formalin-fixed paraffine-embedded (FFPE) slices (**Fig. S3B**).

We next examined the impact of the immersion medium during photobleaching. We compared five immersion media: air, ethanol, mineral oil, mixture of benzyl alcohol and benzyl benzoate (BABB), and phosphate buffer saline (PBS), using paraformaldehyde (PFA)-fixed human cortical tissue slices. BABB is a common clearing solvent composed. We confirmed a reduction in lipofuscin autofluorescence in all media with the greatest reduction in PBS (**Fig. 3A and 3B**). These differences among the media may reflect how much the medium can retain oxygen surrounding the fluorophore. In tissues photobleached in ethanol, mineral oil, and BABB, we observed an increase in fluorescence in the extracellular matrices (**Fig. 3C**). We reasoned that this increase was due to tissue shrinkage caused by dehydration. The shrunk tissue compacts the density of the matrices, thereby enhancing the background fluorescence.

**Figure 3.**
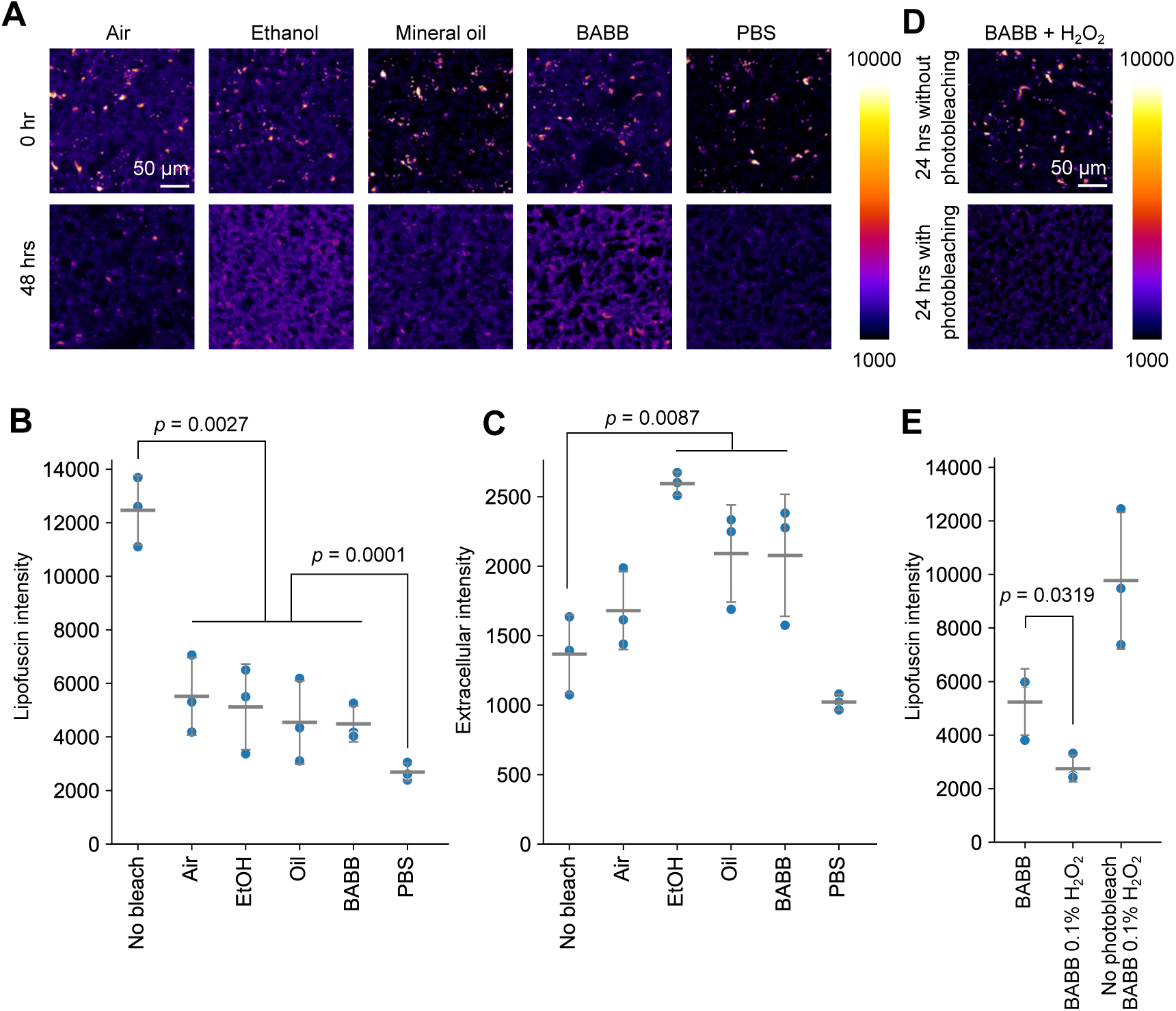
Photobleaching in various immersion media. (**A**) Human cortical slices before and after photobleaching in various immersion media. The images are shown in a pseudocolor according to the intensities. (**B**) Quantification of the averaged lipofuscin intensity (N = 3). Welch’s t-test. (**C**) Quantification of the averaged intensity of the extracellular areas (N = 3). Welch’s t-test. (**D**) Human cortical slices before and after photobleaching in BABB with hydrogen peroxide. The images are shown in a pseudocolor according to the intensities. (**E**) Quantification of the averaged lipofuscin intensity after photobleaching in BABB or BABB with hydrogen peroxide (N = 3). Student’s t-test. For **A** ∼ **E**, excitation with 555 nm laser was used. Errorbars indicate standard deviations. EtOH: ethanol.

We next explored the possibility of increasing the amount of reactive oxygen by adding hydrogen peroxide (H2O2) to the medium. Upon strong illumination, hydrogen peroxide is known to produce hydroxyl radicals via photolysis (Baxendale and Wilson, 1957). These hydroxyl radicals subsequently degrade the fluorophore. This process has been used to remove pigments from large biological tissues (Darche et al., 2023). We dissolved a small amount of H2O2 in BABB and examined whether the addition of H2O2 further promotes photobleaching. We confirmed a significant reduction in autofluorescence after the addition of H2O2 when photobleached (**Fig. 3D and 3E**). It is important to note that the combination of H2O2 and photobleaching is possible in hydrophobic environment by dissolving H2O2 in BABB. This eliminates water from the medium, which can otherwise cause hydrolysis of molecules of interest, such as RNA. Another important implication is that the photolysis process can be achieved in cleared volumetric tissue.

### Application of the photobleacher to lipofuscin-rich human brain tissue slices

Photobleaching has a significant advantage over chemical-based quenchers, such as Sudan Black B, which are known to reduce not only autofluorescence signals but also target signals of interest (Sun and Chakrabartty, 2016; Pigoli et al., 2019; Tsuneoka et al., 2022). We aimed to determine whether our photobleacher could selectively reduce autofluorescence without affecting the target fluorescence signals. To assess this, we tested two common molecular staining methods: fluorescent in situ hybridization (FISH) and immunohistochemistry (IHC). For FISH, we used the commercially available RNAscope kit to probe tissue slices from human cerebral cortex with and without photobleaching, targeting two mRNAs: *Pcp4* (Purkinje cell protein 4) and *Htr2c* (5-hydroxytryptamine receptor 2C). Both genes are predominantly expressed in the deeper cortical layers, with some cells co-expressing both genes. The tissue slices were prepared from fresh-frozen archival samples obtained from Miami’s Brain Endowment Bank, briefly fixed with paraformaldehyde, and then subjected to photobleaching. For labeling, we used TSA Vivid Fluorophore 570 and 650 dyes. In the slice with photobleaching, we confirmed that the signals appeared predominantly in the deep cortical layers, which is consistent with the gene expression patterns. On the other hand, the slice without photobleaching showed the signals throughout the cortical layers (**Fig. 4A**). Higher magnification views revealed that the intense autofluorescence was contaminating the visible fluorescence channels, potentially leading to over-estimation of the number of the cells expressing the genes. In contrast, photobleaching effectively quenched the autofluorescence, allowing for a clearer identification of cells that express one or both genes (**Fig. 4B**). We observed no noticeable signal reduction after the photobleaching.

**Figure 4.**
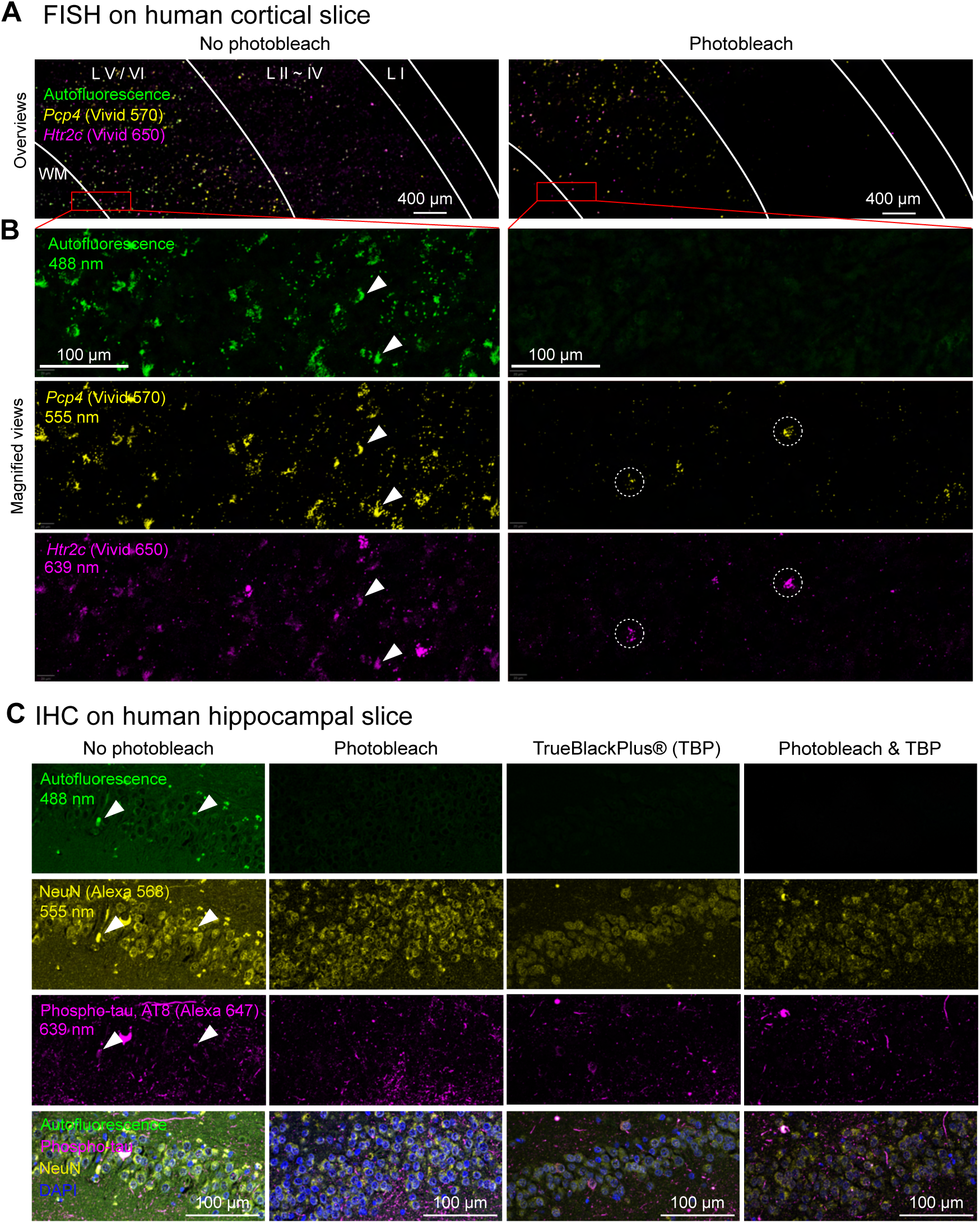
Molecular staining of human tissue slices after photobleaching. (**A**) Overview of the fluorescent in situ hybridization (FISH) targeting two types of transcripts, *Pcp4* and *Htr2c*, in thin slices of PFA-fixed human cerebral cortex. Left: tissue slice without photobleaching. Right: tissue slice with photobleaching. The approximated the borderlines of the cortical laminar are indicated by white lines. WM: white matter. (**B**) Magnified views of the red squares from panel **A**. The arrowheads indicate possible cross-talk between autofluorescence and visible wavelengths. The dotted circles highlight cells exhibiting double-gene expression, without confounding from autofluorescence. (**C**) Immunohistochemistry (IHC) against NeuN and phospho-tau in the hippocampal dentate gyrus from a donor with Alzheimer’s disease. Thin slices were made from FFPE tissue blocks. The arrowheads indicate the possible cross-talks of the autofluorescence to the visible wavelength.

For IHC, we used anti-NeuN and anti-phospho-tau antibodies to stain hippocampal tissues from Alzheimer’s disease (AD) patients. We used Alexa Fluor 568 and Alexa Fluor 647 respectively for fluorescent labeling. The tissue slices were prepared from FFPE blocks obtained from Miami’s Brain Endowment Bank and were photobleached before de-paraffinization in air. In a slice without photobleaching, autofluorescence masked the true fluorescence signals across all visible channels, complicating the interpretation of the results. However, photobleaching largely alleviated this issue (**Fig. 4C**). For comparison, we also included slices treated with TrueBlack Plus, a water-soluble alternative to Sudan Black B. Although TrueBlack Plus successfully reduced autofluorescence, it caused a noticeable decrease in the signal intensity compared to slides without treatment. In contrast, photobleaching alone had little to no impact on the signal strength while effectively mitigating autofluorescence interference.

### Application of the photobleacher to cleared pigment-rich organs for three-dimensional imaging

Single-cell-resolution three-dimensional imaging of autofluorescence-rich tissues, such as human organs, is crucial for understanding the pathogenesis of neurological disorders (Murray et al., 2015; Liu et al., 2016; Nojima et al., 2017; Lai et al., 2018; Morawski et al., 2018; Hildebrand et al., 2019), and the development of the urinary tract and vascular systems (Tainaka et al., 2018; Zhao et al., 2020; Mai et al., 2022). As the demand for three-dimensional imaging rises, so does the need for effective photobleaching of autofluorescence. There have been successful applications of photobleaching in 3D-cleared tissues (Darche et al., 2023; Park et al., 2024; Zheng et al., 2024), but the effects and limitations of this technique have not been thoroughly discussed. First, we prepared a PFA-fixed mouse heart, known to be heme-rich and have high autofluorescence. The tissues were cleared using BABB after methanol-based dehydration, and we assessed the efficacy of photobleaching in the clearing medium using light-sheet microscopy (**Fig. 5A**). We examined two photobleaching conditions: regular photobleaching in BABB and photolysis-assisted photobleaching in BABB with the addition of H2O2. Both photobleaching methods resulted in a drastic reduction of autofluorescence (**Fig. 5B**). After photolysis- assisted photobleaching in BABB, the heart tissue was rehydrated and subjected to immunostaining using a modified iDISCO protocol. We stained for alpha-smooth muscle actin (αSMA) using an anti-αSMA antibody conjugated with Alexa Fluor 647, which primarily stains mature myofibroblasts in the heart (Shinde et al., 2017). The staining clearly highlighted the vasculature of the heart in 3D, demonstrating the effectiveness of the photobleaching method (**Fig. 5B and 5C**). Next, we examined the effect of photobleaching in lipofuscin-rich human brain tissue. Sub-centimeter-sized pieces of tissue were dissected from a fresh-frozen cerebral cortex obtained from Science Care, and was subjected to the photobleaching after PFA-fixation and clearing. We found that both photobleaching and photolysis-assisted photobleaching effectively reduced lipofuscin autofluorescence (**Fig. 5D**). The tissue after photolysis- assisted photobleaching was subjected to immunostaining targeting αSMA and glial fibrillary acidic protein (GFAP) using antibodies conjugated with Alexa Fluor 647 and Cy3 respectively. The resulting tissue showed specific staining with high contrast (**Fig. 5E**), demonstrating the utility of our photobleaching technique in volumetric human tissues.

**Figure 5.**
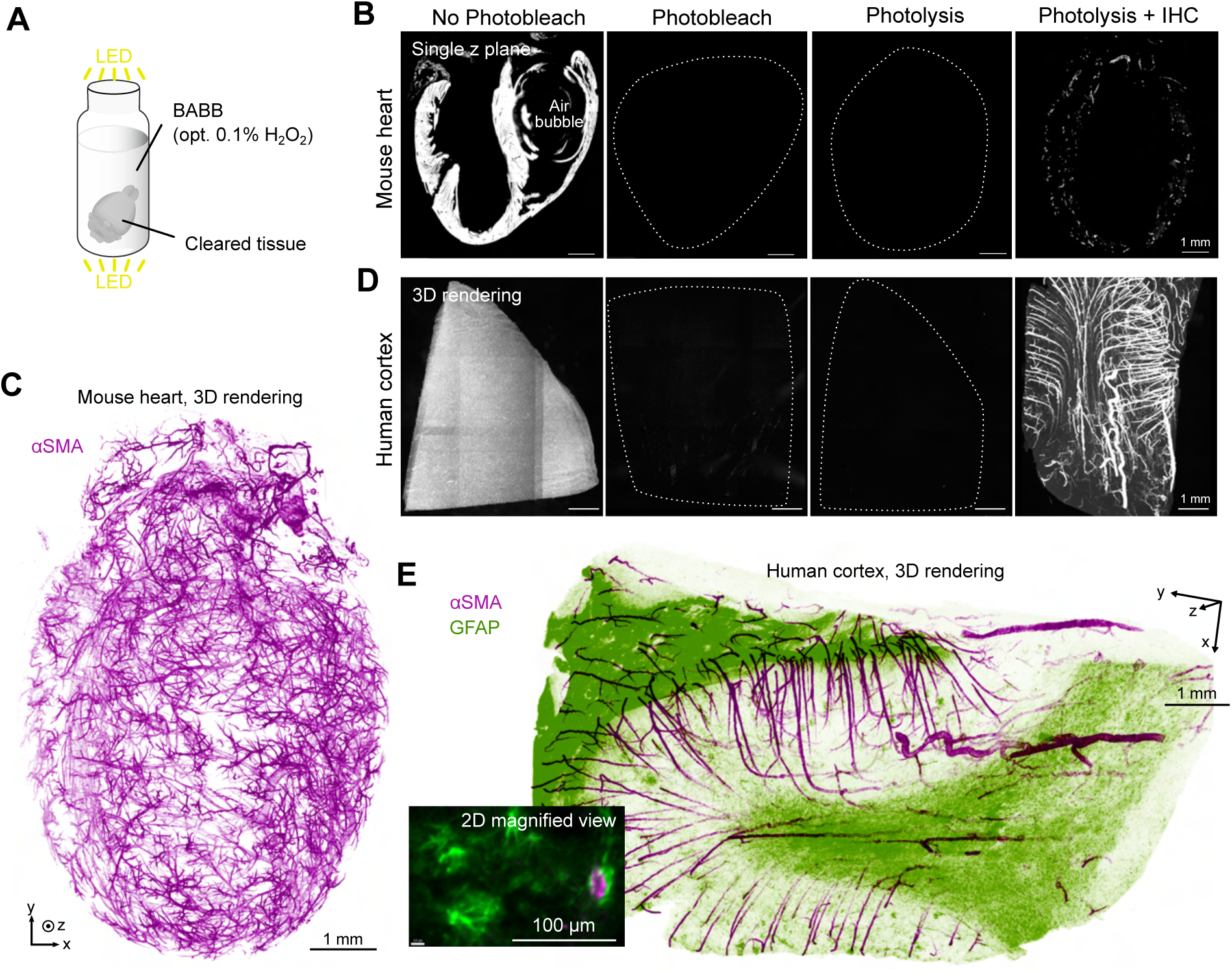
Application of the photobleacher to the pigment-rich volumetric tissues after clearing. (**A**) Schematic illustration of the photobleaching process. The tissues were illuminated after clearing, with hydrogen peroxide included when necessary. (**B**) Mouse hearts with various photobleaching conditions. Representative single z-planes are shown. (**C**) Three-dimensional rendering of a mouse heart after IHC staining using an anti-αSMA antibody. (**D**) Pieces of human cortex from an identical donor, with various photobleaching conditions. Three-dimensional renderings are shown. Outlines of tissues were shown in dot lines when necessary. (**E**) Three-dimensional rendering of a human cerebral cortex after IHC staining using anti-αSMA and anti-GFAP antibodies.

## DISCUSSION

In this study, we propose an open-source photobleacher designed to assist researchers in performing fluorescent imaging of pigment-rich tissues. Along with our simulator, this tool allows users to freely explore and redesign the photobleacher based on their scientific objectives and the scale of their experiments. The project is available under an open-source license, enabling researchers to modify and distribute it as needed.

Our simulator is reliable for predicting illuminance patterns (**Fig. 1F**), but the interpretation of its absolute values should be made with caution, as it is not intended to predict the actual luminance of the light sources. The luminance of LEDs is a function of both temperature and current, and the theoretical maximum luminance is typically not achievable due to various factors that affect both. While heating is a major factor that reduces the efficiency and brightness of LEDs, we chose not to complicate the cooling system in order to keep the design as user-friendly as possible. If users wish to increase brightness by replacing our LEDs with ones that have higher luminance or to increase the density of the LEDs, it is important to consider additional cooling systems, such as air fins, liquid coolants, or placing the device in a cold room (Duong and Han, 2013).

Expected use cases for the photobleacher include, but are not limited to, human tissue pathology, heme-rich tissue histology, and plant histology. To help users get started, we have provided a general guide on how to choose the appropriate photobleaching conditions (**Fig. 6**). For imaging peptides, photobleaching in PBS is the first choice. For imaging RNAs, we do not recommend using PBS, as we have observed a degradation in imaging quality, likely due to RNA hydrolysis. Instead, we advise performing photobleaching in a water-free environment, such as alcohol, air, or an organic clearing solvent, to prevent unwanted signal loss. If residual autofluorescence interferes with imaging of the signals of interest, H2O2 can be added during photobleaching. The use of H2O2 for de-pigmentation, particularly to remove melanin, has been reported in previous studies (Pigoli et al., 2019). We found that 0.1% is sufficient to eliminate autofluorescence without significantly reducing signal strength. Please note that the required concentration of H2O2 varies depending on the amount and type of pigment present. It is important to determine the lowest effective concentration of H2O2 that removes autofluorescence while minimizing potential damage to target molecules from radical reactions during photolysis. Additionally, it is crucial to follow the safety instructions in this manuscript to prevent possible accidents, as mixing hydrogen peroxide and benzyl benzoate can form an explosive chemical (see **Safety Manual**).

**Figure 6.**
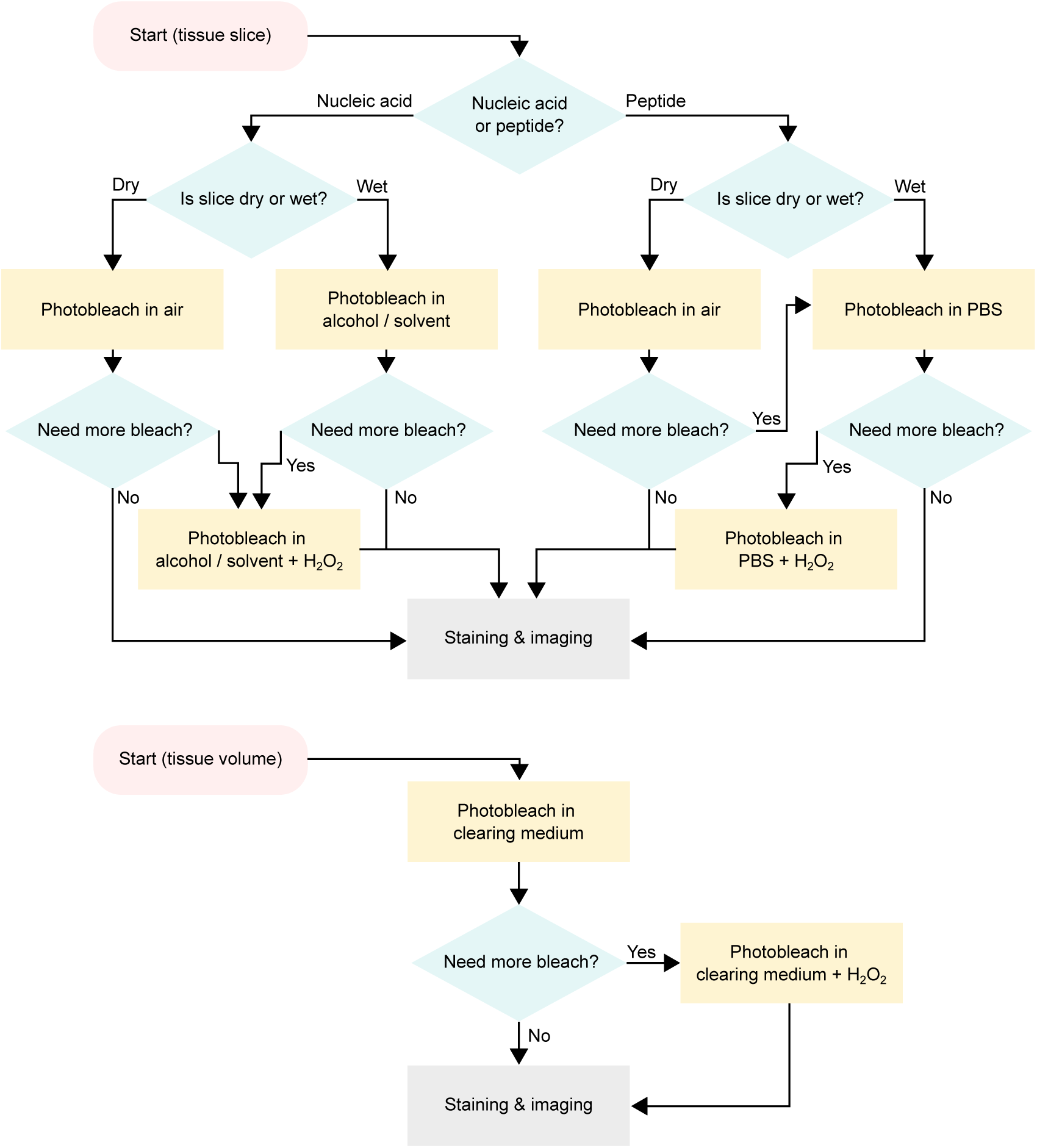
Flowchart guide for photobeaching. Flowcharts illustrating the choice of immersion medium for both two-dimensional tissue slices (top) and volumetric tissues (bottom).

For volumetric tissue samples, we used BABB as a clearing solvent in this study. One limitation of organic solvent-based clearing methods, such as BABB, is that the alcohol dehydration process can reduce peptide antigenicity, leading to the loss of signals for some peptides. The compatibility of alcohol dehydration with immunostaining has been previously discussed (Renier et al., 2014). Although we have not explored this in detail, aqueous clearing solutions, such as ELAST and CUBIC, offer promising alternatives to organic solvents, providing better antigen retention and potentially expanding the repertoire of applicable antibodies (Susaki et al., 2020; Park et al., 2024).

It is important for users to be aware that residual autofluorescence can interfere with the interpretation of images. The photobleacher is designed to reduce autofluorescence, but it does not guarantee complete eradication. To date, no reported photobleacher or chemical quencher has comprehensively demonstrated the full eradication of autofluorescence. Therefore, validation of signal specificity is crucial to ensure that signals originate only from the targeted molecules. In our study, we intentionally left the 488-nm excitation channel unused for mRNA/peptide staining to confirm that signals in other visible channels did not arise from autofluorescence. If autofluorescence-derived signals were present in other channels, we would expect to observe corresponding patterns in the 488-nm excitation channel. The absence of such a pattern confirms that the signals solely originate from the targeted staining.

While leaving one excitation channel unused for autofluorescence is the recommended practice, using all excitation channels for targeted staining may occasionally be necessary to expand the capacity for imaging multiple molecules simultaneously. In this case, we strongly recommend using short-wavelength excitation for the most abundant molecules, along with a blank control that lacks staining in the short-wavelength channel.

Given the customizability and scalability of our photobleacher, we believe it will accelerate 3D histology for large biological specimens, such as large human brain samples. By providing an open-source, adaptable platform, this tool has the potential to advance fluorescence-based imaging techniques across a wide range of applications, from basic research to clinical diagnostics, fostering greater accessibility in the study of complex biological systems.

## MATERIALS AND METHODS

### Model Systems and Permissions

#### Mice

All animal procedures were approved by The Rockefeller University Institutional Animal Care and Use Committee (IACUC) and complied with the guidelines set by the National Institutes of Health. Mice were always allowed ad libitum access to food and water, weaned at three weeks of age, and maintained on 12-hour light/dark cycle.

#### Human Tissue

Postmortem human tissue samples used for this study were obtained either from Miami’s Brain Endowment Bank or Science Care, Inc. The tissues were collected and banked in a fresh frozen state or fixed and embedded in paraffin. The tissues donors were de-identified before receipt by The Rockefeller University.

#### Simulation

The detail of the simulation is stated in the main text of this manuscript. The Python code used for the simulation is available at GitHub (https://github.com/tatz-murakami/photobleacher). To calculate the peak luminous intensity, *I*_0_, from the luminous flux, Φ, which is usually shown on the datasheet of the manufacturer, we used the following conversion formula under the assumption of the Lambertian distribution.

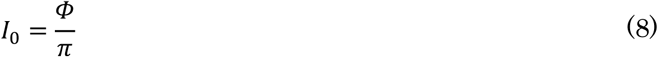

#### Measurement of illuminance

To verify the simulation, we assembled a photobleacher with downscaled luminance. We initially attempted to measure the illuminance of our open-source photobleacher, but were unable to do so because the illuminance exceeded the measurable range of a commercially available light meter. For the downscaled luminance, we used LEDs with 20 lumens (RA-IP20-80CRI-5m), arranged with 17 mm and 15.5 mm intervals along the x and y axes, respectively. The light meter (Dr.Meter, LX1330B) was placed 45 mm away from the array, and the illuminance was measured at intervals of 2.92 mm. 54 measurement points were recorded (7 along the x-axis and 9 along the y-axis). The measured values were converted to a heatmap using linear interpolation, as shown in **Fig. 1F**.

#### Assembly of photobleacher

The latest version of the assembly instruction is available on GitHub (https://github.com/NBelenko/Photobleacher).

#### 2D immunostaining

An FFPE tissue block was obtained from Miami’s Brain Endowment Bank. A hippocampal piece was collected from an 83-year-old male donor with Braak stage III - IV of Alzheimer’s disease. Five-micrometer-thick tissue slices were prepared using a microtome (Leica Biosystems) and placed onto slides. The tissue slices underwent photobleaching for 3 days in air. To deparaffinize the tissue, slices were sequentially treated with xylene twice, 100% ethanol twice, 95% ethanol, 75% ethanol, and PBS for 5 minutes each. Antigen retrieval was performed in 1× citrate buffer at 99°C for 10 minutes, followed by cooling to room temperature and washing twice with 1× PBS. The tissue slices were blocked with 5% normal donkey serum and 0.5% Triton X-100 in 1× PBS for 1 hour at room temperature. Primary antibodies, anti-phospho-tau (AT8, Cat# MN1020) and anti-NeuN, were diluted 1:100 in a buffer containing 3% normal donkey serum and 0.3% Tween 20 in 1× PBS, and the slides were incubated at 4°C overnight. After washing five times with 1× PBS for 5 minutes each, the secondary antibodies—donkey-derived anti-chicken antibody conjugated with Alexa Fluor 568 and donkey-derived anti-mouse antibody conjugated with Alexa Fluor 647—were added at a 1:500 dilution in a buffer containing 3% normal donkey serum and 0.3% Tween 20 in 1× PBS. The slides were incubated at room temperature for 2 hours, then washed three times with 1× PBS for 5 minutes each. If necessary, tissues were treated with 1:40 diluted TrueBlack Plus in 1× PBS for 5 minutes at room temperature. After washing with 1× PBS for 5 minutes three times, the slides were treated with 2 µg/ml DAPI (1:2500) in 1× PBS for 10 minutes at room temperature. Finally, the tissue slices were washed again with 1× PBS for 5 minutes three times and mounted with ProLong Gold antifade mountant under a cover slip.

### 2D fluorescent in situ hybridization

A fresh frozen tissue block was obtained from Miami’s Brain Endowment Bank. A motor area of the cerebral cortex was collected from a 63-year-old male donor with no known history of neuropsychiatric or neurological conditions. Fourteen- micrometer tissue slices were prepared using a cryostat (Leica Biosystems) after embedding the tissue block in O.C.T. compound, and then placed onto slides. The tissue slices were fixed with 4% PFA for 15 minutes at 4°C and dehydrated with ethanol according to the RNAscope instructions. Photobleaching was performed for 3 days in ethanol immediately following dehydration. The rehydration and staining processes were carried out according to the RNAscope instructions. We stained mRNAs in human brain slices using the RNAscope Multiplex Fluorescent v2 assay.

### 3D immunostaining

#### Mouse heart

C57BL/6J mice obtained from Jackson Laboratory were used in this study. The animals were euthanized with an overdose of pentobarbital (>100 mg/kg), followed by transcardial perfusion with 15 ml of ice-cold PBS and 20 ml of 4% paraformaldehyde (PFA, Electron Microscopy Sciences) dissolved in PBS. The hearts were carefully removed and post-fixed in 4% PFA-PBS at 4°C overnight. The photobleaching was performed as described below in the human brain section. Immunostaining was performed using a standard iDISCO protocol with modifications to the clearing step. For antibody staining, 1:200 diluted anti-αSMA and anti-GFAP antibodies were used. The staining process took 10–14 days. After dehydration with methanol (MeOH), the samples were cleared using BABB.

#### Human brain

A fresh frozen tissue block was obtained from Science Care from a 65-year-old female donor with no known history of neuropsychiatric or neurological conditions. The fresh frozen tissue was placed at -20°C for 3 hours prior to PFA fixation to allow mild defrosting. The tissue was then transferred to 4% PFA in 1× PBS at 4°C and gently shaken overnight. The tissue was washed with 1× PBS for 2 hours, with gentle shaking twice. Dehydration was performed using 50%, 80%, and 100% MeOH, with a 1-hour incubation for each step. The 100% MeOH solution was refreshed and the tissue was gently shaken for three nights at 37°C. The solution was replaced with a 1:2 mixture of benzyl alcohol and benzyl benzoate (BABB), and the tissue was shaken in this solvent for 2 hours. If necessary, 0.1% H2O2 was added to the BABB solution. For the detail on the use of H2O2 in the clearing solvent, carefully read the **Safety Manual** and follow the protocol outlined in the manual. This process was repeated until the tissues became transparent. The cleared tissues were placed in glass vials filled with BABB (+0.1% H2O2) and subjected to photobleaching. We photobleached the samples for three days without H2O2 and for one day with H2O2. After photobleaching, the samples were immersed in 100% MeOH and gently shaken for 2 hours at room temperature, with MeOH refreshed three times. Immunostaining was performed using a standard iDISCO protocol with modifications to the clearing step. For antibody staining, 1:200 diluted anti-αSMA and anti-GFAP antibodies were used. The staining process took 14 days at 37°C with gentle shaking. After dehydration with MeOH, the samples were cleared using BABB.

### Microscopy

#### Confocal Microscopy

We used the LSM700 (Zeiss) equipped with lasers operating at wavelengths of 405, 488, 555, and 649 nm. Imaging was performed using either a 10× Plan-Apochromat (0.45 NA, WD = 2.0 mm, Zeiss) or a 20× Plan-Apochromat (0.8 NA, WD = 0.55 mm, M27, Zeiss) detection objective. The scanning fields were covered by horizontally tiling multiple fields of view, with tiles stitched using 10% overlaps. Image acquisition was controlled via Zen Black software.

#### Light-sheet Fluorescence Microscopy (LSFM)

Imaging of cleared tissues was conducted using the Cleared Tissue LightSheet (CTLS) system from Intelligent Imaging Innovations. This system includes fiber- coupled lasers operating at 405, 488, 561, 640, and 785 nm wavelengths. A spatial light modulator shapes the beam, and the light sheet is generated by a galvanometric mirror. Illumination objectives (5×/0.14NA) are positioned on both the left and right sides of the imaging chamber, which is filled with BABB (refractive index = 1.56). Image acquisition is achieved through a Zeiss PlanNeoFluar Z 1×/0.25NA objective with a working distance of 56 mm. The ORCA-fusion BT serves as the sCMOS detector.

The illumination wavelength was alternated for each z-stack while chromatic focus shifts were corrected by automatically moving the detection lens. The motorized stage moved the sample in the XY plane to capture the entire tissue volume. Illumination focus was swept at three positions in a single z-plane. Optical zoom was applied at 5×, yielding pixel sizes of 1.3 μm. The z-step interval was set at 2 μm. All system components were managed by Slidebook software (Intelligent Imaging Innovations), and the data was saved on a Dell PowerEdge R740XD server with twelve 16 TB HDDs with RAID configuration, connected via 10-GB Ethernet fiber.

### Image Analysis

#### Browsing, overlaying, cropping, and linear intensity adjustment

We used different software for image adjustment depending on the data size and dimensions of the image. For 2D images smaller than 4 GB, we used FIJI. For 2D images larger than 4 GB, we used QuPath. For 3D images obtained from light-sheet microscopy, we used Napari with Python coding.

#### Fluorescent Decay Curve

The images were registered using the registration plugin in FIJI. Images from multiple time points were stacked along the time dimension, and descriptor-based series registration was performed. We randomly selected peaks of the lipofuscin autofluorescence for each tissue slice and measured the transitions of the averaged intensities over time.

### Statistical Analysis

Statistical analyses were performed using SciPy with Python 3.9. A p-value of < 0.05 was considered as significant. Welch’s t-test was used when multiple conditions were grouped into a single condition, while Student’s t-test was applied otherwise.

### Key Resources Table

| REAGENT or RESOURCE | SOURCE | IDENTIFIER |
| --- | --- | --- |
| Antibodies |  |  |
| anti-GFAP, Cy3 conjugated | Sigma-Aldrich | C9205 |
| anti-αSMA, Alexa Fluor 647 conjugated | Novus Biologicals | IC1420R-100UG |
| anti-NeuN | Sigma-Aldrich | ABN91 |
| anti-Phospho-Tau (AT8) | Invitrogen | MN1020 |
| Donkey anti-chicken Alexa Fluor 568 | Thermo Fisher Scientific | A78950 |
| Donkey anti-mouse Alexa Fluor 647 | Thermo Fisher Scientific | A32787 |
| Dyes |  |  |
| DAPI | Sigma-Aldrich | D9642 |
| RNAscope |  |  |
| Pcp4 | ACD | 446111 |
| Htr2c | ACD | 450651-C2 |
| RNAscope Multiplex Fluorescent Reagent Kit v2 | ACD | 323270 |
| HybEZ oven | ACD | 321710 |
| Humidity control tray | ACD | 310012 |
| EZ-Batch Wash Tray | ACD | 321717 |
| EZ-Batch Slide Holder | ACD | 321716 |
| Chemicals and reagents |  |  |
| Pentobarbital |  |  |
| Paraformaldehyde | Electron microscopy sciences | 15714 |
| PBS (10×) | Sigma-Aldrich | P3813 |
| Ethanol | Sigma-Aldrich | E7023 |
| Methanol | Fisher Scientific | A412SK-4 |
| Hydrogen peroxide (for 2D staining) | Thermo Scientific | H325 |
| Hydrogen peroxide (for photobleaching) | Sigma-Aldrich | 516813 |
| Benzyl alcohol | Sigma-Aldrich | 24122 |
| Benzyl benzoate | Sigma-Aldrich | W213802 |
| Donkey Serum | Jackson ImmunoResearch | 017-000-121 |
| Xylene | Fisher Scientific | HC-700 |
| TrueBlack Plus | Biotium | 23014 |

## ACKNOWLEDGEMENTS

This work was supported by Japanese Society for Promotion of Science Overseas Research Fellowship (to T.C.M), and Leon Levy Scholarships in Neuroscience (to T.C.M.). Jie Xing assisted with the perfusion of a mouse. Human tissues were obtained from either the University of Miami’s Brain Endowment Bank or Science Care, Inc.

## AUTHOR CONTRIBUTIONS

T.C.M. supervised and designed the project, and performed simulations and experiments. N.B. designed and assembled the prototype and N.B. and G.D. designed and assembled the photobleacher. N.H. brought the grant and supervised the project. C.W., E.S., and Y.H. performed experiments using biological tissues. C.P. supervised and designed the FISH experiment.

## DATA AND SOFTWARE AVAILABILITY

The Python code used for the simulation is available at https://github.com/tatz-murakami/photobleacher. The latest version of the assembly instructions and all files needed to replicate the photobleacher are available at https://github.com/NBelenko/Photobleacher. Due to the large size of the imaging datasets generated in the study, these datasets are not available in a public repository but can be obtained from the authors upon request.

---

**Table S1.** List of published photobleachers using array LEDs. List does not include photobleachers with a single LED. *The estimated illumination range was calculated by multiplying the interval by the number of LEDs. This calculation is based on our simulation, which predicts that the illumination outside of this range drops sharply. †The number is an inferred value based on information provided by the article or the product description.

| Authors<br>(Published<br>year) | Device | Distance<br>between<br>sample<br>and LED | Interval | Number<br>of LEDs | Power | Estimated<br>illum.<br>range | Color | Illuminat<br>ion<br>direction |
| --- | --- | --- | --- | --- | --- | --- | --- | --- |
| Duong and<br>Han (2013) | GrowLight,<br>HTG Supply<br>Inc., Orange,<br>CT (Product<br>discontinued) | 0.7 cm | not<br>specified | 80 | 3 W /<br>LED (240<br>lm /<br>LED†) | 25 cm x<br>15 cm* | Mlx of<br>390, 430,<br>460, 630,<br>660 and<br>850 nm | Single |
| Sun and<br>Chakrabartty<br>(2016) | DBPOWER<br>Eye-<br>protection<br>LED Desk<br>(Product<br>discontinued) | ~ 1.0<br>cm† | ~ 1 cm† | 2 x 11 | 27.3 lm<br>/LED | 2 cm x 11<br>cm* | White,<br>~6000<br>K† | Single |
| Pigoli et al.<br>(2019) | 300 W<br>Viugreum™<br>spotlight<br>(Yilangde<br>Illumination<br>CO. Ltd.,<br>China) | 8.8 cm | 1 - 1.2<br>cm† | 108 | 0.455<br>mW/cm²<br>(92.6 lm /<br>LED) | 24 cm x<br>12 cm | White,<br>6500K | Single |
| Ku et al.<br>(2020) | LEDMO<br>SMD5630<br>LED Strip on<br>a regular<br>glass beaker | less than<br>5 cm† | ~1.7<br>cm† | Not<br>specified | 25 lm /<br>LED | Not<br>specified | White,<br>3000 K | Circled |
| Tsuneoka et<br>al. (2022) | TiYo, LED<br>from<br>NV9W149AM,<br>Nichia<br>Corporation,<br>Japan | 0.5 cm | 4 cm | 2 x 6 | 4,131 lm<br>at a<br>3,000-mA drive<br>current /<br>LED | 8 cm x 24<br>cm | White,<br>5000 K† | Dual with<br>mirror |
| Park et al.<br>(2024) | Custom-made | Not<br>specified | Not<br>specified | Not<br>specified | Not<br>specified | Larger<br>than 8 cm<br>x 8 cm† | White | Not<br>specified,<br>single or<br>dual |

**Fig. S1.**
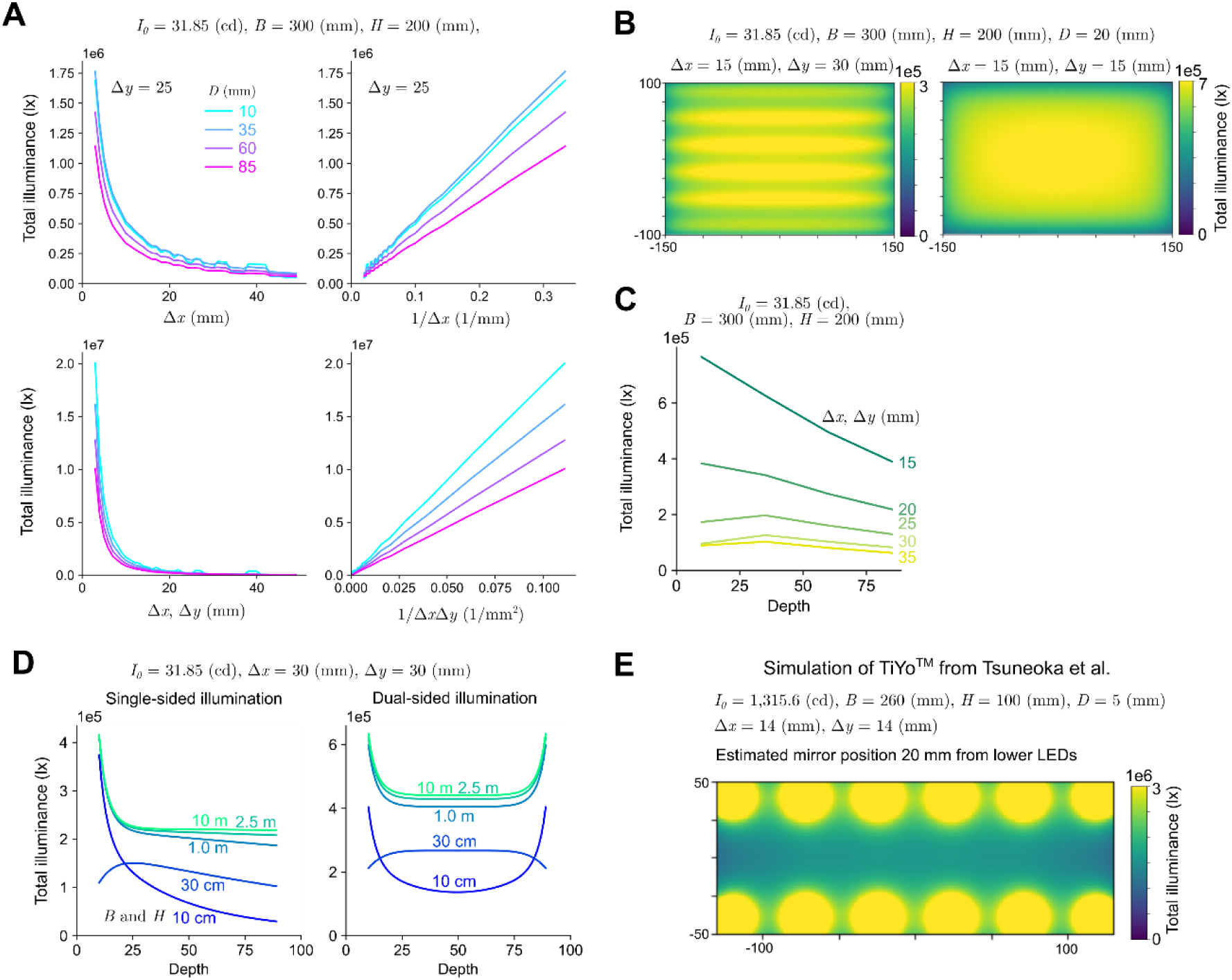
Simulation of total illuminance from multiple light sources. (**A**) Inverse proportional relationship between interval length and total illuminance. (**B**) Heatmap representations of the simulation of the total illuminance for non-isotropic (left) and isotropic (right) interval length. (**C**) Decay of illuminance with increasing depth. (**D**) Effect of board size on illuminance decay with a 30 mm isotropic interval. Left: single-sided illumination; Right: dual-sided illumination. (**E**) Heatmap representation of the simulation of total illuminance based on the design from Tsuneoka et al.

**Fig. S2.**
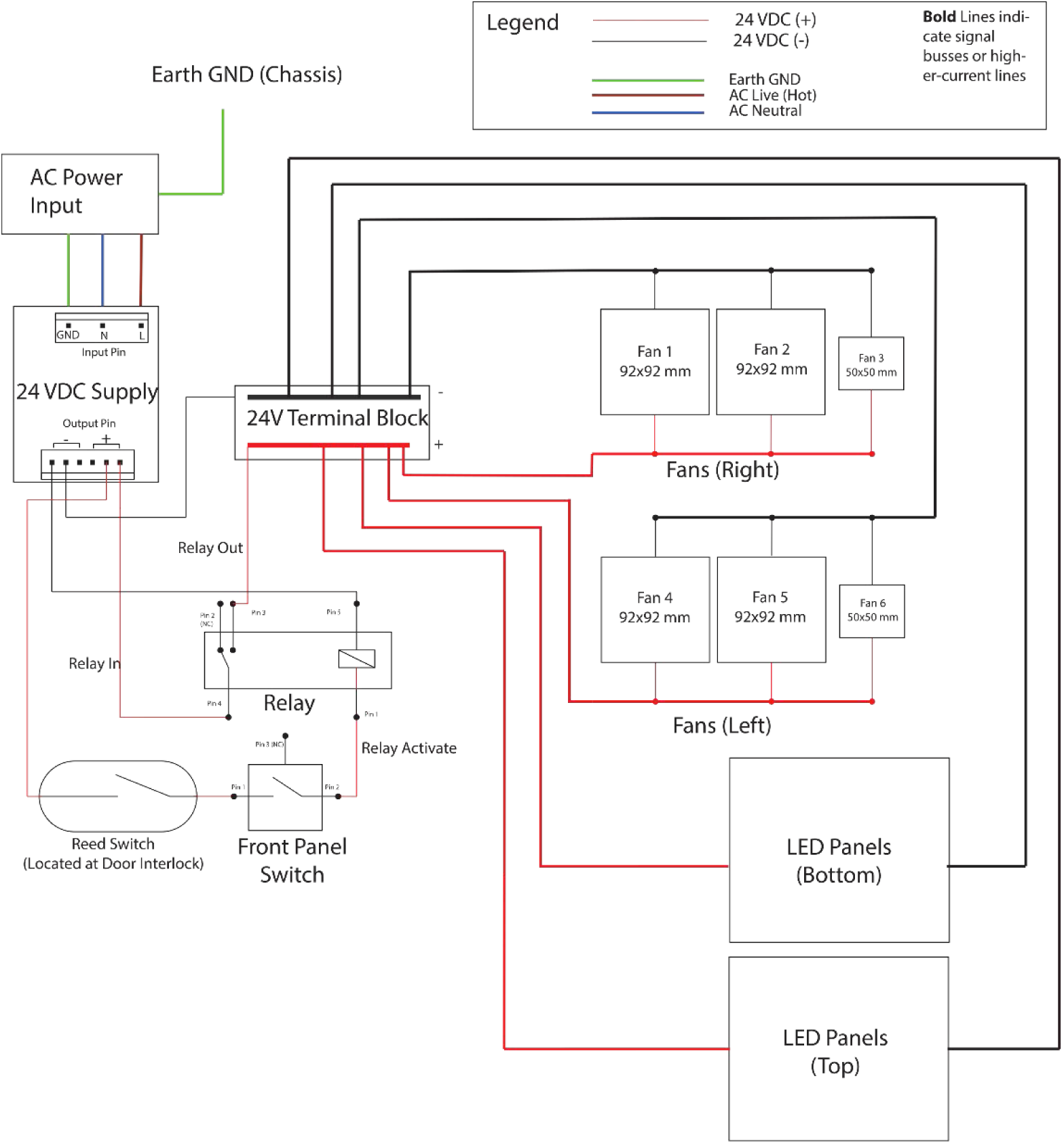
Wiring diagram of the open-source photobleacher.

**Fig. S3.**
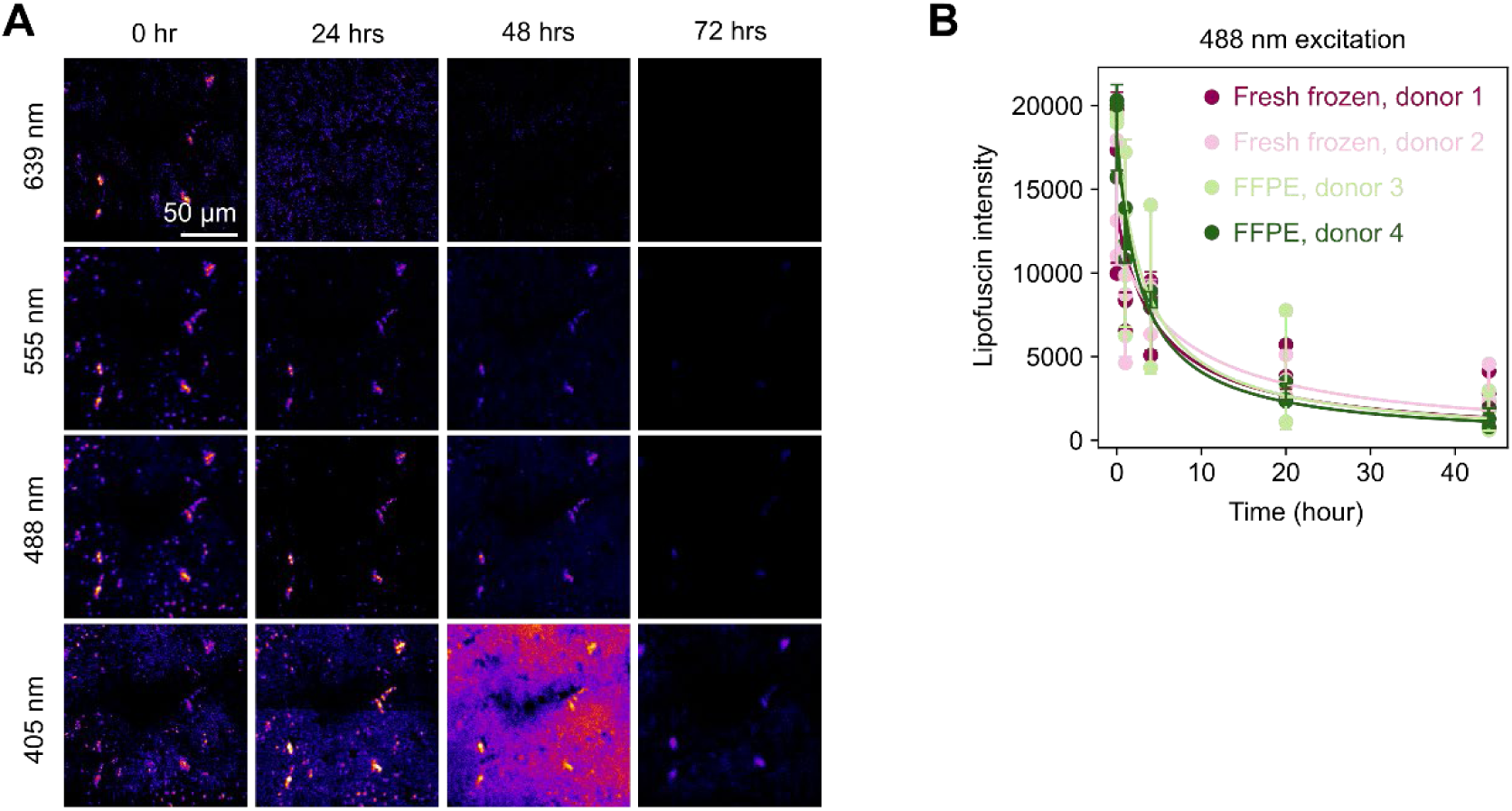
Time series observation of the effect of the photobleacher on human cortical slices. (**A**) Time-series observation of a dried human tissue slice during photobleaching. (**B**) Quantification of lipofuscin intensity during photobleaching. Two FFPE tissue slices and two dried-fresh frozen tissue slices were obtained from independent donors.

## Supporting text

### Supplementary Note for Photobleacher Device

Construction of Device:

The Photobleacher consists of a stainless steel enclosure with LED panels along the floor and ceiling. Samples are placed on a borosilicate glass shelf, located approximately in the middle of the box. 6 High powered fans are used for cooling. The device is powered with a 24VDC 200W power supply.

Electrical Design Procedure:

We’ve designed this Photobleacher to have a minimum total luminous flux *Lmt* of about 9600 lumens. For choice consideration in our LEDs, we wanted ones that were powerful but still energy efficient, with our desired colored temperature of 3000k. We chose the <u>MP-3030</u> model LED from Luminus Devices Inc. Each LED has a luminous flux of *LMled* = 38lm, where *LMled* is the luminous flux of each LED. A single a continuous current of 65mA and a forward voltage of 2.68V. They are SMD mounted and must be placed onto a PCB substrate.

To determine the minimum number of LEDs needed, we divided *Lmt* by *LMled*. The result was

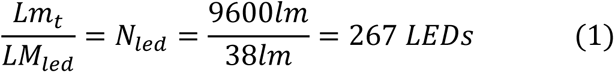

Where *Nled* is the total number of LEDs. This comes to 133 LEDs per side. Since each LED has an area *AL* of 3mm x 3mm = 9mm^2^, the total minimum area *Amin* comes is calculated by

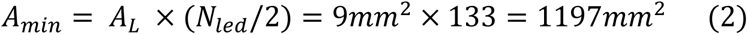

Duration required for bleaching is inversely related to the total luminous flux, which influenced two design decisions. First, we wanted to have uniform illumination on the shelf holding the samples to avoid having the samples bleach at different times at different parts on the shelf. Second, we wanted to include the ability to add more LEDs, and thus higher illumination, to the system with ease.

To that end, we designed a modular PCB with hermaphroditic connectors on all 4 sides. This provides the ability to easily connect multiple PCB’s together in a grid configuration. The boards can be connected in any orientation. Since these LEDs have a beam angle of 120(deg) an additive effect can be achieved where the uncollimated light from adjacent boards will pass through the same sample volume. To achieve this, and to keep things constrained when it comes to size, we designed a PCB around a 100mm pitch tiling pattern. The PCB’s can be connected together in a grid pattern. We chose a grid size of 3 x 2, or 6 PCB panels maximum for each top and bottom (12x total). This is elaborated on further in the enclosure design section.

In order to calculate the power needed, we determined our desired voltage and current draw for each board. Since 24V power supplies are common and relatively safe, we wanted each board to be fully powered at 24V. Since the forward voltage of each LED is 2.87V, the total number of LEDs was calculated to be

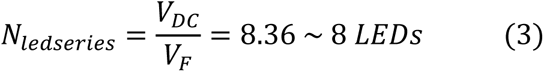

where *Nledseries* is the number of LEDs in series, *VDC* is the total DC voltage, and *VF* is the forward voltage of each LED.

To increase the luminous flux, each board has 8 parallel columns of 8 LEDs in series, for a total of 64 LEDs per board. For 6 boards, this comes to a total of 384 LEDs per side, or 768 LEDs total. To limit the current and avoid damage to the LEDs, a 39 Ohm resistor was placed in series in each column.

For calculating the power drawn from each board, we must find the required voltage and current to drive the LEDs to full brightness. For a series circuit, the voltage and current is calculated by

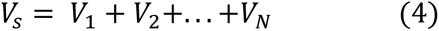

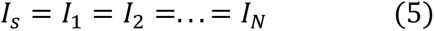

where *Vs* and *Is* are the total voltage drop and current draw, respectively, across all components in series. *VN* and *IN* are the voltage drop and current draw for each component. For parallel circuits, the voltage and current are calculated by

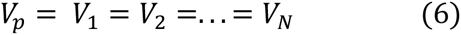

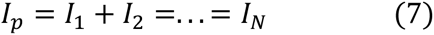

Where *Vp* and *Ip* are the total voltage drop and current draw, respectively, across all components in parallel.

The current draw of individual LEDs *Iled* is 65mA, while the forward voltage of individual LEDs *VF* is 2.87V. Therefore for each column of 8 LEDs in series, the voltage and current are calculated using equations (4) and (5), with the results being *Vs* =22.96V and *Is* = *Iled* = 65mA.

Since 8 series circuits are connected parallel to each other, by (6) *Vp* = *Vs*, and by (7) *Ip* = *Iled* * 8 = 0.52A. Therefore the board voltage *Vboard* = *Vs = Vp =* 22.96V and board current *Iboard*=0.52A. With our design procedure, we can take *Vboard* = 24V.

Since each board can be considered as parallel to each other, the required voltage stays at 24V, while the current is additive. The total current draw *It* for 12 LED panels is therefore

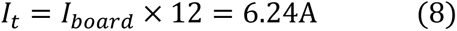

Power, in watts, is calculated by the formula

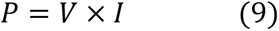

and for each board having power *Pboard*,

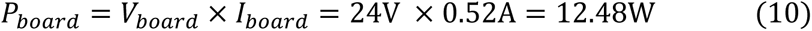

Therefore, the total power needed for all 12 LED panels is *PL* = 12 x (*Pboard*), where *PL* is the total power from the LEDs, is

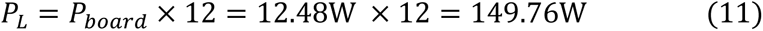

To provide cooling, we chose fans that were efficient yet had a strong airflow. Two sets of fans were used, one set to provide air intake and another for air outflow. Each set of fans contains 2 model AFB0924HD fans and one model AFB0524VHD fan, both from Delta electronics. Each fan draws 2.4W of power according to their respective datasheets. For 6 fans total, the power drawn by the fans, *Pfans*, is

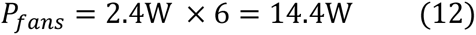

Therefore, the total power *PT* needed for the device is

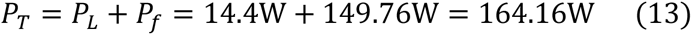

We therefore chose <u>this</u> 24V 200W power supply from TDK-Lambda. This provides plenty of power for our maximum requirement and allows some room to add even more LED panels if desired.

Design of the enclosure for the photobleacher:

In the design of the enclosure of the photobleacher, we needed to satisfy a few general requirements. The first was that of thermal management. The brain tissue samples cannot exceed an ambient temperature of 38C or so without sustaining damage. Ideally, we want to keep the ambient temperature as close to room temperature as possible. While the LEDs employed in this photobleacher are very efficient, their large number and close spacing mean that there is a large possible temperature rise if the chamber is left alone and unventilated for long periods of time.

Given the constraints of the 100mm-pitch, tiled PCB design (see electrical design section) we opted for a 2x3 tile configuration for both the top and bottom lighting panels (12x total boards) this 200mmx300mm area was chosen based on the physical space constraints in the laboratory space where the photobleacher was to be installed as well as to keep the power requirements to a reasonable level.

Theoretically one can extend the tile pattern so long as the amperage does not exceed the respective ratings of the connectors.

The other major consideration in the enclosure design is the height of the chamber. We want to maximize luminous flux so we tried to find the smallest height possible based on the existing containment vessels used for samples in the laboratory. Measuring the height of the containment vessels, we chose a chamber height of approximately 70mm.

We determined that an air cooling system for the enclosure would be sufficient based on our experience with the first iteration of the photobleacher. We designed the enclosure around two 92mm fans running on each side of the chamber (each blowing in the same direction to maximize airflow) and two 50mm fans running on each side of the power supply. The 92mm fans were chosen to provide airflow over the full height of the sample chamber as well as across the LED boards.

The initial CAD for the enclosure was done in Autodesk Inventor using the built-in sheet metal design tools. From this design we were able to create a projected model for a laser-cut prototype of the enclosure to test thermals with. After cutting and assembling a prototype of the box out of black acrylic, we set up an IR temperature probe in the sample chamber, aimed at a target made from black gaffer’s tape. We left the photobleacher running for extended periods to see where the temperature leveled off. After numerous tests, including one continuous run lasting over 4 hours, we found that the measured temperature never exceeded 32C. This was more than satisfactory and well within the defined thermal limits.

After confirming the thermal design, we sent drawings and models of the enclosure in its sheet metal form to Protocase for production. After finalizing the design with them we put in an order for the case in white powdercoated steel sheet, complete with a door and latch.

There were some modifications that needed to be made to the case once it was received. Initially there was no interlock on the door of the enclosure to ensure that the lights could not operate with the door open. We wired in the interlock in the form of a magnetic proximity switch and a neodymium magnet glued in place on the door itself. The opening for the power inlet module also had to be modified as we had not accounted for the thickness of the powdercoat when specifying the opening dimensions.

In addition to the modifications to the enclosure itself, we made two modifications to the glass stage to protect the electronics below in case of a spill. First, a 3D printed barrier made of white PETG was glued in place around the stage with white silicone sealant (Dowsil 732). Captive sealing washers were also added to the mounting points to prevent liquid ingress.

## Supplementary Materials: Photobleacher Assembly Instructions

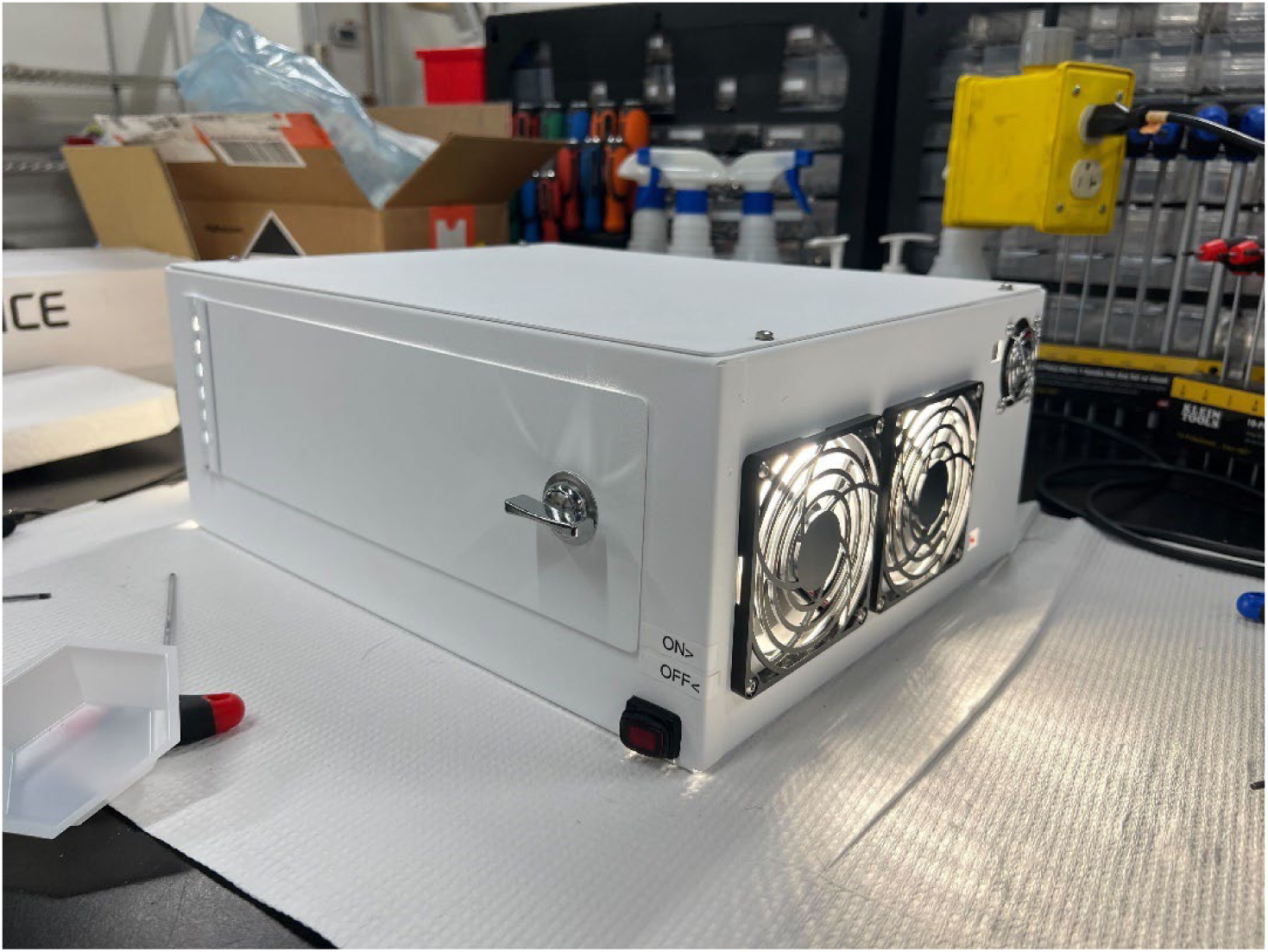

### I - SAFETY CONSIDERATIONS

Before beginning the assembly, please take the following safety precautions.

1. **Make sure the device is unplugged and switched off before performing any maintenance or connection checks.** The device is designed to operate at a constant +24VDC, which is a safe level for humans. However, please use caution when making connections with the power input module and any AC voltage. Any short could potentially cause damage to the components.
2. **DO NOT look directly at the LEDs when they’re on**. The LEDs used in this are extremely bright. Use appropriate eyewear protection or use a physical shield to block the light when using and testing the system.
3. Maintain standard electronics and workshop safety standards. This includes wearing eye protection, an emergency plan, and any other safety rules your space requires.

### II - PRE-ASSEMBLY NOTES

#### Notes on Wiring and Cable Management

For all the electronic connections to be made, the wires of the components may need to be extended. This is especially true for the fans and the front panel on/off switch. We offer several solutions for this in these instructions depending on soldering experience and soldering iron access. If you have access to a soldering iron, but are inexperienced with soldering, we recommend looking up tutorials on “lineman splice”, a technique to connect 2 wires together. If assembling the LED PCBs yourself, a tutorial on how to solder through-hole and surface mount components may be helpful.

With the many electrical connections within the device enclosure, it’s very easy for the wires to become tangled and messy. We recommend using various forms of cable management items, such as zip-ties, adhesive backed zip-tie holders, and Velcro straps. We’ve also added gaps in the bulkhead panel separating the bleaching chamber from the power supply portion. These gaps will allow you to bundle up wires and feed them through, making the inside much cleaner. Proper cable management and wire bundling will make it easier to fix any issues that may arise.

Finally, we have color coded the wires in these instructions. We’ve followed standard industry practices and coded the wires as: Red for +DC voltage, Black for -DC voltage, Brown for Live (Hot) AC signal, Blue for Neutral AC signal, and Green for Ground. These color codes are helpful for following signal lines and debugging. Also note that the -DC voltage is connected to the protective earth/chassis ground (denoted as GND) of the AC power input and may be used interchangeably. Earth Ground is connected to the chassis via the mounting screw on the DC power supply.

#### Note on Bulkhead Panel Placement

The bulkhead panel, which is the metal sheet that separates the power supply and power input section from the LEDs and fans, is a tight fit within the enclosure. When assembling this for the first time, we found the panel was most easily installed before mounting the fans on the enclosure. **Make sure to install the bulkhead before the final mounting of the fans, otherwise you may not be able to get the bulkhead in place.** Specifically, the large fans in the middle, when mounted, will block the panel. If desired, you can mount all the other fans, then install and secure the panel. After the panel is installed, you can mount the middle fans. When installing the panel, you may need to rotate it diagonally for it to fit inside the enclosure.

Custom-made Materials:

| Material | File Names | Notes |
| --- | --- | --- |
| Chassis/Enclosure | HEI-2400 RevA Photobleacher Enclosure Main Assembly | We used the third-party vendor, Protocase (https://www.protocase.com). A white powder-coated steel sheet was selected. |
| LED PCB | 100mm_LED_Panel (KiCad Project File) | We used the third-party vendor, PCBWay (https://www.pcbway.com). |
| Glass Sample Stage | GlassShelf.pdf | A 1/4"-thick borosilicate glass was manually trimmed using a glass cutter and drill. See Chapter X for the details. |
| Spill Protection Bezel | Spill_Tray_bezel.stl | 3D printed with a FDM printer, Prusa XL, using PETG. If a large-format 3D printer is not available, find a third-party 3D printing service. |

### III - TOOLS LIST

For a detailed list of parts, please refer to the Bill of Materials (BOM) included in the Supplementary Materials.

In addition to the BOM, some tools are required to complete the assembly. Other tools are not necessary but can be very helpful in debugging. A table for each set of tools is shown below.

### IV – LED PCB Assembly

We recommend having your PCB vendor do the assembly of the LED PCBs. You will need to provide all part names and numbers, which can be found in the PCB BOM. If you’ve received the boards pre-assembled, you can skip to Part 2 of this section, Testing & Verification. We ordered our boards from PCBWay and assembled them ourselves.

If you decide to assemble the PCBs yourself, you will need both Hot-Air Rework and solder paste. We also recommend ordering a Solder Stencil along with the PCBs. If available, a reflow oven is ideal for attaching the surface mount components to the front (LED) side of the PCB.

#### Part 1 – PCB Assembly

1. Start with the side with the LED footprints. Use the solder stencil to align the holes in the stencil with the footprints. Use a putty knife to spread the solder paste onto the board and onto the footprints through the holes. Remove the stencil and you should see each footprint filled with solder paste.
2. Carefully place the LEDs on the footprints (marked D1-D64 on the board). The corner with the white triangle tab indicates the cathode of the LED and should be aligned with the bottom-left corner, next the “D” mark, and towards the resistor footprints. See Figure 2 for reference.
3. Place the resistors on the resistor footprints (marked R1-R7 on the board). Polarity for the resistors does not matter; just align the resistors with the footprint.
4. Once all components are placed, use either a reflow oven (if available), or a hot air rework (heat gun) set to ∼500-600F. You can also use a standard heat gun but be aware of the high air flow moving the components.
5. Turn the board over and apply paste to the footprints (either with or without a solder stencil).
6. Place the remaining resistor on the R8 footprint.
7. Place the fuse chip on the F1 footprint.
8. Place the capacitors on the footprints marked C1-C8. The capacitors used in this assembly are polarized; make sure the line on the backage is connected to the +24VDC line.
9. Place the inter-board connectors onto the footprints marked J1-J4. Make sure the pins are aligned with the tab and that the pegs on the connectors are through the corresponding holes on the board. You can omit inter-board connectors where no connections will be made.
10. Repeat step 4 for this side of the board. **Note**: You may want to use a lower heat setting when reflowing the inter-board connectors, as too high a heat can cause the plastic to melt and the connection pins to become mis-aligned.
11. Let the board cool down after reflowing, then solder the tab connectors to the through holes marked J5-J12. You will only need to solder one set of tab connectors, and only to adjacent through holes (for example, J6 and J8 can be soldered in, and the left rest blank). The tab connectors should be sticking out of the bottom of the board and faced towards the edge of the board. See Figure 3 for reference.

#### Part 2 – Testing & Verification

The board should now be ready for testing. Use the following steps to ensure your boards are operational.

1. Prepare 2 wires and crimp one end with quick tab connectors, and the other end bare.
2. Connect the wires to the tab connectors on the board as shown in Figure 4. Note which one is the +24VDC line and which one is the negative line (marked GND on the board).
3. Using a variable benchtop power supply, first turn on the supply and bring the voltage down to 0V. Connect the supply probes to the wires and double check that the polarity is correct. Then, slowly turn up the voltage and watch the current reading. **If the current spikes when the voltage is below 1V, there is likely a short.** The fuse mounted on the bottom of the board should protect the components from excessive current. Refer to the troubleshooting guide of this section.

If there are no current spikes, slowly increase the DC voltage to 24V and verify that all LEDs are on. They should turn on dimly around 15-17V and start increasing in brightness as you turn up the voltage. If you notice any LEDs not turning on, refer to the troubleshooting part below.

You can speed up the testing of the rest of the LED boards once you have 12 boards assembled. Connect them together with the inter-board connectors (as described in the LED panel Section later in this guide). Connect up to 6 boards together; any more and you may exceed the current limit on your power supply. You can then test the 6 LED panels at once and check for any LEDs that are off.

#### Part 3 - LED PCB Troubleshooting

A - If no LEDs turn on:

1. Check the fuse on the bottom of the board. Use a multimeter and probe each side of the fuse to verify the connection. If there is no connection, then a current spike likely destroyed it, or it is not soldered on properly. Remove the fuse and check the part of the board for damage. You may also want to probe between the +24VDC and GND lines to see if there is still a short. If there is, remove all the capacitors from the back and probe again. If there is still a short, remove the inter-board connectors and probe again. If you are still getting a short, it is likely that the board is damaged, and a new one must be assembled.
2. If there are no shorts and the fuse is still intact, check the LEDs at the top of each LED column (the bottom of the board is located where the resistors are). The LED may be misaligned or orientated incorrectly or may be not connected altogether. Make sure the cathode marker is in the right place (next to the “D” mark) and reflow the LED again. If your multimeter has a diode check setting, you can verify the LED is still working by probing the anode with the positive probe and the cathode with the negative probe. The LED should light up if properly connected. Test again with the benchtop power supply and verify.

B - If some columns of LEDs turn on while others do not:

1. The issue is likely an LED that is not connected to the pad or in the wrong orientation, which interrupts the path for the electrical current. With a multimeter, use the diode function to test each LED in the column. Using fine-tipped probes, touch the small, exposed pad on the anode of the top LED to the anode of the next LED. You should see the LED light up and a voltage appear on the multimeter. Repeat this process down the column until you see the LED that doesn’t light up when probed. Reflow the offending LED and continue the process down the column until all LEDs are verified.

C - If individual LEDs don’t turn on:

1. The issue is likely a short on the pad of the LED that’s off. This can be caused by too much solder on the pad or misalignment of the LED. Remove the LED and inspect the bottom of it. If there’s any solder bridging the anode and cathode terminals, reflow the LED and lightly scrape the excess solder away with a pair of tweezers (if you have extra LEDs, you can simply replace the LED with a new one). Next, check the LED footprint pad for extra solder and scrape away any that you see while reflowing. Test the pads with a multimeter to ensure there’s no short. Replace the LED and reflow with the hot-air rework. Repeat for any other LEDs that are off.

### V - Power Input Assembly

For this section, use the wiring diagram for reference. The wiring diagram is included in the supplementary materials section

1. Place the Fuse in the fuse drawer as shown in Figure 5. Insert the drawer into the power input module.
2. Place the Power Input Module into the square slot in the back so the ON/OFF switch and power plug socket is facing the outside shown in Figure 6. Note: the fit might be tight, you will need to squeeze the tabs on the power input module inward to properly install. You also may need to file down the opening with a metal file.
3. Using the mounting holes in the back, screw in the DC Power Supply. The Terminal Bus Block should be attached between the Power Supply and the Relay mounting holes with double-sided VHB tape. Refer to Figure 7 for placement references. Note: If desired, you can drill mounting holes in the enclosure to align with the mounting holes on the Terminal Bus Block.
4. Prepare 3 wires of 3 colors: Blue, Brown, and Green, of at least 18AWG. One end of each wire should be crimped with quick tab connections, while the other end should be crimped with SVH-21T-P1.1 crimps and placed into the mating connector (part # VHR- 5N) for the AC/DC power supply header. The wire colors are coded as such: Brown – Hot; Blue – Neutral; Green- Ground. Connect the quick-tab ends onto the Power Input Module (refer to the labels on the module for each connection). Then plug in the connector to the header on the AC/DC power supply. The connection is shown in Figure 8.
5. Prepare 2 wires: 1 red and 1 black (the red wire should be at least the length of the back of the chassis plus the side of the chassis; the black wire should be cut short). Solder the wires onto the 2 vertical tabs on the relay (labeled pins 1 & 5; the connections are non-polar; you can use either tab for either wire. You may want to unmount the relay to make the soldering easier). Crimp the other end of the black wire with a ring terminal and screw into the terminal block on the negative DC side. See Figure 9 for reference
6. Prepare 4 wires: 3 red and 1 black. One Red should be long, while the rest should be cut short. The long red, 1 short red, and 1 black should be crimped with SVH-41T-P1.1 crimps and placed into the mating connector for the DC output of the AC/DC power supply (part # VHR-6N). The other end of the black wire should be crimped with a ring terminal and screwed into the negative side of the terminal bus. The other end of the short red wire should be crimped with a quick connect and plugged into one of the horizontal tabs on the relay. The last short red wire should have one end crimped with a ring connector, and the other with a quick connect tab. Plug the quick connect tab end onto the other tab on the relay next to the previous tab connection, and screw in the ring connector side to the positive side of the terminal bus. See Figure 10 for reference.

### VI - Fan Assembly and Bulkhead Panel Installation

Part 1 - Wiring:

1. Each fan has 2 wires: a red one for the positive DC voltage and a black one for the negative DC voltage. Each one must be connected to the terminal block on the appropriate side.
2. There are several wiring options based on available materials and expertise:

a. Tie all the positive wires of the fans **on one side** together and tie all the negative wires together. You can solder them, extend them (either lineman splice or a connector) and crimp the other end so they only have 1 connection to the terminal block. This will reduce clutter and cable management but may make it more difficult to troubleshoot if a fan stops working.
b. Extend the wires of each fan by soldering longer wires, following the red/black color scheme. Crimp the ends of the wires with ring terminals and screw into the appropriate sides of the terminal block.
c. Use extension cable with a matching connector to the fan connector. This was not done in this assembly but can be done as an alternative. The other ends of the cable still must be crimped with ring connectors and screwed into the terminal block.
3. Once all the fans are connected to the terminal block, do any necessary cable management and be sure to align groups of wires with the slots at the bottom of the bulkhead.
4. Repeat the process for the fans on the other side.

#### Part 2 - Bulkhead Panel Installation

**Note: The Bulkhead Panel can be tricky to place. Placing it before mounting the fans seemed to be the easiest option during our assembly.

1. Before inserting the panel, align any wires/cables to the gaps at the bottom of the panel.
2. Orient the Bulkhead so that the tabs are facing towards the back closer to the floor. Carefully lower it into the enclosure and align it with the mounting holes. See Figure 11 for reference. Note: You may have to turn the bulkhead diagonally to fit it into the enclosure.
3. Using the small M3 screws, attach the Bulkhead Panel to the Enclosure. See Figure 11 for reference. Make sure the wires are coming through the gaps at the bottom and are not being crushed by the bulkhead.

#### Part 3 – Fan Mounting

1. Prepare the fan guards on the intake fan side (left side when looking at the enclosure door in this case). The foam filter should be placed in the outer fan guard as shown in Figure 12.
2. Screw in the Mounting Frame into the Fan Mounting holes as shown in Figure 13 using the long M3 screws.
3. Align the mounting holes on the fan with the screws coming through the mounting frame. Screw and tighten nuts on each screw. See Figure 14 for reference
4. Repeat process for the other large fan.
5. For the small fan in the power supply section, you will only need the small metal fan guard attached to the outside. Mount the small fan. See Figure 15 for reference.
6. For the outtake side, the mounting process is the same except there is no fan filter guard. The labels of the fan should be facing out. See Figure 16 for reference.

### VII - Front Panel Switch and Reed Switch

Part 1 – Front Panel and Reed Switch Wiring and Mounting:

1. Mount the front panel I/O switch to the small mounting hole on the front of the enclosure, to the right of the door. Squeeze the tabs on the side of the switch to insert.
2. Using the long red wire from the relay, crimp the other end with a quick connect and connect it to the middle tab of the front panel switch, as shown in Figure 17.
3. 3. Prepare a red wire with one end crimped with a quick connect tab. Leave the other bare.

**Note**: If you’d like to test the reed switch positioning before powering the system, you can skip to step 6. Use a multimeter to check the connection once the magnet is glued

1. 4. Prepare the reed switch. Strip one end of one of the wires (it doesn’t matter which one) and connect it to the wire from step 3 (either by soldering it or attaching connectors). Connect the quick connect tab to the one remaining silver tab on the front panel switch.
2. 5. Take the other wire of the reed switch and strip the end. Connect the end of that wire with the long red wire coming from the DC power supply.
3. 6. Using double sided VHB tape, attach the reed switch to the inside of the front of the enclosure next to the door. **Note** you may need to adjust the positioning so that it activates when the magnet on the door latch comes close to it. Refer to Figure 18 for appropriate positioning.
4. 7. Take a small metal magnet and super glue it to the latch of the door on the inside, as shown in Figure 18. You will need to let it sit depending on the working time of your chosen adhesive. **The positioning should be such that the reed switch should activate when the latch is fully closed (turned all the way clockwise)**.

#### Part 2- First System Check

At this point of the assembly, it may be helpful to check your connections and power on the system to see if the fans turn on. **Before powering on the system, use a multimeter to check for a short between the positive and negative sections of the terminal block**. If there is a short, refer to the Troubleshooting Guide for more info.

Open the front door of the enclosure and perform the following steps:

1. Turn on the back panel switch to the “I” position.
2. Turn on the front panel switch.
3. Close the door and turn the handle fully clockwise.

When the door closes, all the fans should turn on, with air flow entering the enclosure on one side and leaving on the other. If one or more of the fans don’t turn on, refer to the Troubleshooting Guide for more information.

### **VIII** - LED Board Ceiling Assembly and Mounting

The next two sections assume that the PCB’s have been prepared and verified. If doing your own PCB assembly, please refer to the PCB assembly section.

1. Prepare 6 LED boards for the ceiling mounting. If the boards have not been assembled and verified, do so at this time.
2. Take one board that has the tab connectors and mount it using M3 screws as shown in Figure 19. Orient the tabs so that they point towards the side edge of the ceiling panel (or towards the edge where there is a large gap between the edge and the mounting holes).
3. Prepare 2 wires: One red and one black. Crimp one end of each wire with quick connect tabs, and the other ends with a ring terminal.
4. Attach the quick connect tabs with the tabs on the PCB. **Check the polarity of the connection: Red should be on the +24V and Black should be on the GND.** You may need to unmount the board to double check. See Figure 20 for reference.
5. Connect the other 5 LED boards. With the hermaphroditic connectors on all 4 edges of each PCB, you should theoretically be able to “drop” them in from above to connect to the adjacent boards. Align the boards with the mounting holes and screw them in with M3 screws. See Figures 21 & 22 for reference
6. Take the wires connecting the PCBs to the terminal bus and carefully push them into the gap at the top of the Bulkhead Panel as shown in Figure 23.

### **IX** - LED Board Floor Assembly and Mounting

1. Repeat Steps 3 and 4 of the previous section. This time, however, orient the PCB connector tabs so that they point towards the Bulkhead Panel, towards the back of the enclosure in the center. See Figure 24 for reference.
2. Connect the rest of the boards together as in Step 5 of the previous section.
3. On the 4 corners of the PCB array, insert an M3 set screw. **Make sure to not screw them in all the way, as they will be used to mount the Sample Stage**. Screw on the M3 standoffs to the set screw until tight. See Figure 25 for reference.
4. Connect the rest of the boards and mount them with M3 screws.

### **X** - Sample Stage and Spill Protection Assembly

To protect the electronics on the bottom of the photobleacher, it is important to install a spill protection bezel onto the glass stage. The 3D models are available in the supplemental materials section and may be printed on a large-format 3D printer such as the Prusa XL.

#### Part 1 - Preparing The Photobleaching Stage

1. The photobleaching stage should be made from borosilicate glass according to the drawings in the supplemental materials. Our design uses glass of 1/4” nominal thickness but 5 or 6mm thick glass may be substituted. You may also want to send the stage out or devise an alternative mounting scheme if you don’t have the tools to work with glass properly. **Note: Acrylic may be used in a pinch, but it is susceptible to corrosion from samples.
2. Using the drawing from the supplemental materials, mark out the hole positions on the glass plate with a marker or scribe and drill the holes with a diamond or carbide drill of appropriate size. Make sure to flood with water during the drilling process as it keeps the tool sharp and keeps the glass dust at bay. Make sure to wear a mask and other appropriate PPE when drilling or cutting glass.
3. Check that the completed holes have the correct spacing according to the drawing.

#### Part 2 – Adding Spill Protection Bezel and Stage Installation

1. Prepare a caulk gun with a white or clear RTV Silicone sealant such as Dowsil 732. Place a bead of silicone around the entire inside perimeter shelf of the plastic bezel. The silicone bead should be approximately 3-6mm wide. You do not need much silicone to make a good seal, so you can add some extra and scrape off the excess with a razor blade.
2. Lower the drilled glass stage into the bezel and press gently down to seat it in place. Do not worry about excess silicone sealant at this time. If you see any gaps in the seal between the glass and the bezel, either fill them in, or remove the glass panel and start over. Do not press down too hard on the glass; a thin but finitely thick layer of silicone is necessary for a good seal.
3. Allow the silicone to cure for at least two hours before handling.
4. If there is excess silicone intruding on the stage area, gently trim it off with a razor.
5. Use 4x M3x8mm socket head screws to gently secure the bezel retainer piece in place. It may be necessary to tap the 3D printed holes in the bezel. The bolts need only be installed with the gentlest of torque as the retainer will deform if they are tightened too far. The retainer is not strictly necessary since the bezel bears no weight, but the retainer ensures that it does not fall off if it sustains a great impact. See Figure 27 for reference.
6. Lower the stage assembly onto the standoffs inside the lower photobleacher housing. Secure in place with 4x M3x10 bolts. Use an M3 rubber sealing washer (like McMaster P/N 99604A141) to seal the bolt head against the glass. See Figure 28 for reference.

### **XI** - Final System Check and First Turn-On

Make sure all components are connected to the appropriate sides of the terminal bus. Then follow the steps in Section VII, Part 2, and turn on the system. All fans and LED panels should turn on. If anything does not turn on, refer to the Troubleshooting Guide.

#### Closing Up and Final Test

Once all boards are mounted, align the ceiling mounting holes with the screw holes on the top of the enclosure. Secure with M3 screws and tighten. **Note**: If the ceiling doesn’t lie flat on the top, double check the wires coming from the LED board going to the terminal block. Make sure it is in the gap at the top of the Bulkhead Panel. The fully assembled system is shown in Figure 30.

### **XII** - Troubleshooting Guide

**Before checking connections, power down the system, turn all switches to the “OFF” position, and remove the power plug from the back.** You can leave the system on if you want to check voltage levels on any of the connections.

This section should be used if any issues arise during the assembly. Using fine tipped probes with the multimeter is very helpful to debug the system.

If there’s a short between the +24VDC and –VDC (GND) sides of the terminal bus block:

1. Disconnect everything from the terminal bus and check for a short again. There shouldn’t be a short with everything disconnected. **If there’s still a short, the terminal bus may be bad, or there may be some metal from the enclosure connecting the two sides**. In either case, remove the terminal bus to double check. If there’s still a short, you will need to use a new terminal block. If not, check for exposed metal on the enclosure near the terminal bus mounting location.
2. Check for shorts at the following:

a. The DC output connector of the DC power supply. Check between the –VDC line (black wire) and the +24 DC line (both red wires).
b. The power input to the relay (the soldered wires). Check between the –VDC line (black wire) and the +24 DC line (red wire). You may need to remove any heat shrink placed on the tabs to check.

If there is a short at these connections, remove the wires and check the terminals on the relay and DC power supply. If there is still a short, it’s likely the component is damaged and needs to be replaced. If not, re-crimp/re-solder where appropriate and check again.

If the System doesn’t turn on:

1. Remove the fuse drawer from the power input module from the back and check the fuse with the multimeter. You should observe a short if the fuse is still good. If there is no connection, replace the fuse (you can also use a fuse with a higher current rating if it’s the same size as the one used here).
2. Check the connection between DC power supply and the terminal bus. Check both +24VDC and –VDC lines.
3. Check the connection between the DC power supply and the relay, as well as between the relay and the terminal bus.
4. Check the positioning of the reed switch. Depending on the location of the magnet, the reed switch position may need to be adjusted. **The system should power on when the door handle is turned all the way in the clockwise direction**.
5. Double check the connections between the relay and the front panel on/off switch.

If one or more fans don’t turn on:

1. Remove the fan connections from the terminal bus. Check the positive and negative connections between the ring terminal connector and the terminals on the fans (you may need to remove the label to access the fan terminal). If there is no connection, repeat the steps of Section VI and fix as necessary. Double check for a connection before reconnecting to the terminal bus.
2. If rewiring is necessary, try one of the other wiring suggestions mentioned in Section VI to reduce the chance of an unconnected fan.

If one or more LED boards don’t turn on:

1. The LED boards should have been verified individually before mounting. Check connection between the power input to the board that is not turning on and the +24VDC side of the terminal block. Repeat for the -VDC side.
2. Double check the board-to-board connectors. You may need to remove the screws to better align the connection. Make sure they are connected as shown in Figure 21.

---

**Figure 1:**
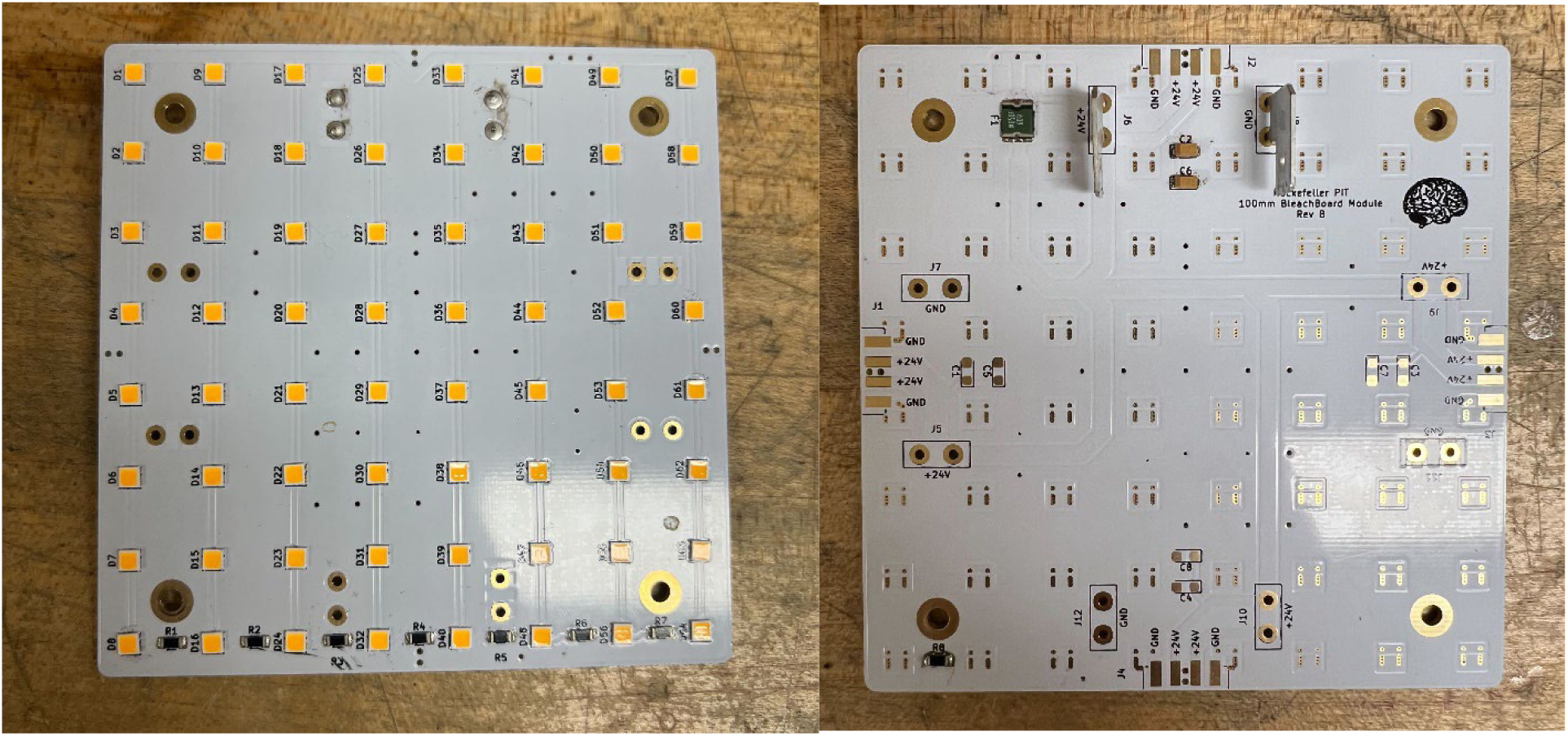
A fully assembled LED PCB board, showing top (left), and bottom (right). The top consists of the LEDs while the bottom consists of the fuse, capacitors, and connectors.

**Figure 2:**
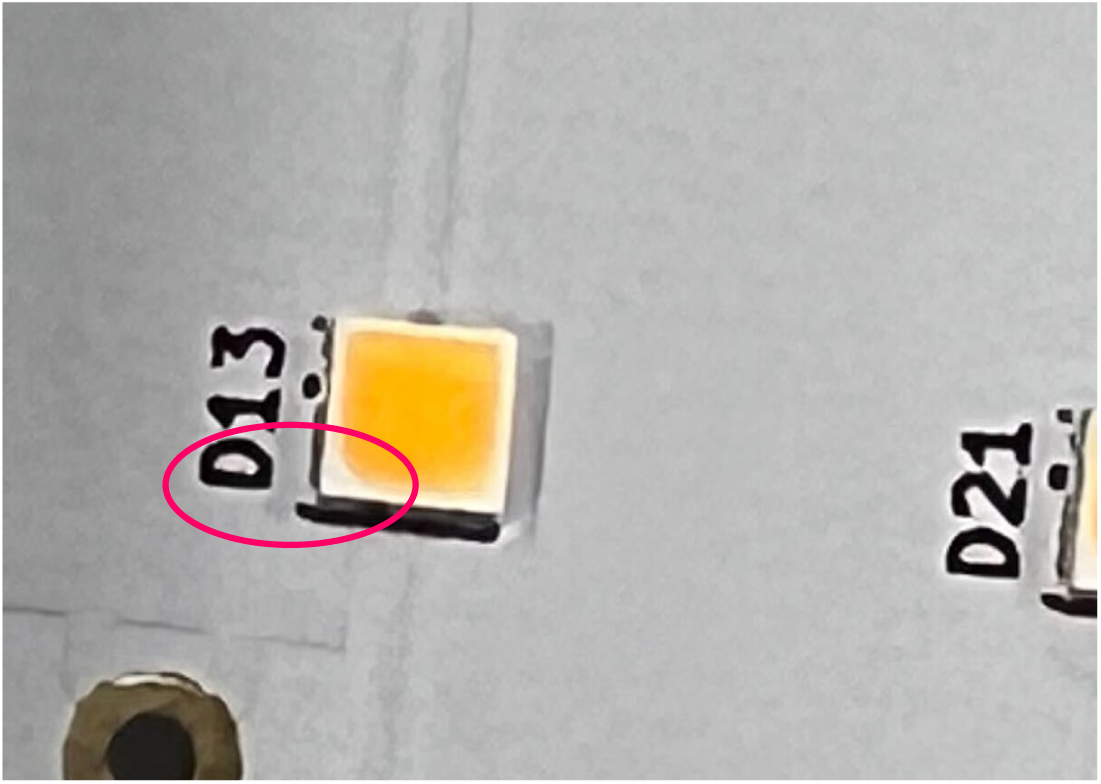
The polarity of the LEDs on the board. Note the white tab, which marks the cathode.

**Figure 3:**
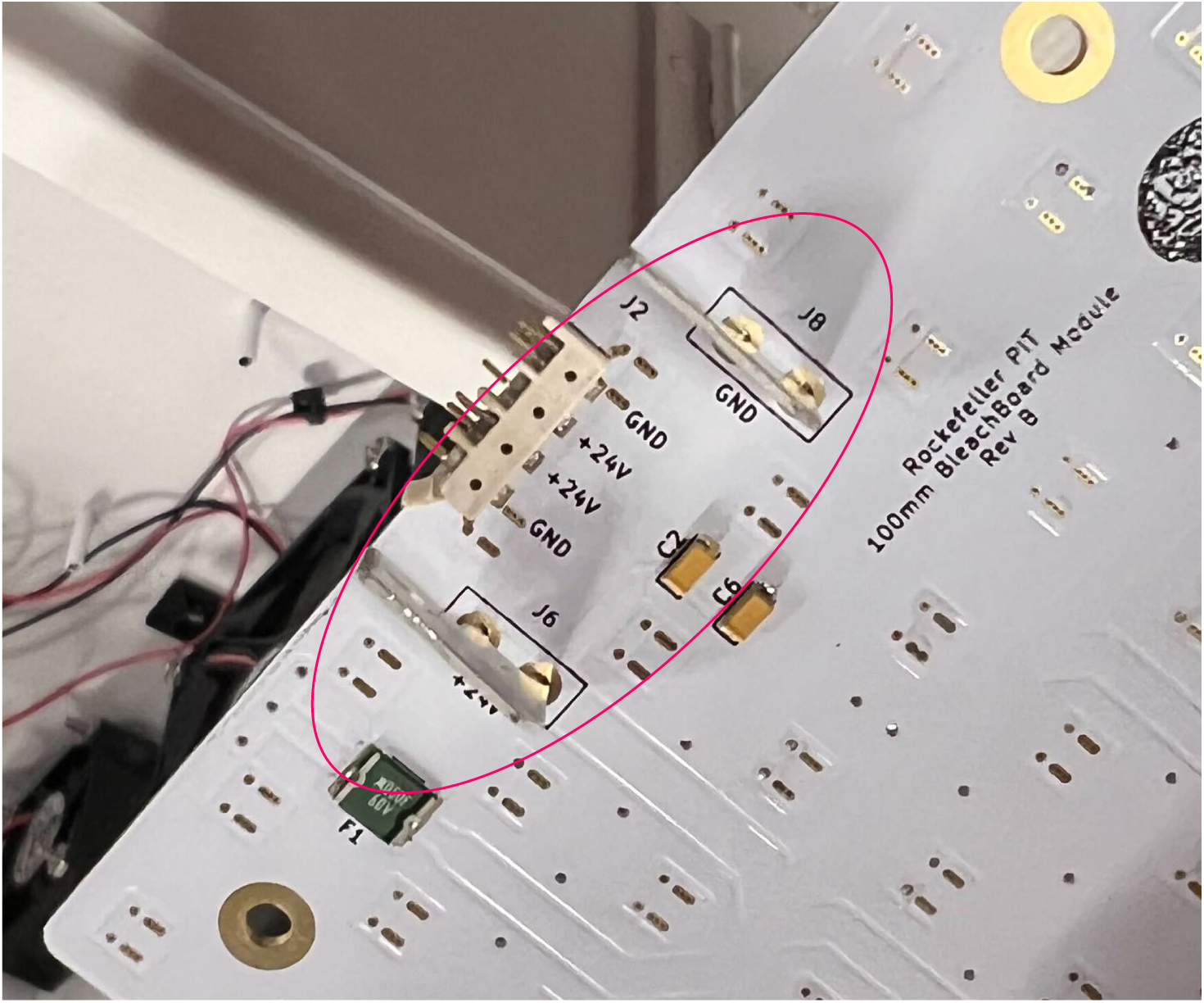
Orientation of the blade connectors. They should be pointed to the edge of the board.

**Figure 4:**
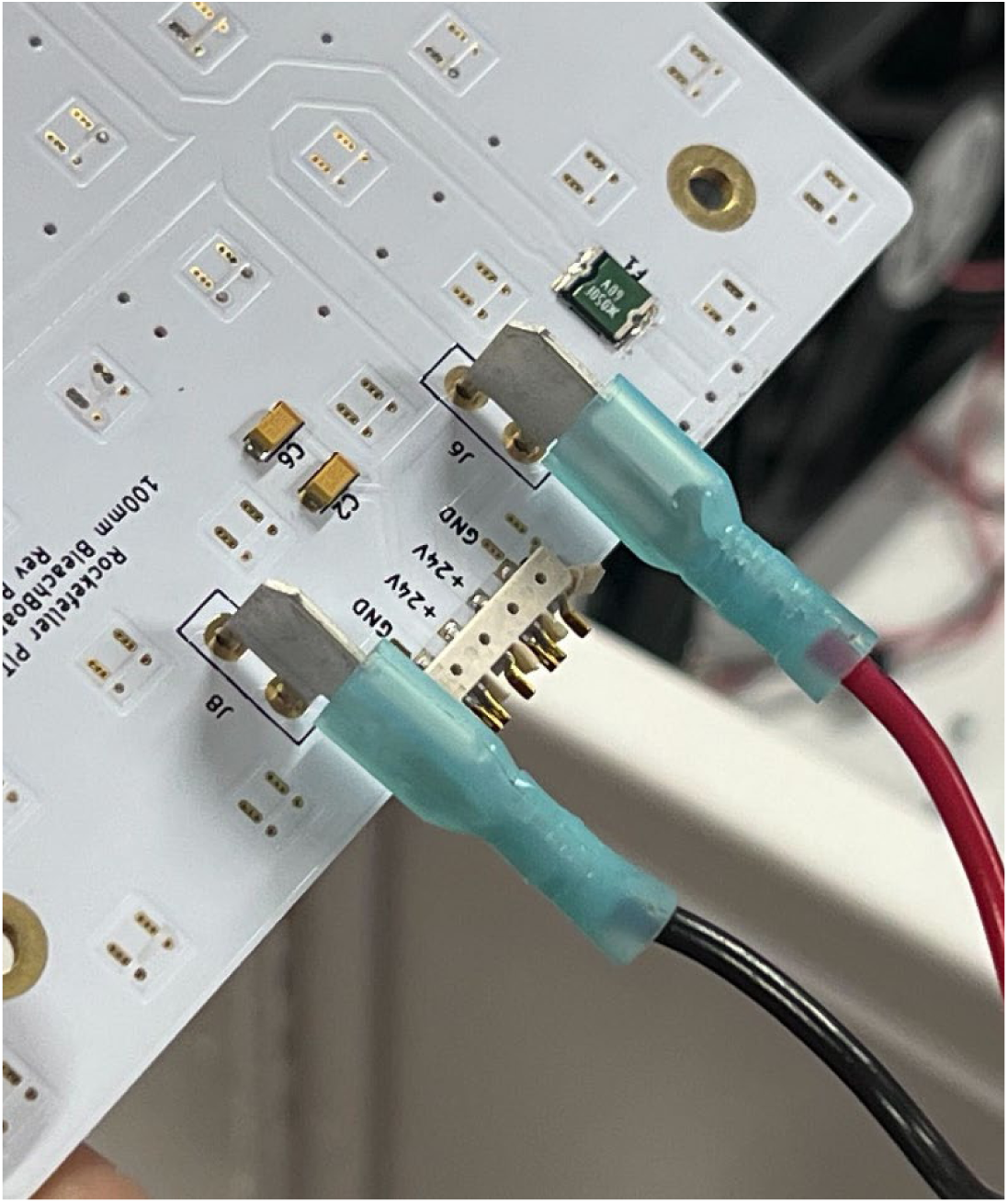
Attachment of wires to the power input blade connectors using quick-tab connectors.

**Figure 5:**
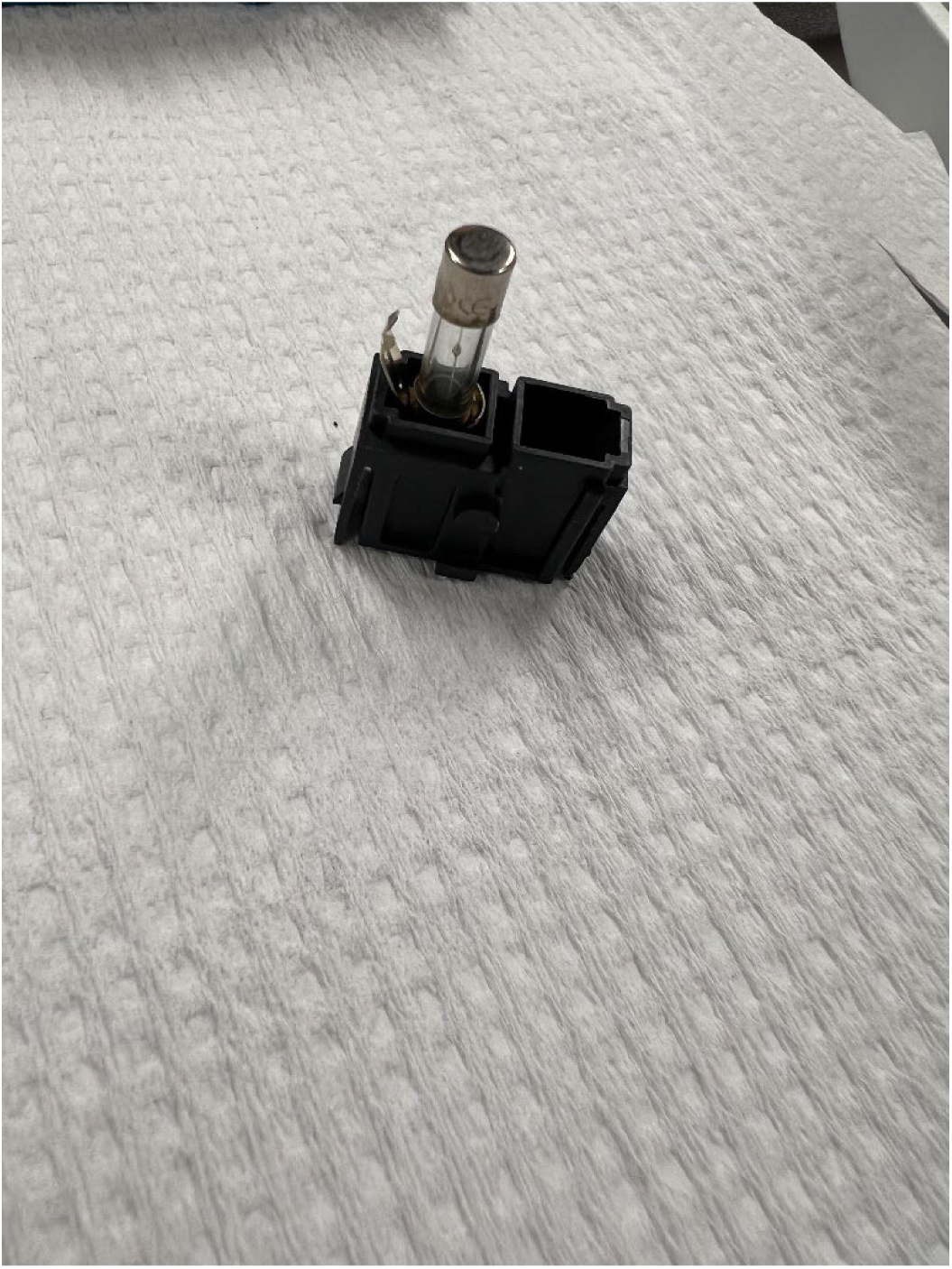
Glass Fuse inserted into the Fuse Drawer

**Figure 6:**
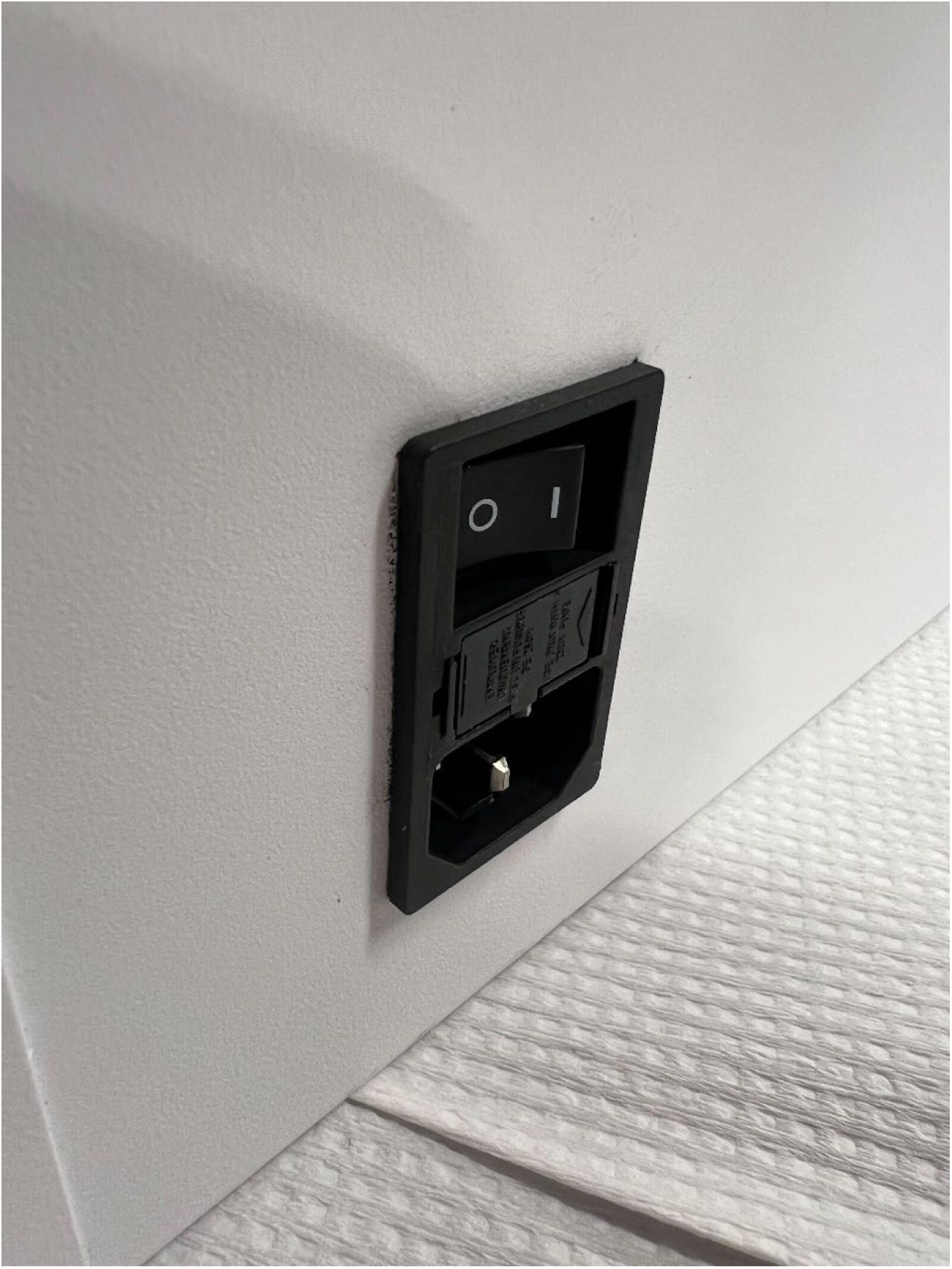
AC Power input module installed in the back panel of the enclosure.

**Figure 7:**
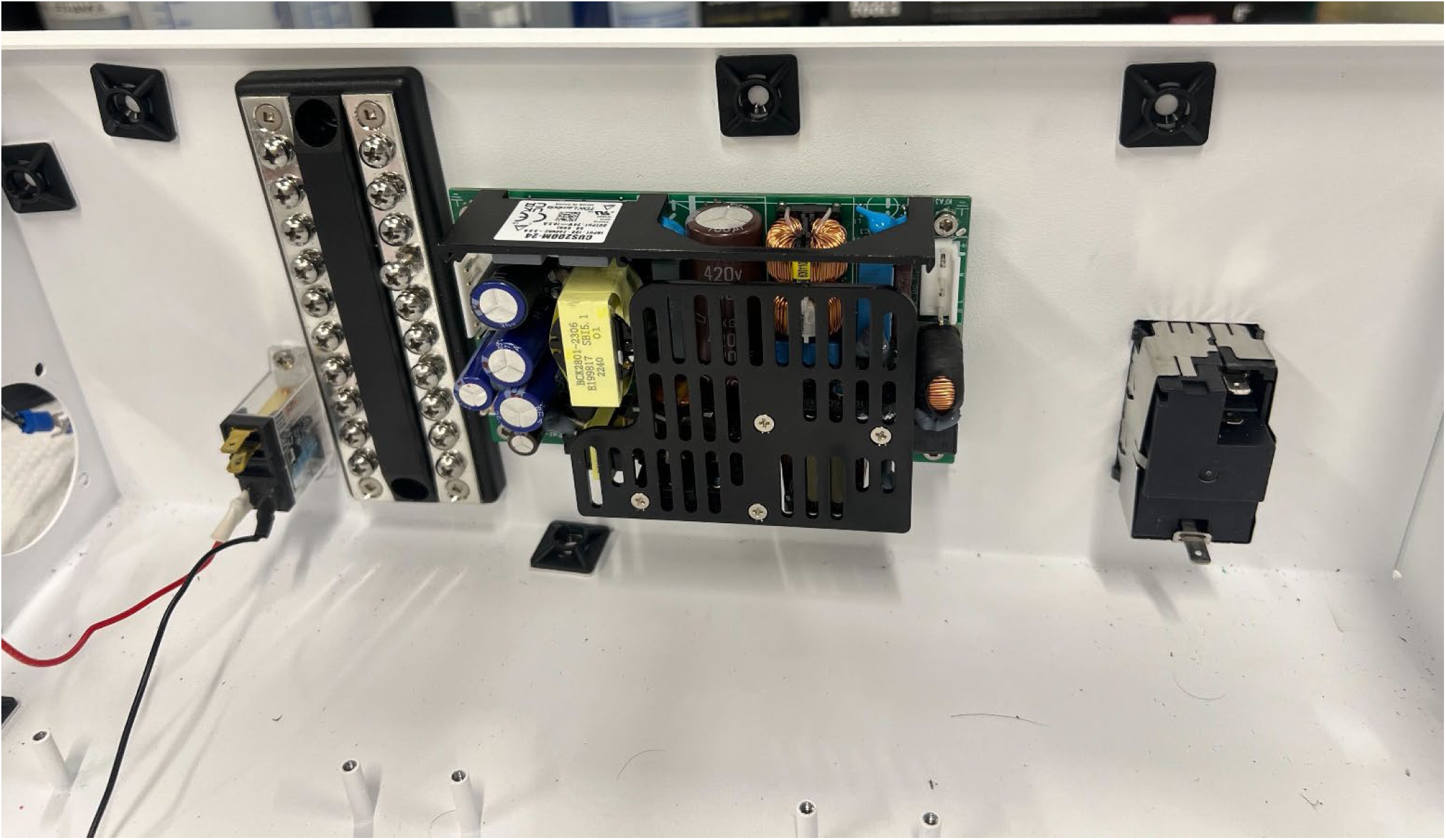
Mounting locations for AC Power Input Module, DC Power Supply, Terminal Bus Block, and Relay

**Figure 8:**
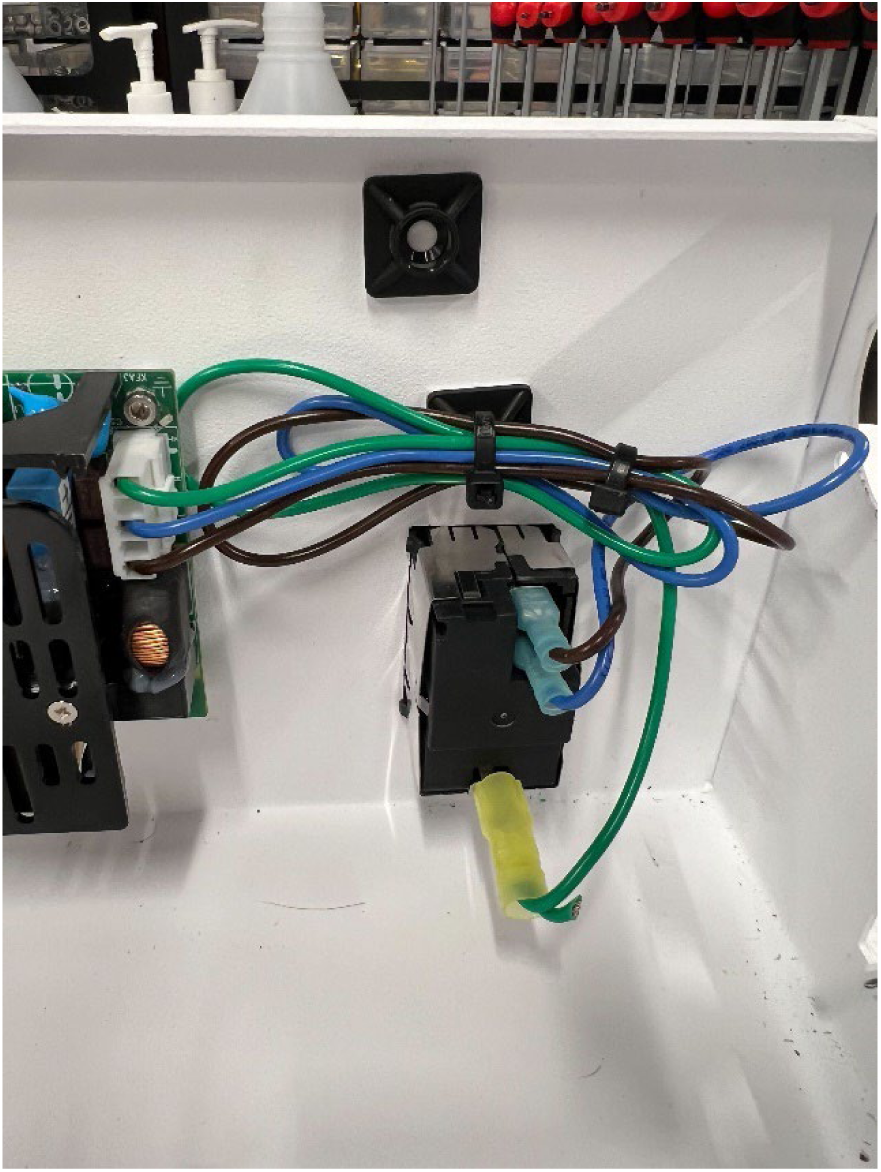
Electrical Connection from the AC Power Input Module to the input of the DC Power Supply

**Figure 9:**
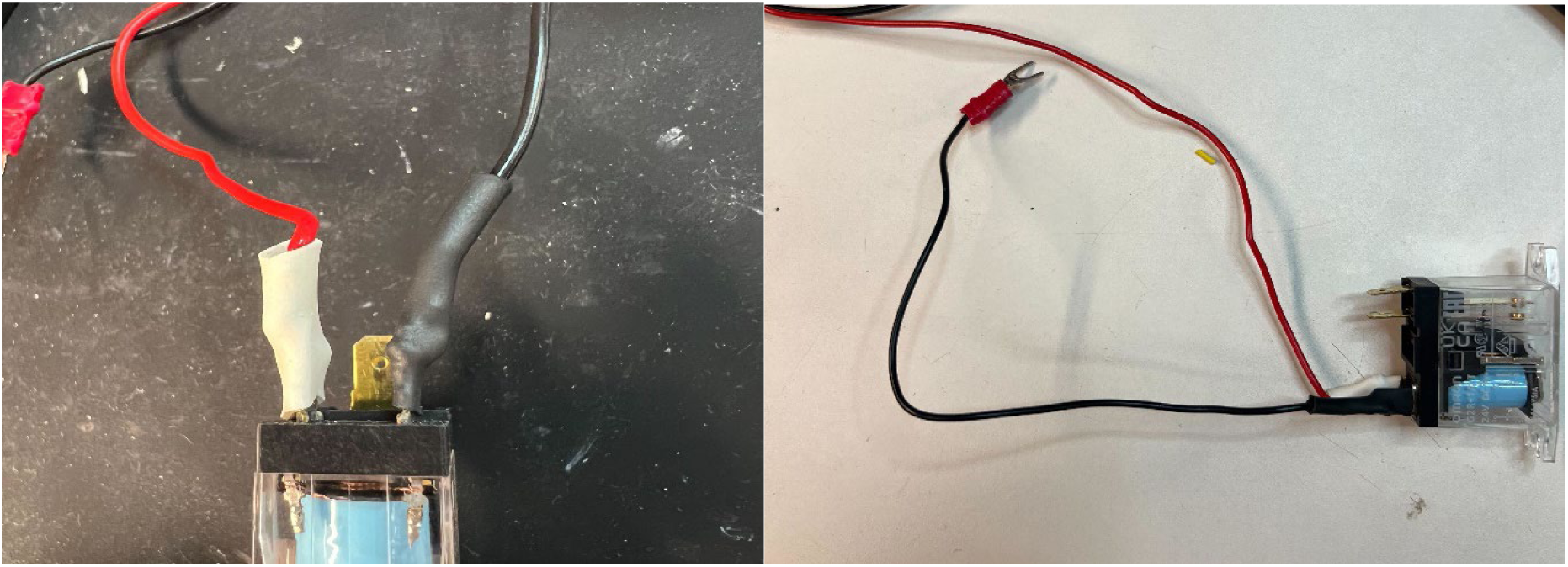
Soldered connections to the Relay. The pins are non-polar can can be switched

**Figure 10:**
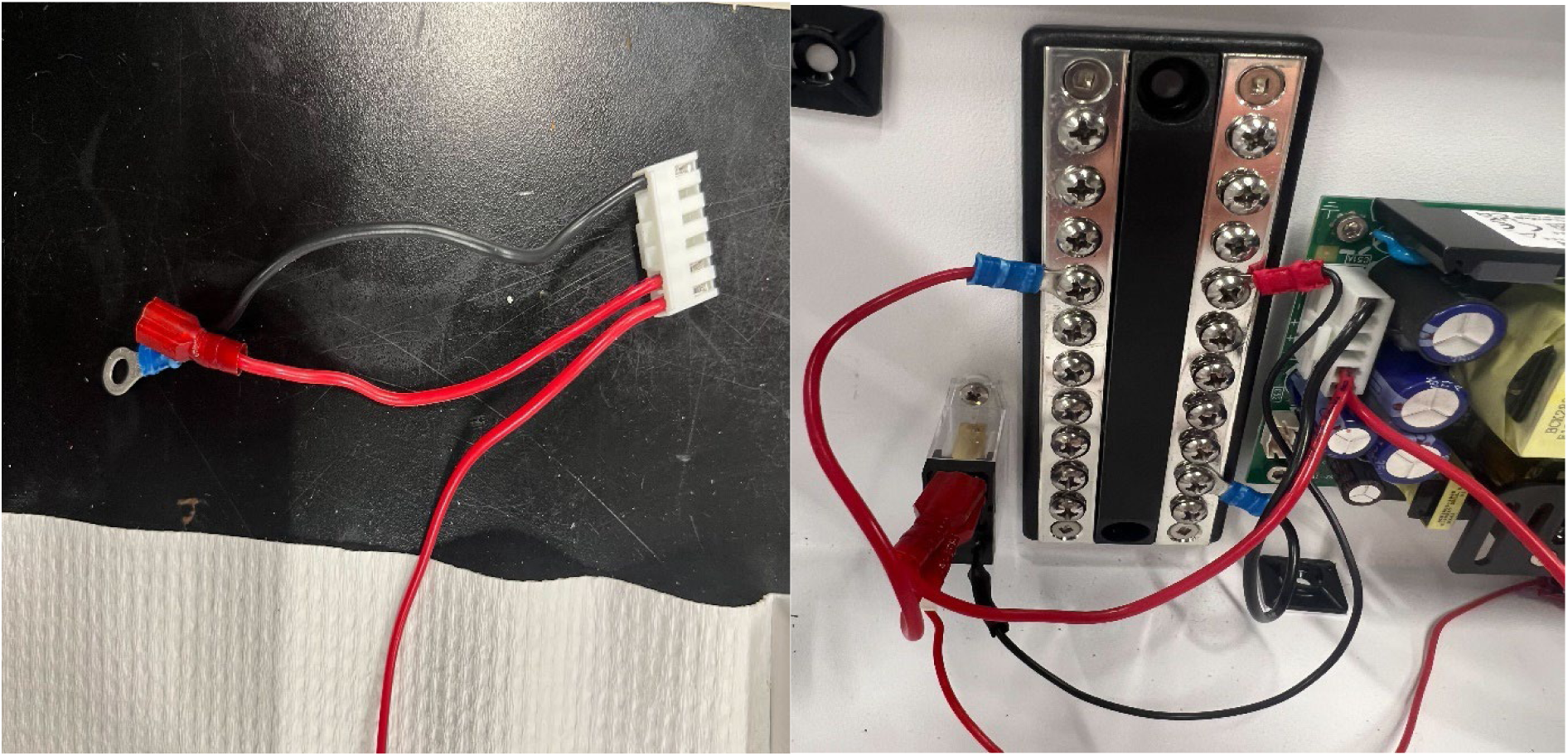
DC output connector (left) and connections for DC Power supply and the Relay to the Terminal Bus Block (right)

**Figure 11:**
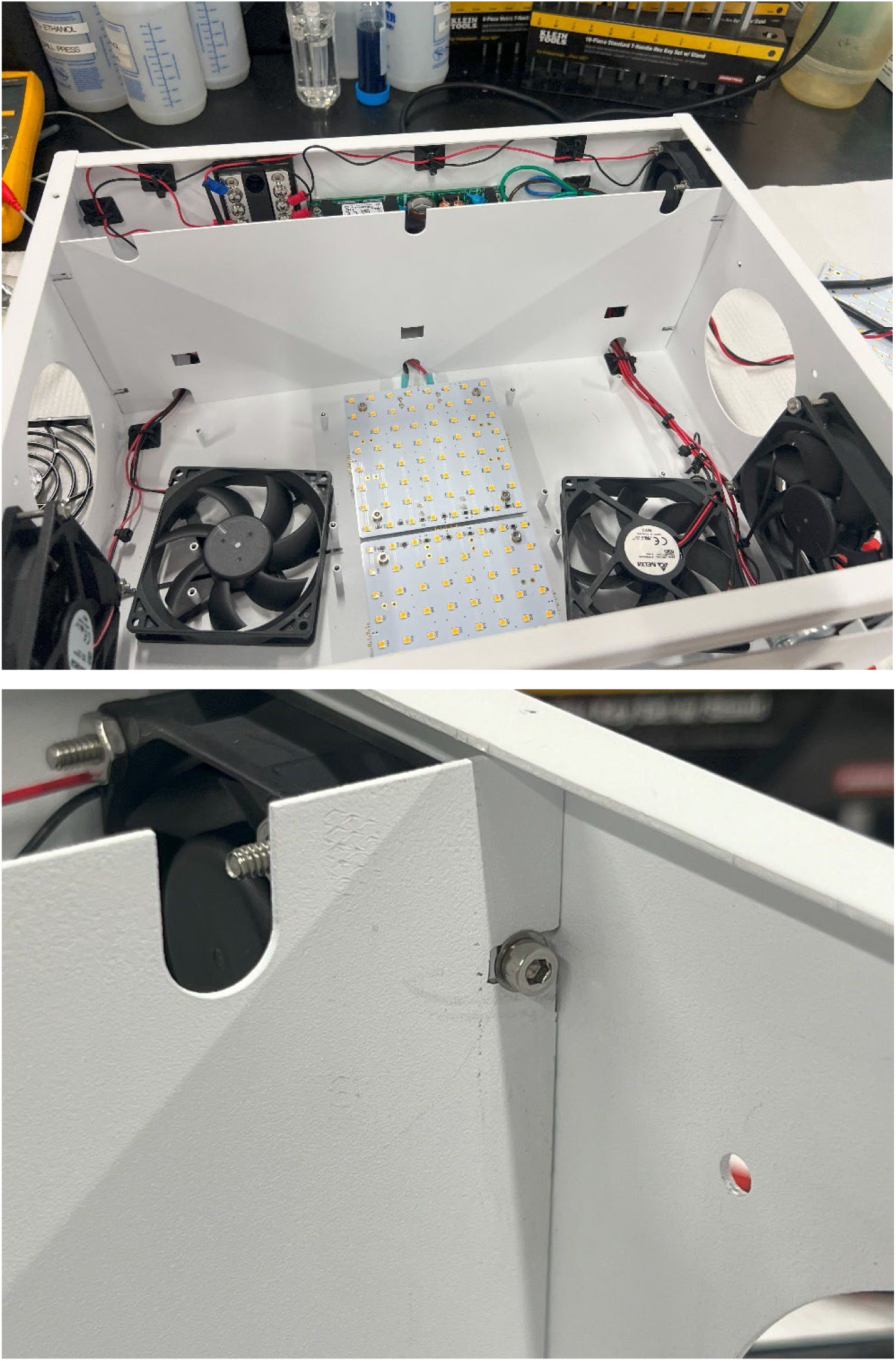
Fully installed bulkhead with wires through the bottom gaps (top). Location of proper mounting with M3 screws (bottom).

**Figure 12:**
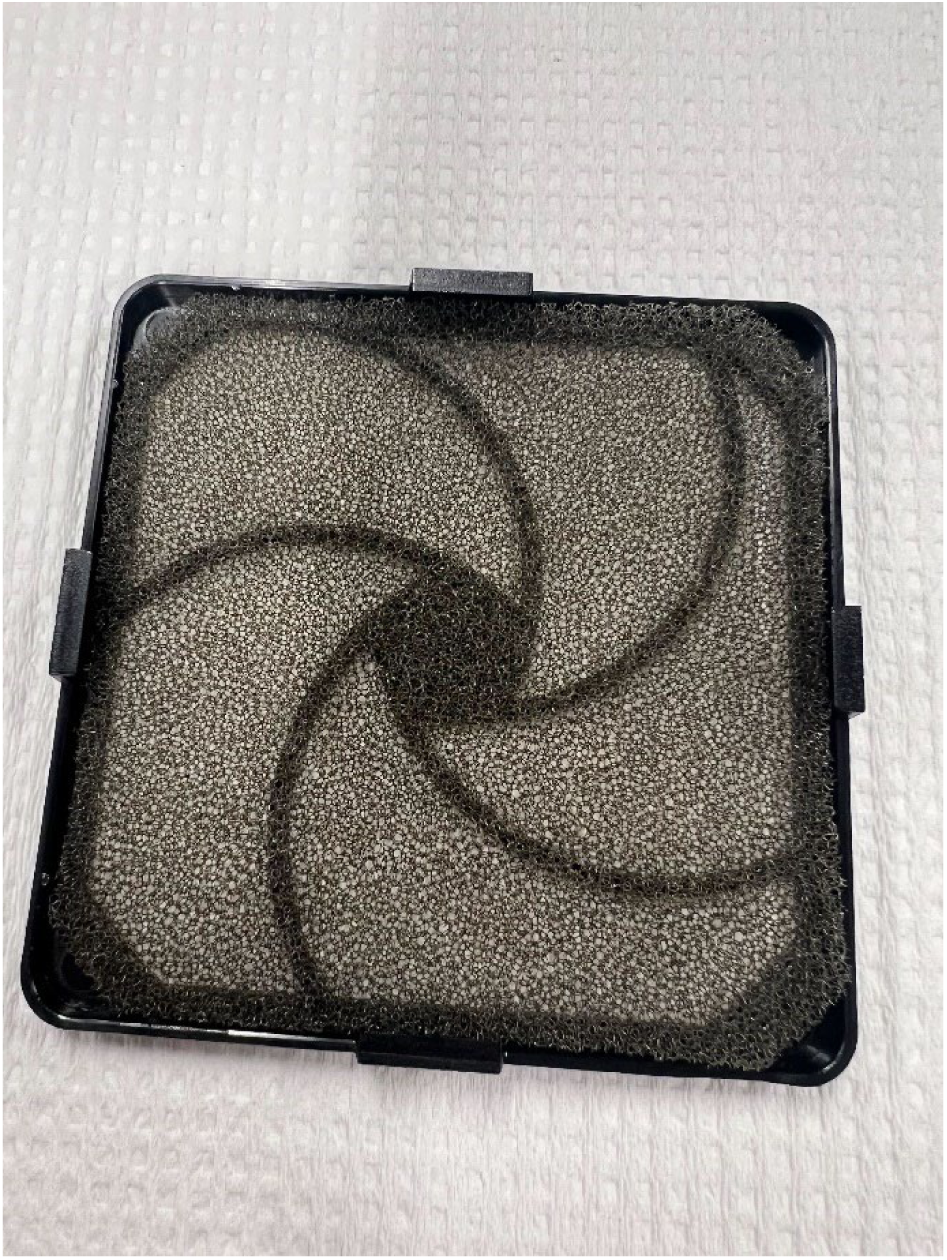
Filter being placed inside the fan snap cover. The snap tabs should be slightly above the foam filter

**Figure 13:**
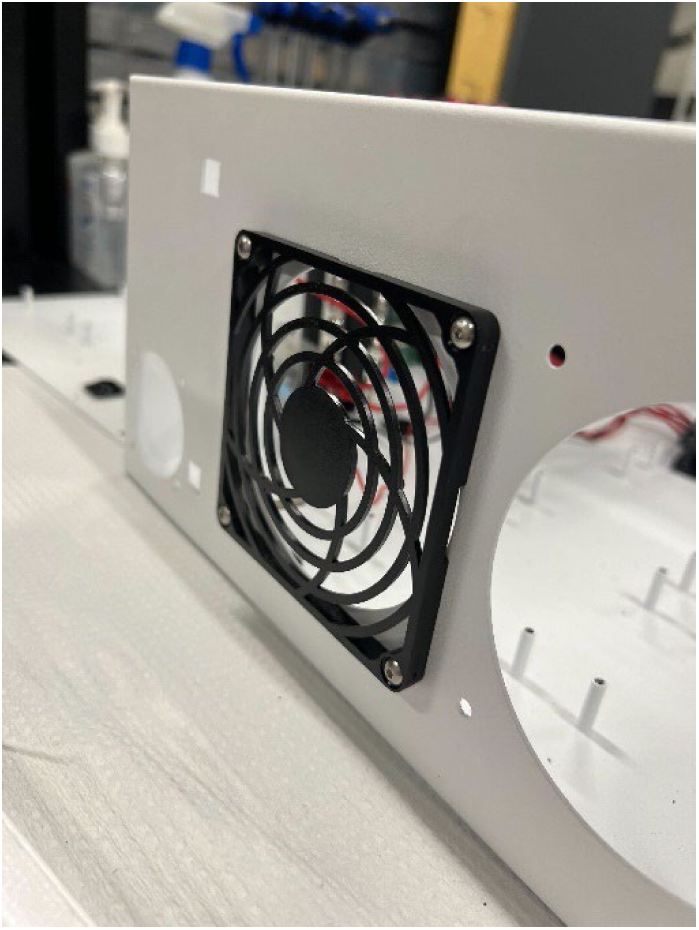
Mounting Frame attachment to the enclosure

**Figure 14:**
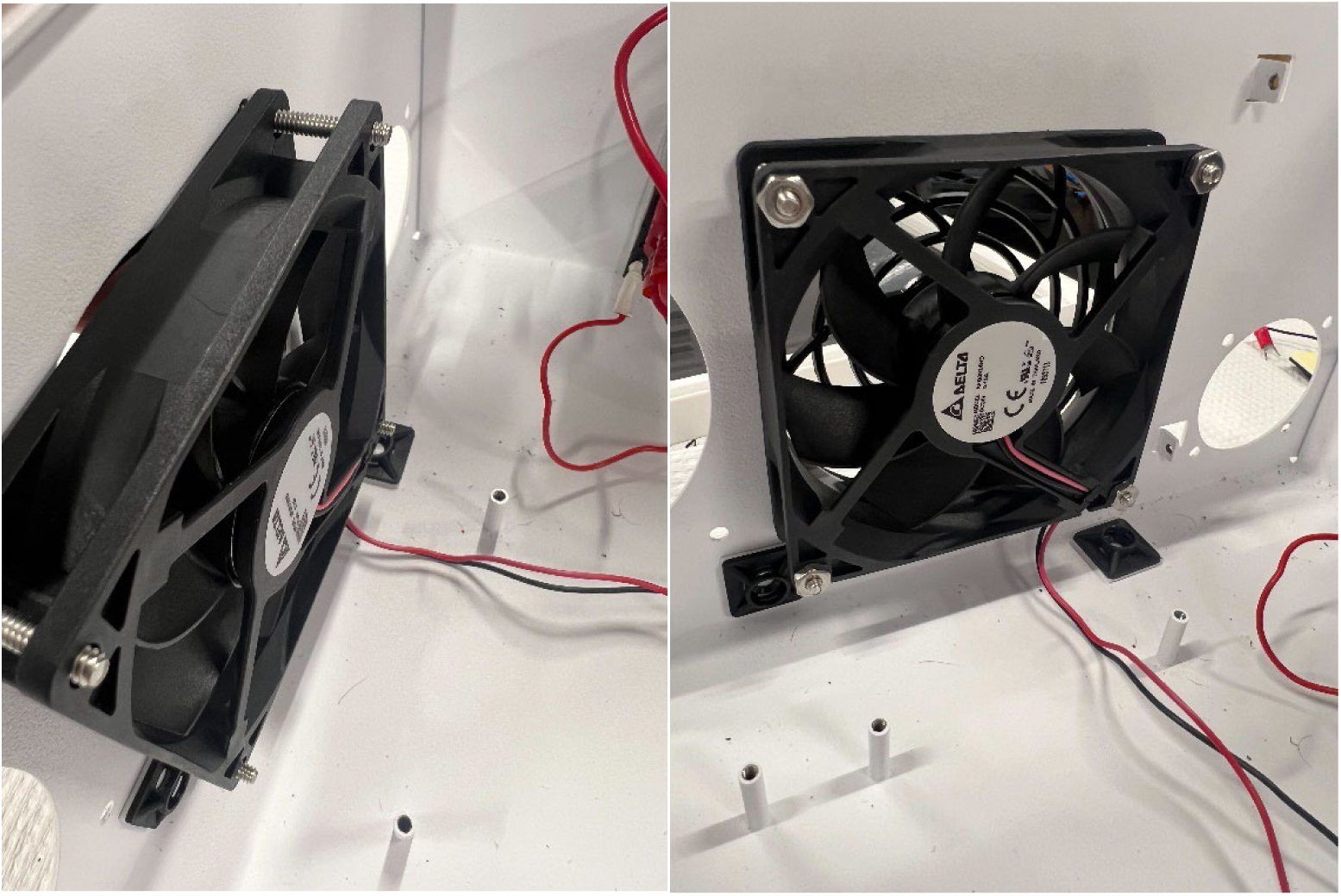
Alignment and mounting of fan with mounting screws (left) and with nuts tightened (right)

**Figure 15:**
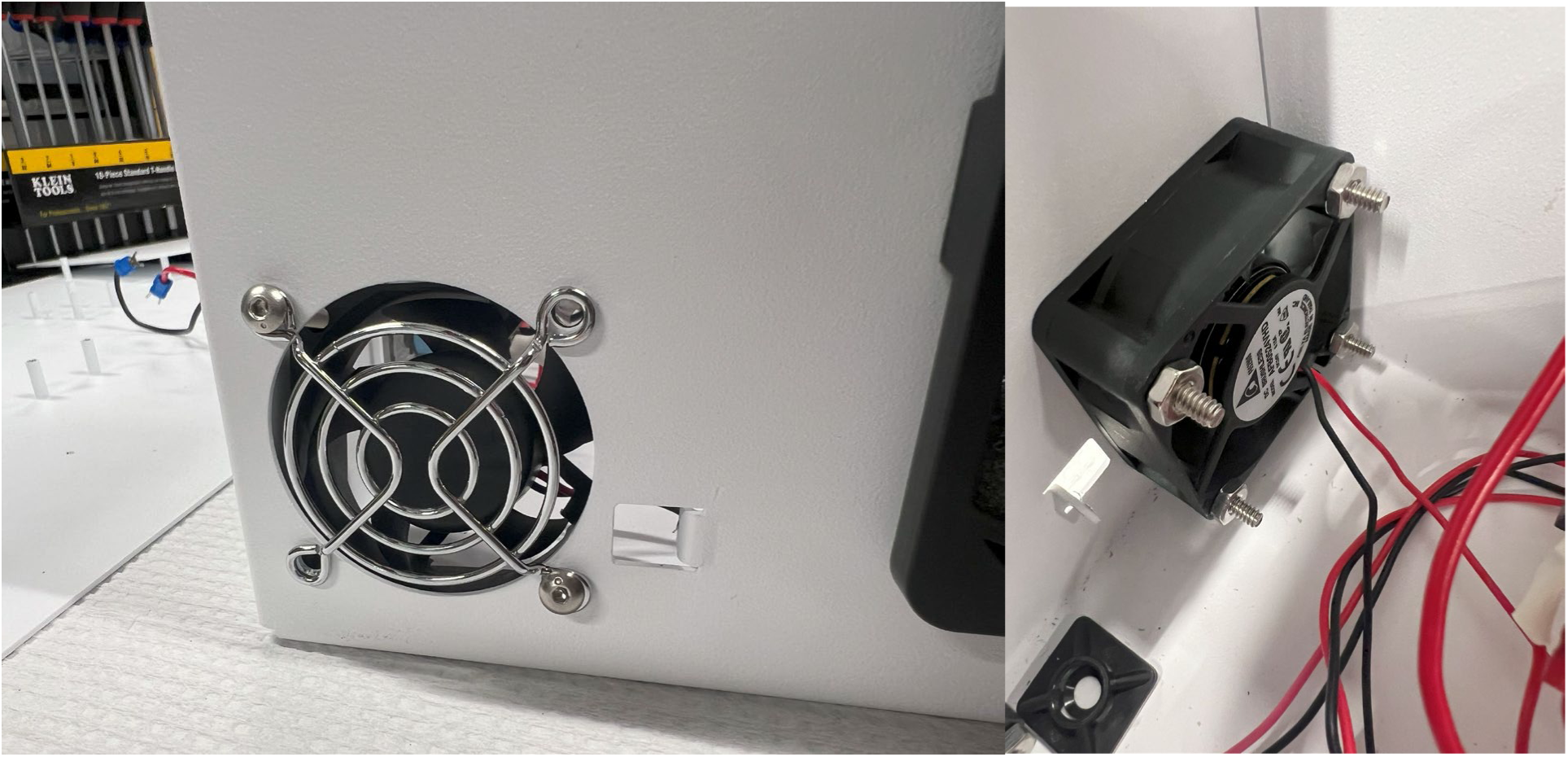
Metal mounting bracket for the small fan attachment to the enclosure (left) and small fan mounted to the inside (right)

**Figure 16:**
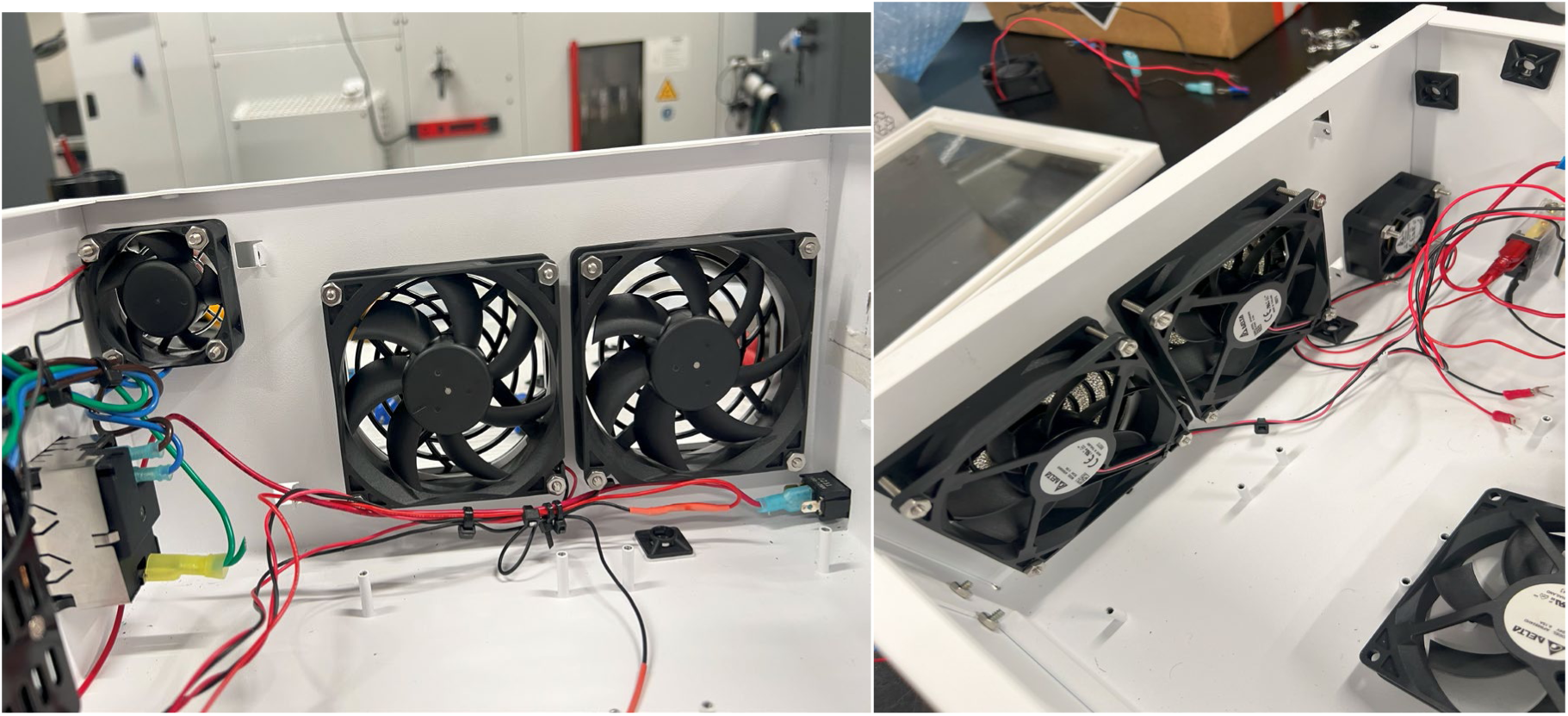
Fully mounted fans on outtake side (right) and intake side (left)

**Figure 17:**
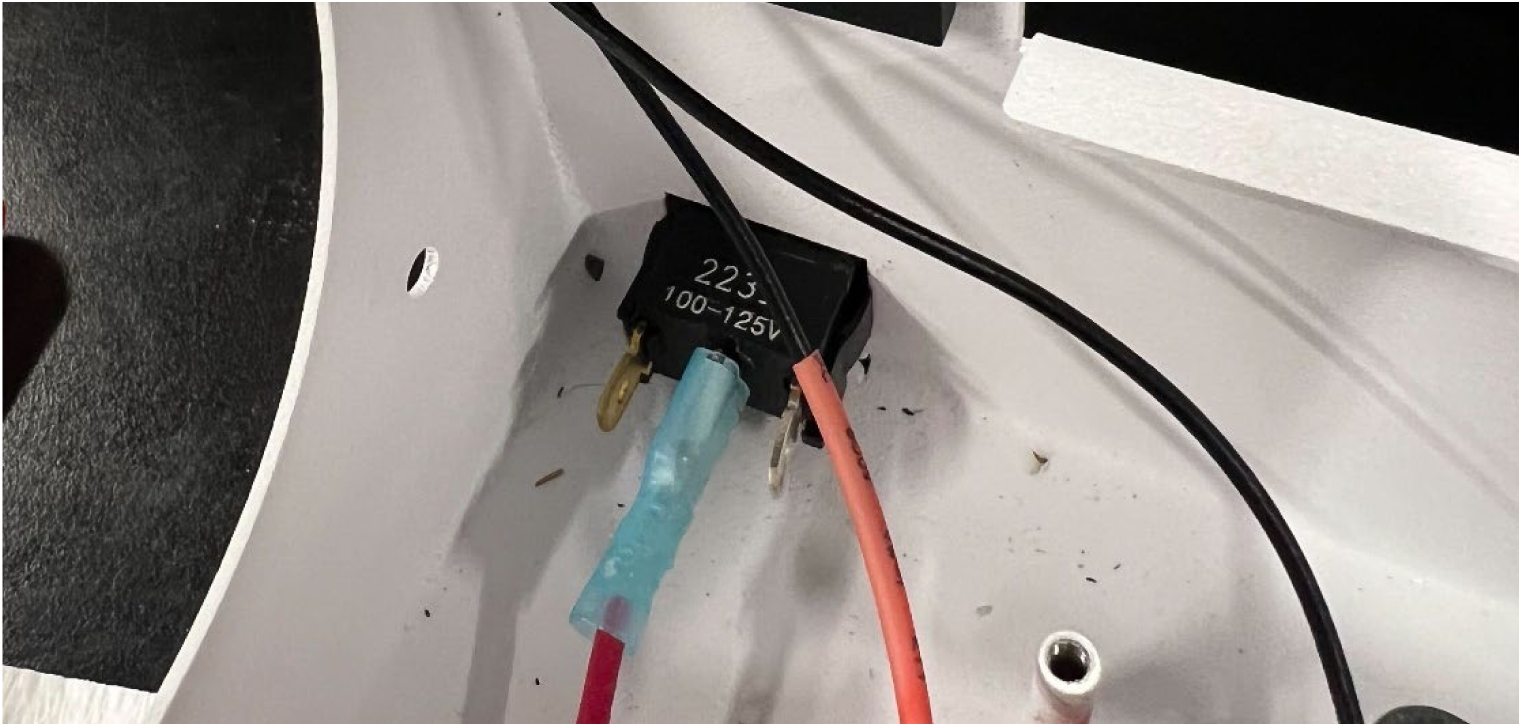
Connection to the middle tab of the rocker switch, which connects to the relay

**Figure 17:**
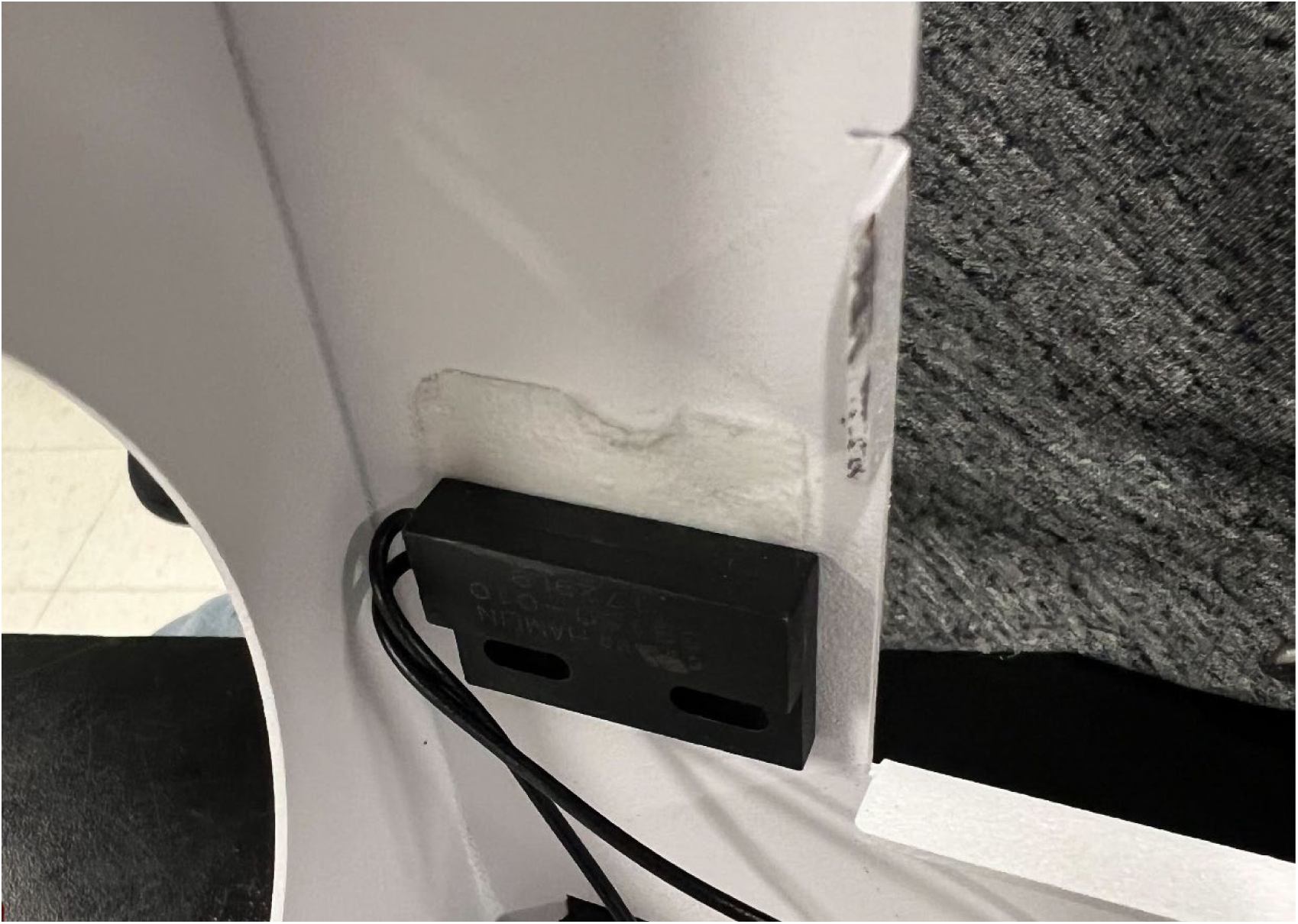
Reed switch orientation and positioning. You may need to test the connectivity to find a more appropriate location depending on where you glue the magnet

**Figure 18:**
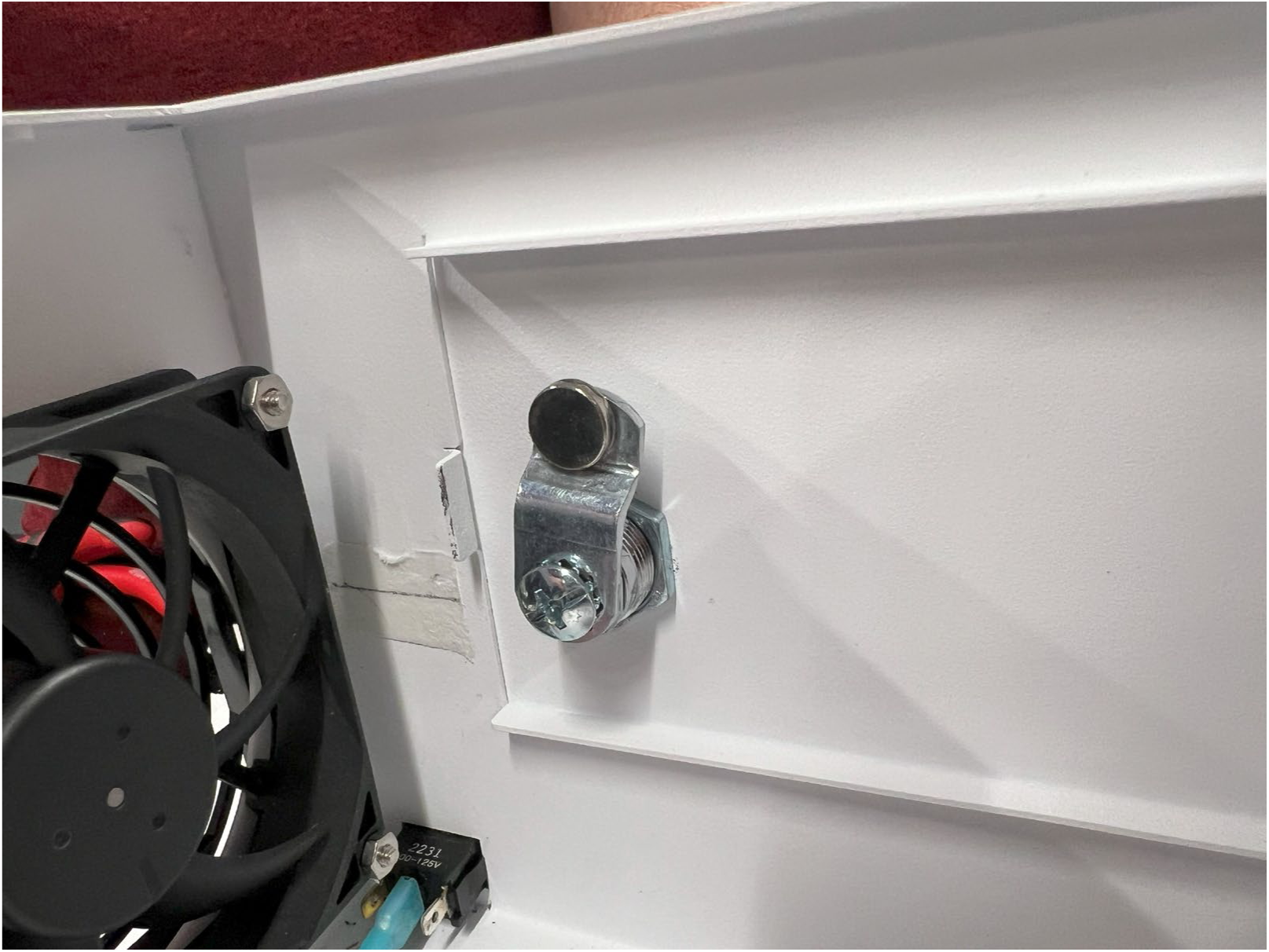
Magnet superglued to the latch on the inside. Depending on the size of the magnet you may need to adjust the position

**Figure 19:**
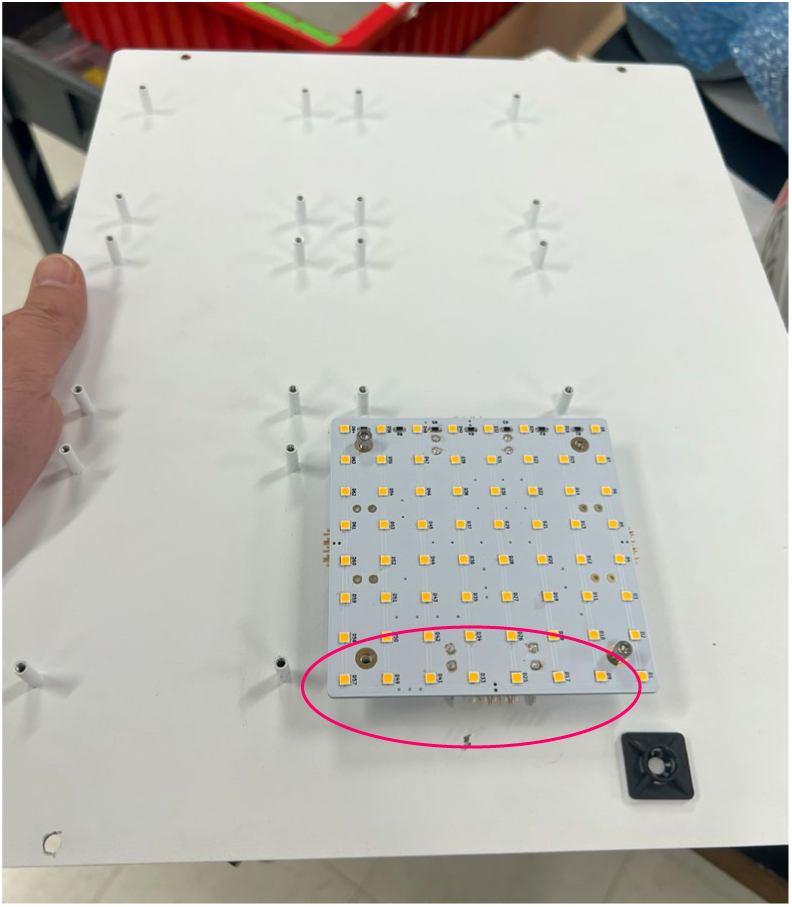
First board orientation and mounting. The pink circle shows the location of the blade connectors, which is the power input to the boards.

**Figure 20:**
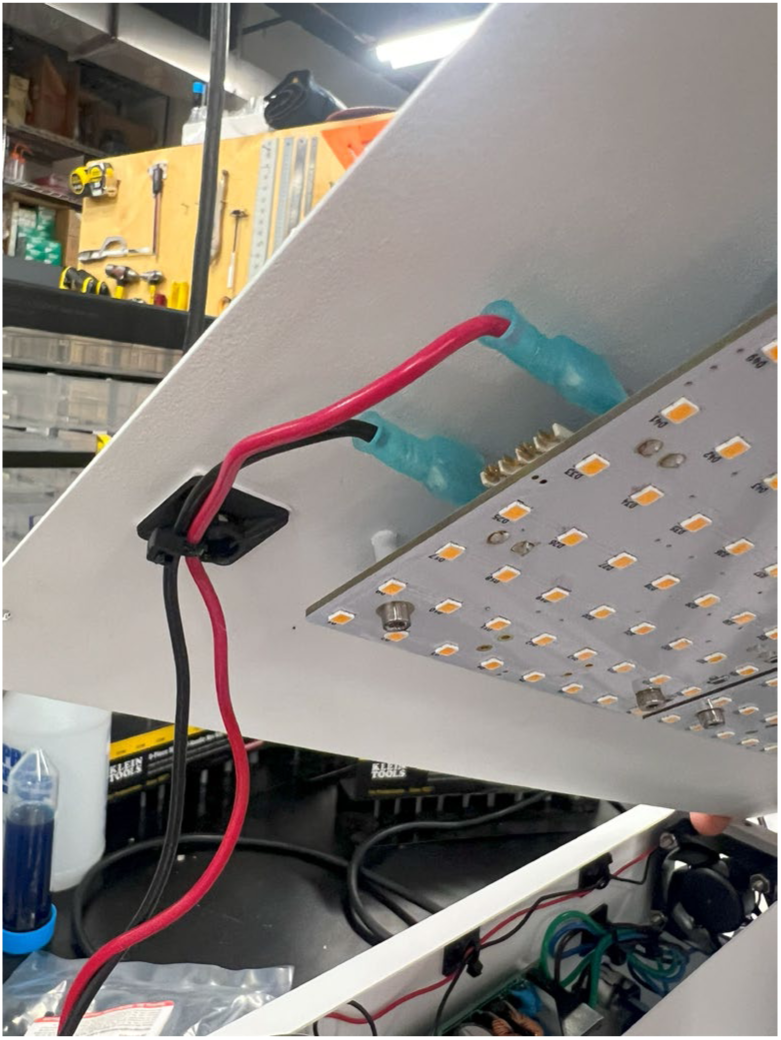
Power input connection to the LED boards. Note the polarity of the tabs.

**Figure 21:**
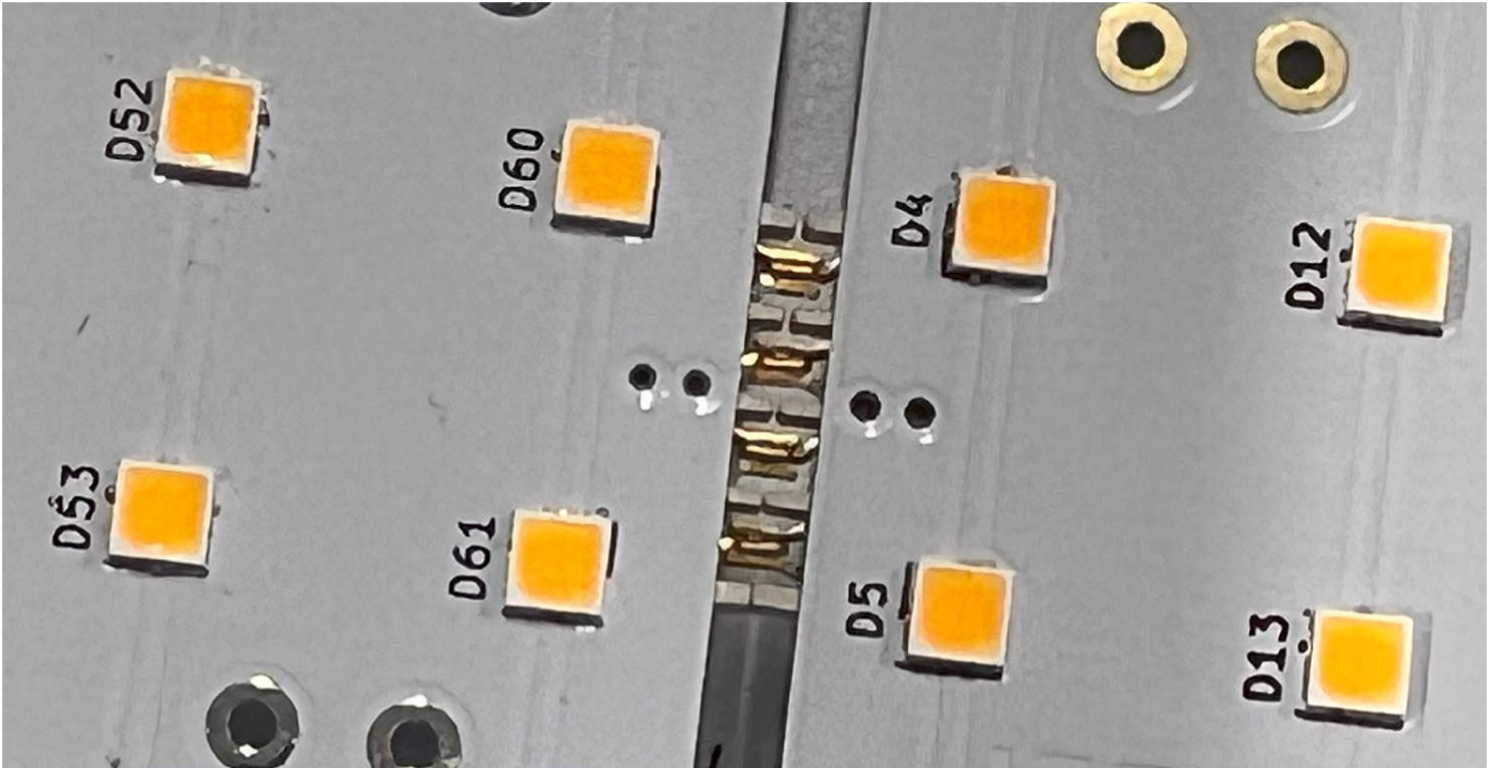
Two LED boards connected using the hermaphroditic interconnector

**Figure 22:**
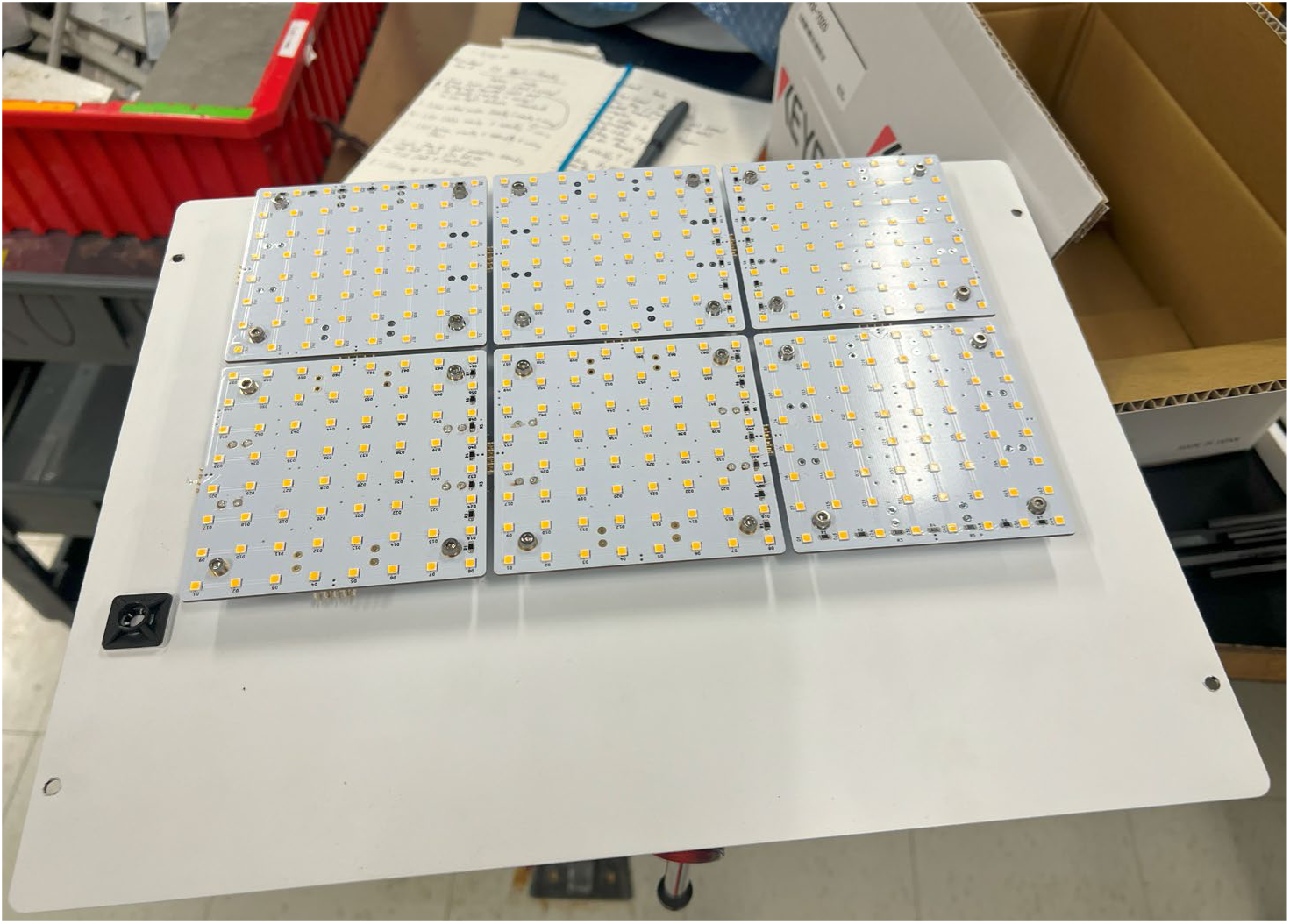
All 6 ceiling LED boards mounted

**Figure 23:**
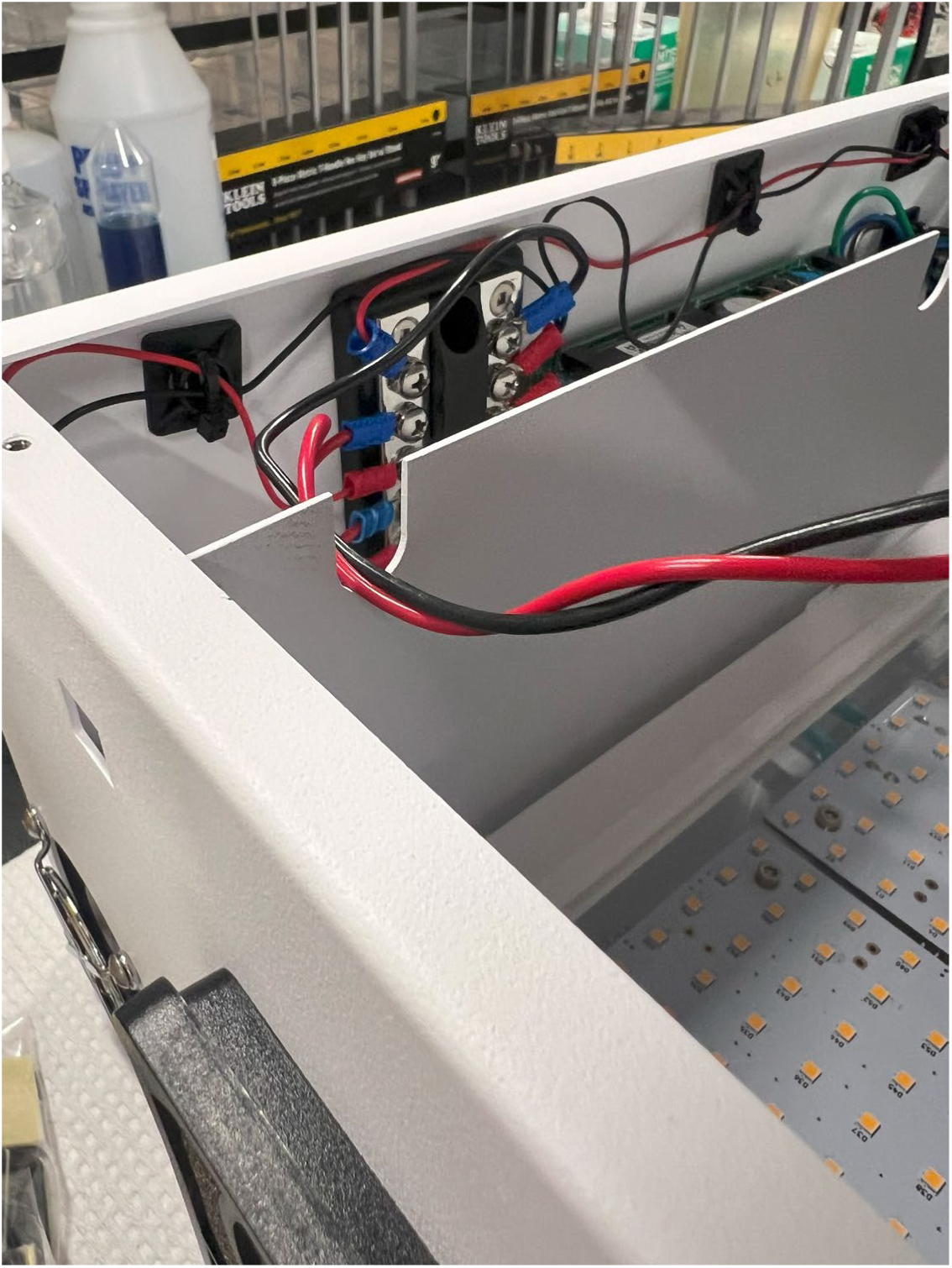
The power input wires from the ceiling LEDs inserted into the groove at the top of the bulkhead panel

**Figure 24:**
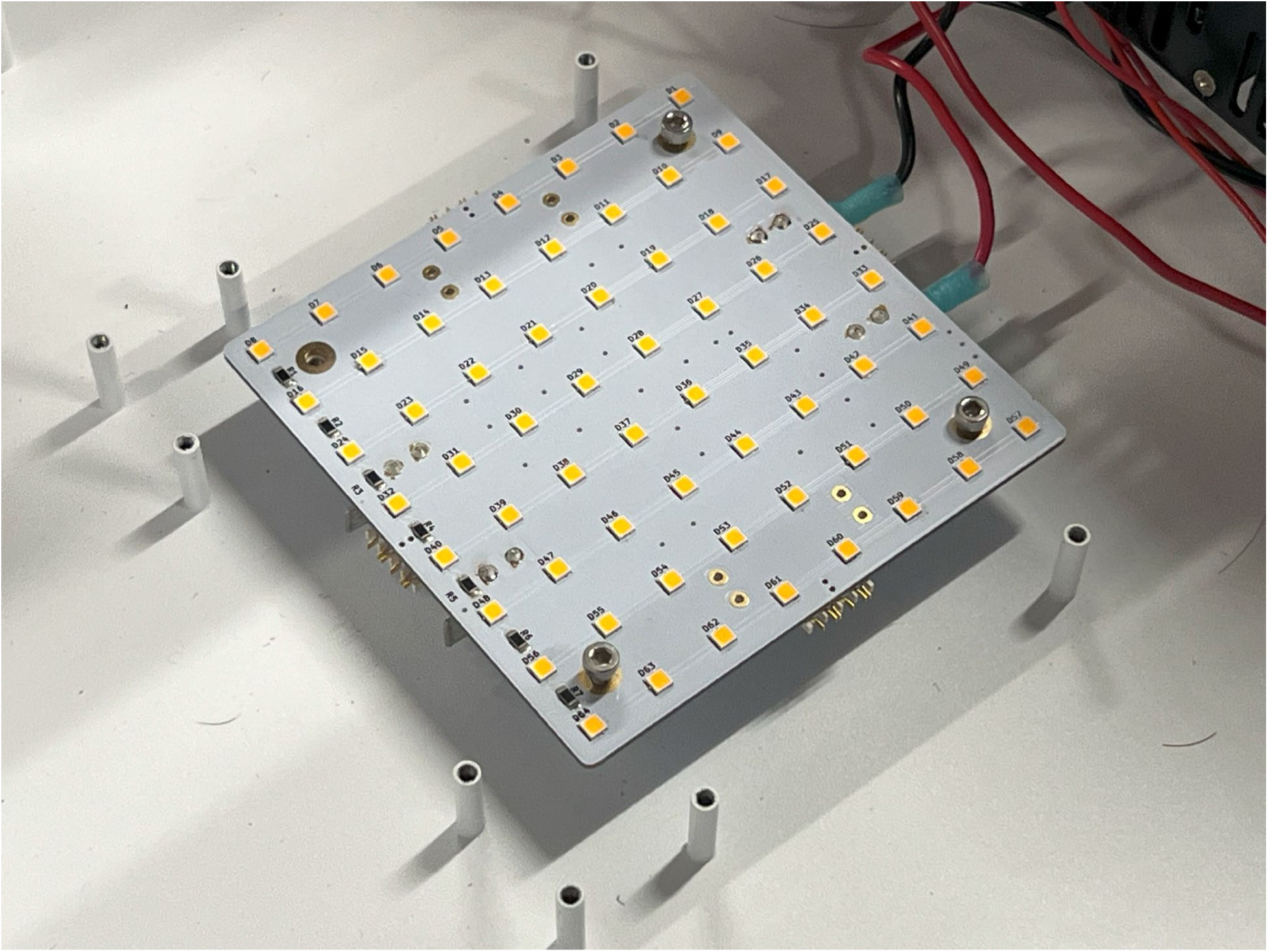
LED board for power input to the floor LEDs

**Figure 25:**
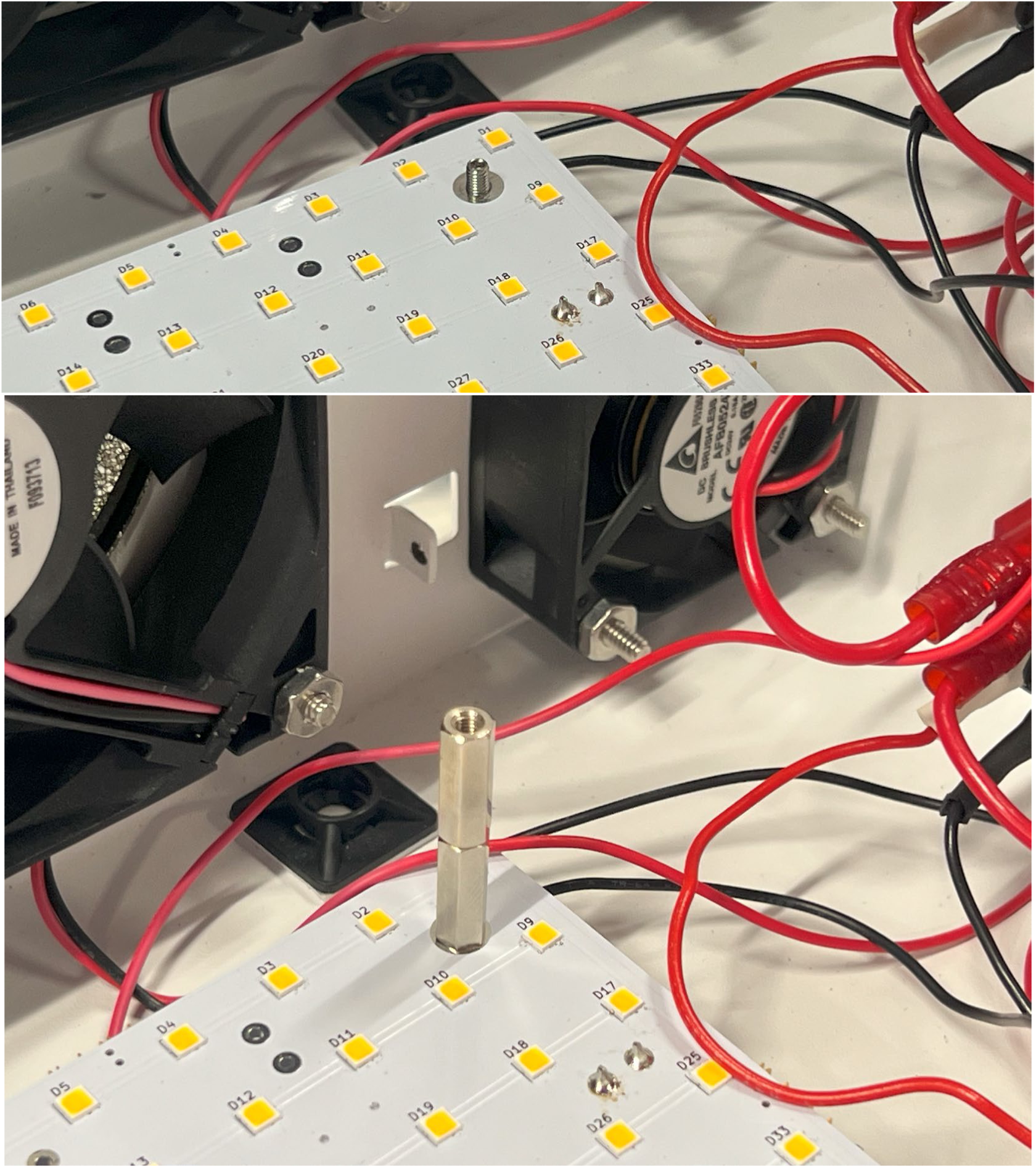
Set screw location and extension above the board (top) with standoffs attached (bottom).

**Figure 26:**
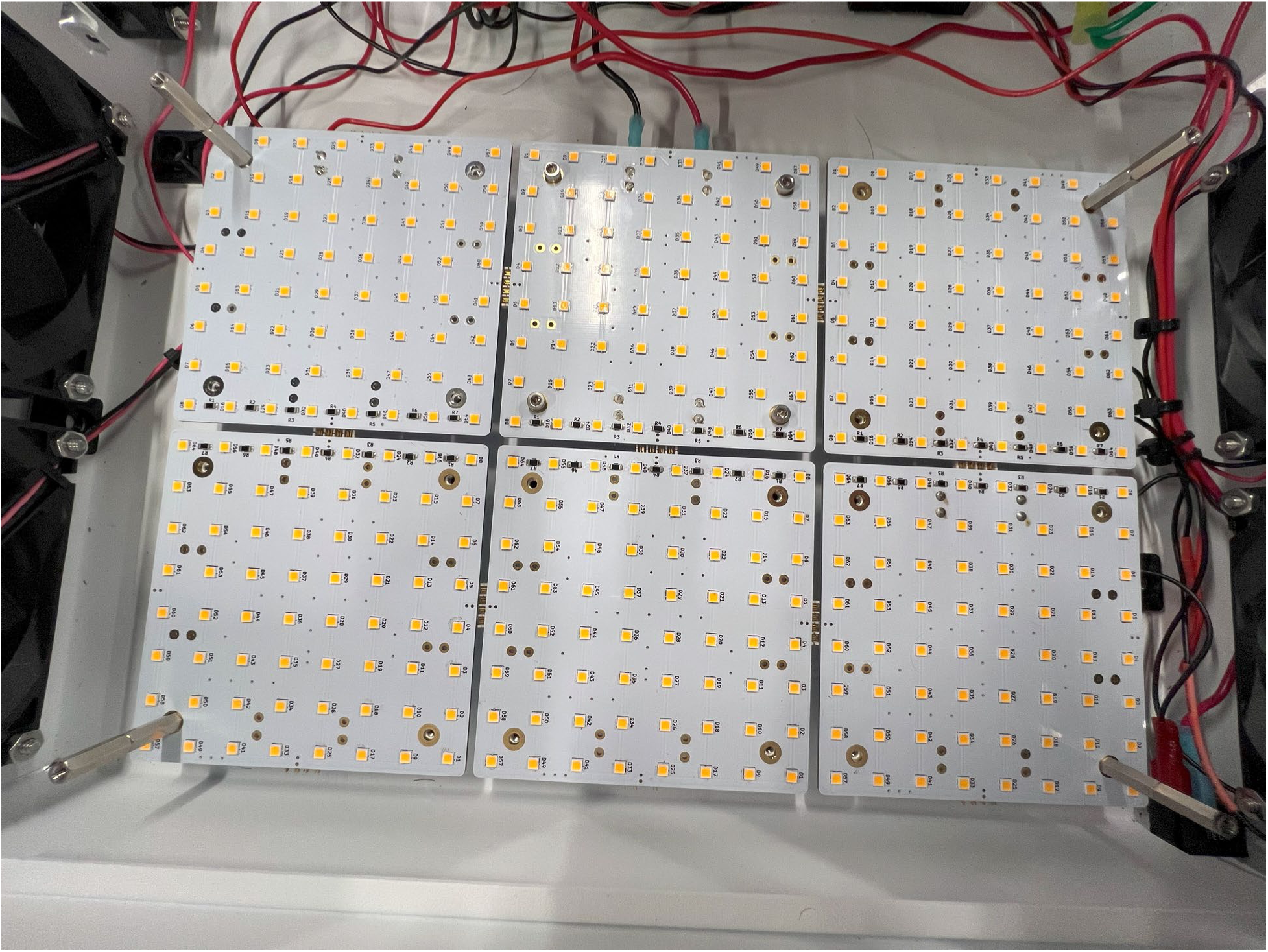
All Floor LEDs assembled and connected. The bulkhead panel is absent in this picture.

**Figure 27:**
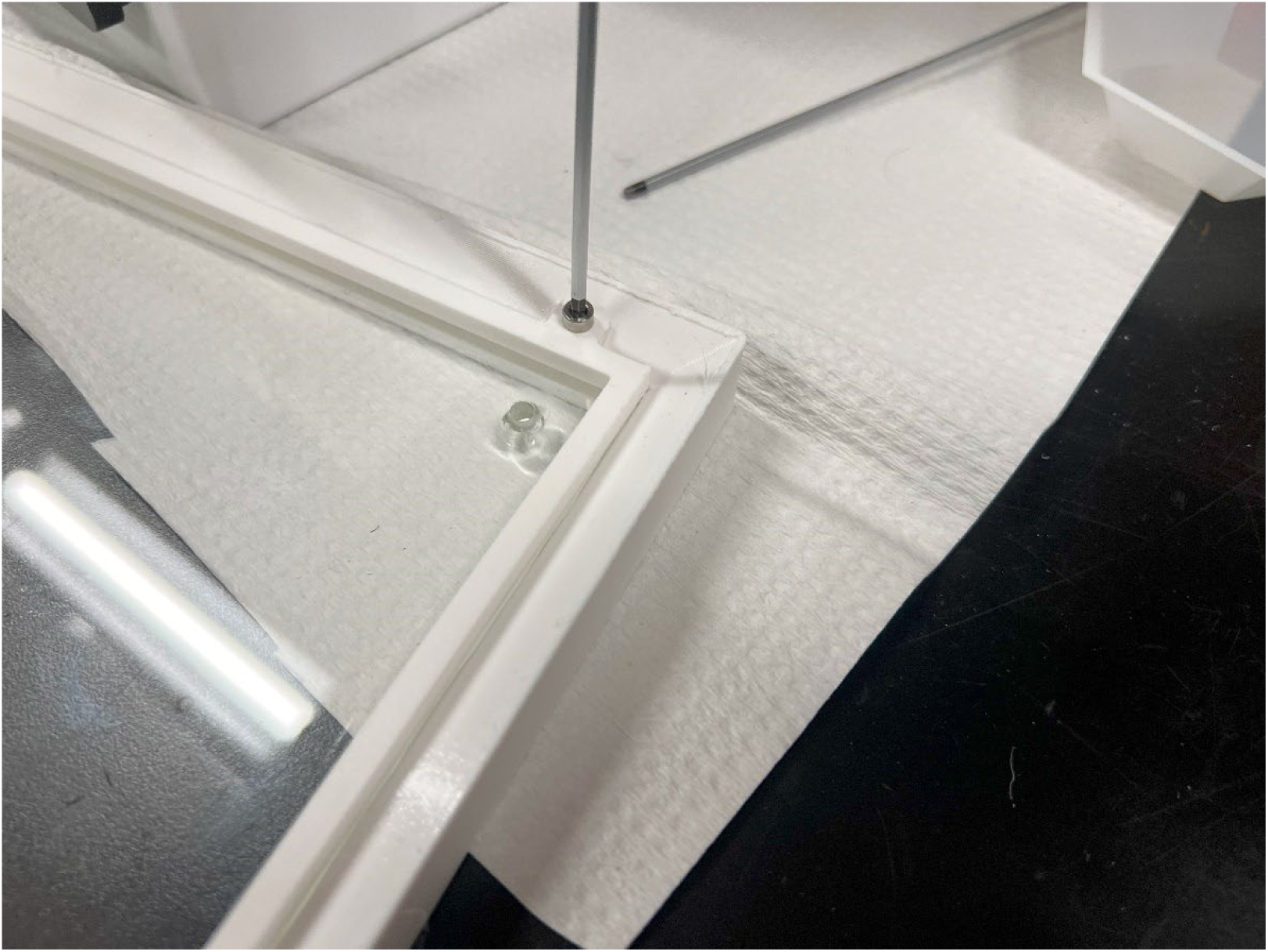
Installation and screw location for the bezel retainer.

**Figure 28:**
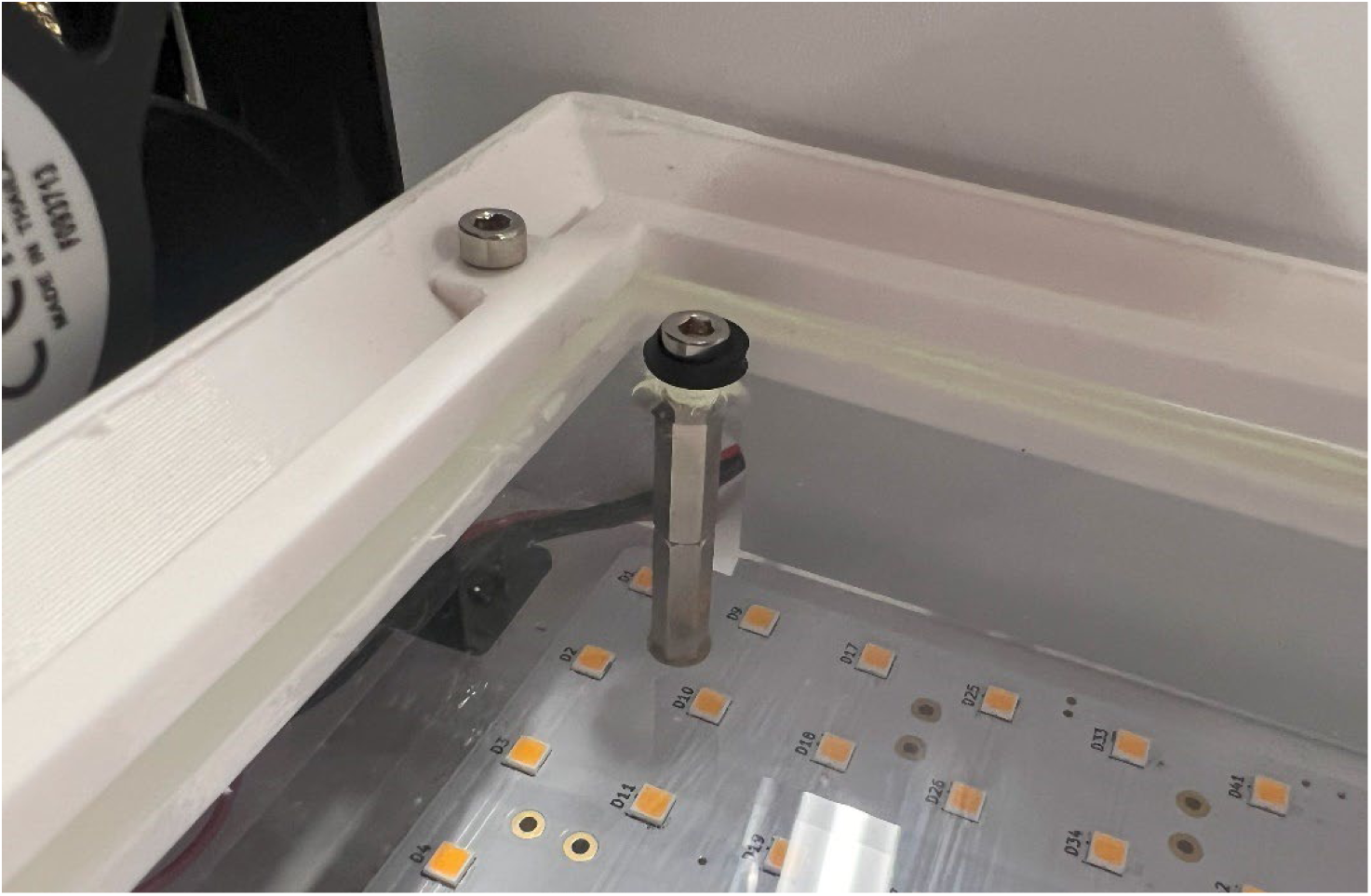
Mounting of the sample stage using rubber washers

**Figure 29:**
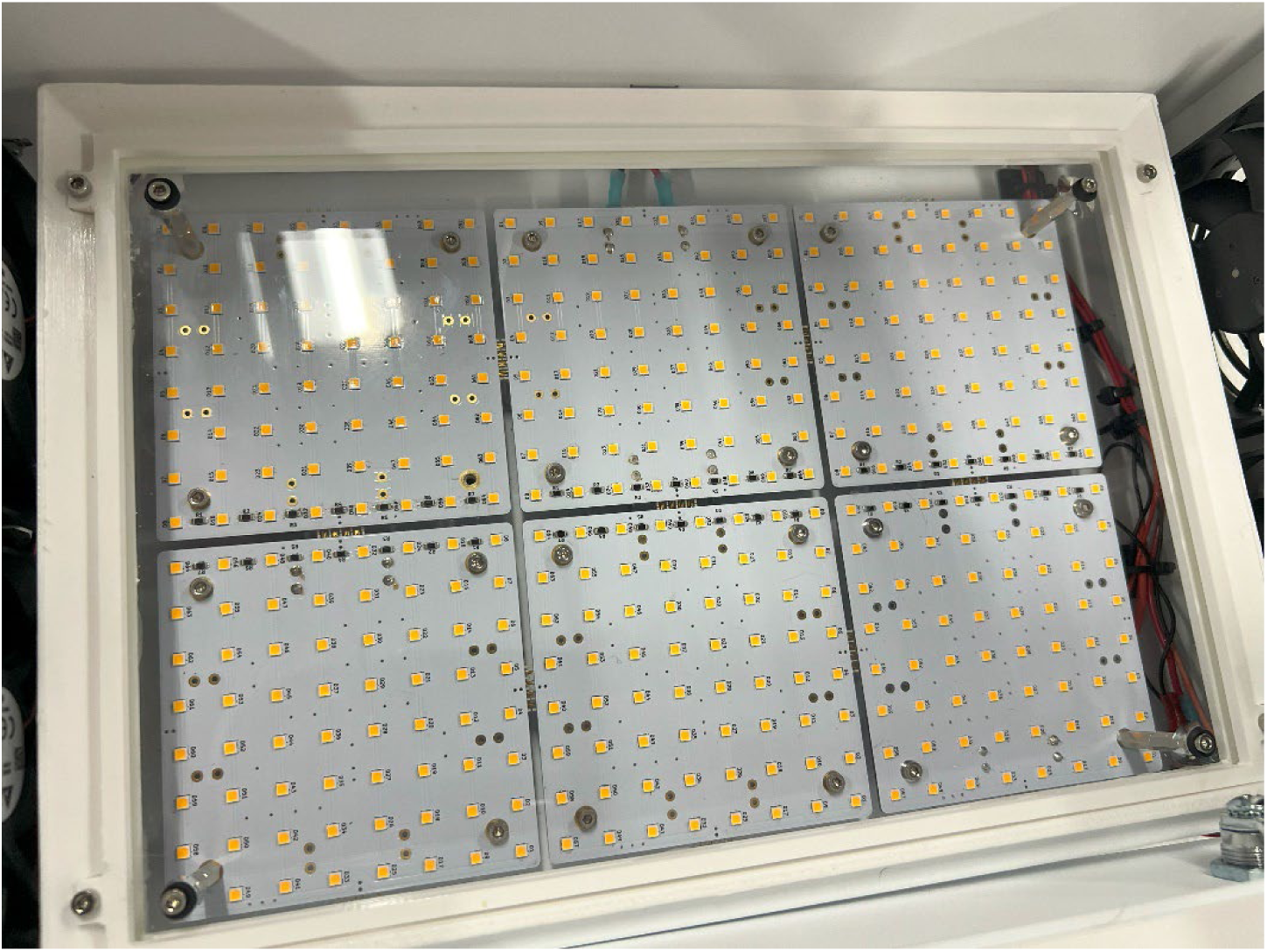
Sample Stage installed and mounted into the photobleacher

**Figure 30:**
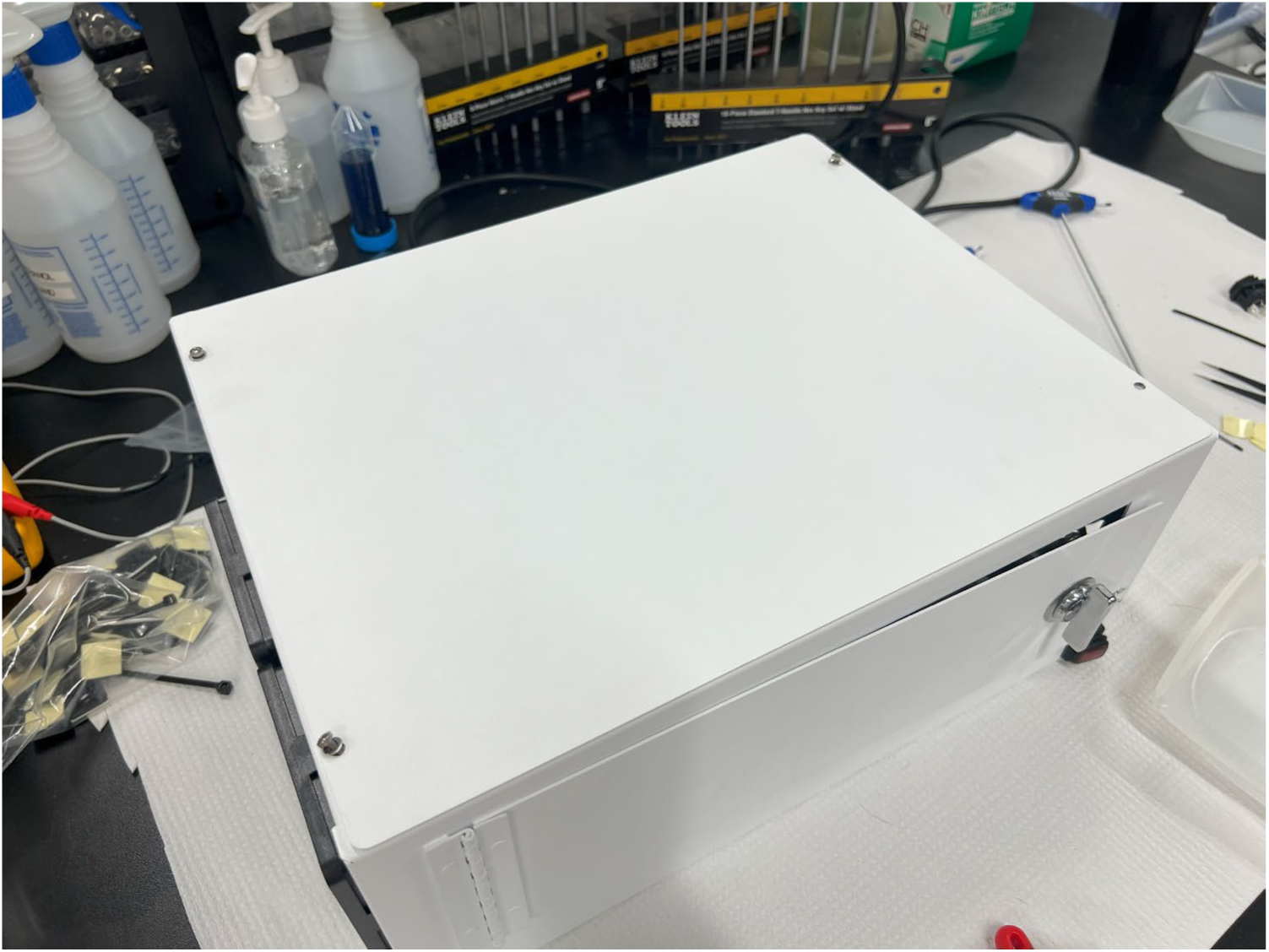
Fully assembled Photobleacher with Ceiling panel screwed in Once the ceiling is mounted, turn on the system again. Verify again that all fans and LEDs are on (you can look inside through the fan to double check). If everything is on, the system is ready for use.

**Table 1:** Required Tools.

| Name of Tool |
| --- |
| Philips-Head Screwdriver(s) |
| Allan Key Set (Metric) |
| Socket Wrench and/or Pliers |
| Crimping Tool |
| Wire Cutters and/or Wire Strippers |

**Table 2:**
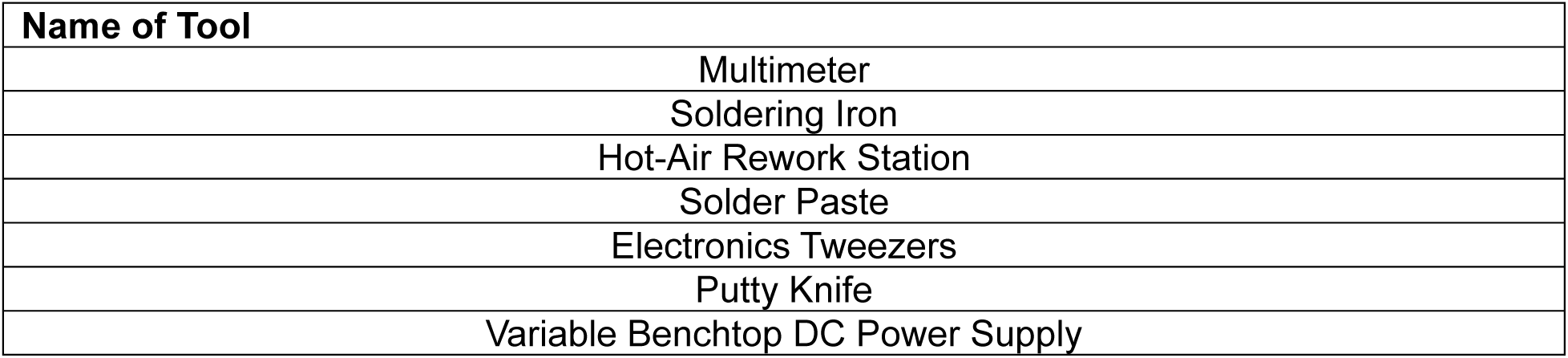
Optional Tools.

| Name of Tool |
| --- |
| Multimeter |
| Soldering Iron |
| Hot-Air Rework Station |
| Solder Paste |
| Electronics Tweezers |
| Putty Knife |
| Variable Benchtop DC Power Supply |

**Table 3:** Consumables.

| Name of Consumable |
| --- |
| Double-sided VHB tape |
| 16 AWG Stranded Wire (Red, Black, Green, Brown, Blue) |
| 18-24 AWG Stranded Wire (Red & Black) |
| SuperGlue |
| <u>Solder Paste</u> |

**Table 4:** Components Required for Section V.

| Quantity | Description | Manufacturer Part Number | Notes |
| --- | --- | --- | --- |
| 1 | AC/DC Converter<br>24V 200W | CUS200M-24 | Main DC Power Supply |
| 1 | PWR ENT MOD<br>RCPT IEC320-C14<br>PNL | KM01.1205.11 | AC Power input module, to be inserted into the back |
| 1 | Fuse Drawer for<br>Power input module:<br>F'GRIP 1PL | 4301.1405 | Insert fuse drawer, then insert fuse into Power Input Module |
| 1 | FUSE GLASS 2.5A<br>125VAC 5X20MM | 7010.345 | Fuse for Power Input Module |
| 1 | 100-Amp DualBus | 2702 | Terminal Bus Block for DC voltage signals |
| 1 | RELAY GEN<br>PURPOSE DPDT<br>10A 120V | LY2F-AC110/120 | Power Output Relay |
| 1 | CONN RCPT HSG<br>5POS 3.96MM | VHR-5N | AC Power input connector |
| 1 | CONN RCPT HSG<br>6POS 3.96MM | VHR-6N | DC Power Output Connector |
| 3 | CONN SOCKET 18-22AWG CRIMP TIN | SVH-21T-P1.1 | Crimp Contact for AC Input Connector |
| 6 | CONN SOCKET 16-20AWG CRIMP TIN | SVH-41T-P1.1 | Crimp Contact for DC Output Connector |
| 3 | CONN QC RCPT 14-16AWG 0.187 | 190030056 | Blue Quick-Connect: Female |
| 2 | CONN QC RCPT 18-22AWG 0.187 | 190030011 | Red Quick-Connect: Female |
| 2 | CONN RING CIRC<br>14-16AWG #8<br>CRIMP | 0190540110 | Ring Connectors for Terminal Block Bus |

**Table 5:** Components Required for Section VI.

| Quantity | Description | Manufacturer Part Number | Notes |
| --- | --- | --- | --- |
| 4 | FAN AXIAL<br>92X20MM 24VDC<br>WIRE | AFB0924HD | Main Stage Fans |
| 2 | FAN AXIAL<br>50X20MM 24VDC<br>WIRE | AFB0524VHD | AC Power input module, to<br>be inserted into the back |
| 2 | FILTR ASSB FAN<br>GUARD 92MM<br>45PPI | 4301.1405 | Inlet Fan Cover |
| 2 | FAN FILTER GUARD<br>ASSY 92MM ABS | LFG92 | Outlet Fan Cover |
| 2 | FAN GUARD 50MM<br>METAL | 08248 | PSU Fan Cover |
| 24 | M3 x 16 Philips Head<br>Screw |  | Mounting Screws for fan |
| 24 | M3 Nut |  | Nuts for mounting screws |
|  | Red Wire 18-24 AWG |  |  |
|  | Black Wire 18-24AWG |  |  |
| 2 - 6 | CONN RING CIRC<br>14-16AWG #8<br>CRIMP | 0190540110 | Ring Connectors for<br>Terminal Block Bus |

**Table 6:** Components Required for Section VII.

| Quantity | Description | Manufacturer Part Number | Notes |
| --- | --- | --- | --- |
| 1 | SWITCH ROCKER<br>SPST 16A 125V | KRE2ALA2RBB | Front Panel Rocker Switch |
| 1 | Magnet |  | Magnet on handle for reed switch |
| 1 | SENSOR REED SW<br>SPST-NO W LEADS | 59150-020 | Front Panel Reed Switch |
|  | SuperGlue |  |  |
|  | VHB tape |  |  |
|  | Red Wire 18-24 AWG |  |  |
|  | Black Wire 18-24AWG |  |  |

**Table 7:** Components Required for Section VIII & IX.

| Quantity | Description | Manufacturer<br>Part Number | Notes |
| --- | --- | --- | --- |
| 12 | LED PCB boards |  | LED boards fully assembled |
| 48 | M3 x 10 Screws |  | Mounting Screws for Boards |
| 4 | CONN QC RCPT 14-16AWG 0.187 | CONN QC<br>RCPT 14-16AWG 0.187 | Blue Quick-Connect: Female |
| 4 | CONN RING CIRC 14-16AWG #8 CRIMP | 0190540110 | Ring Connectors for Terminal Block Bus |
|  | Red Wire 16-18AWG |  | +24VDC wire |
|  | Black Wire 16-18AWG |  | -VDC (GND) wire |
| 4 | M3 Set Screw 10mm |  | Corner Floor Screws |
| 8 | M3 Standoffs 10mm |  | For Sample Stage Mounting |

## Safety Manual

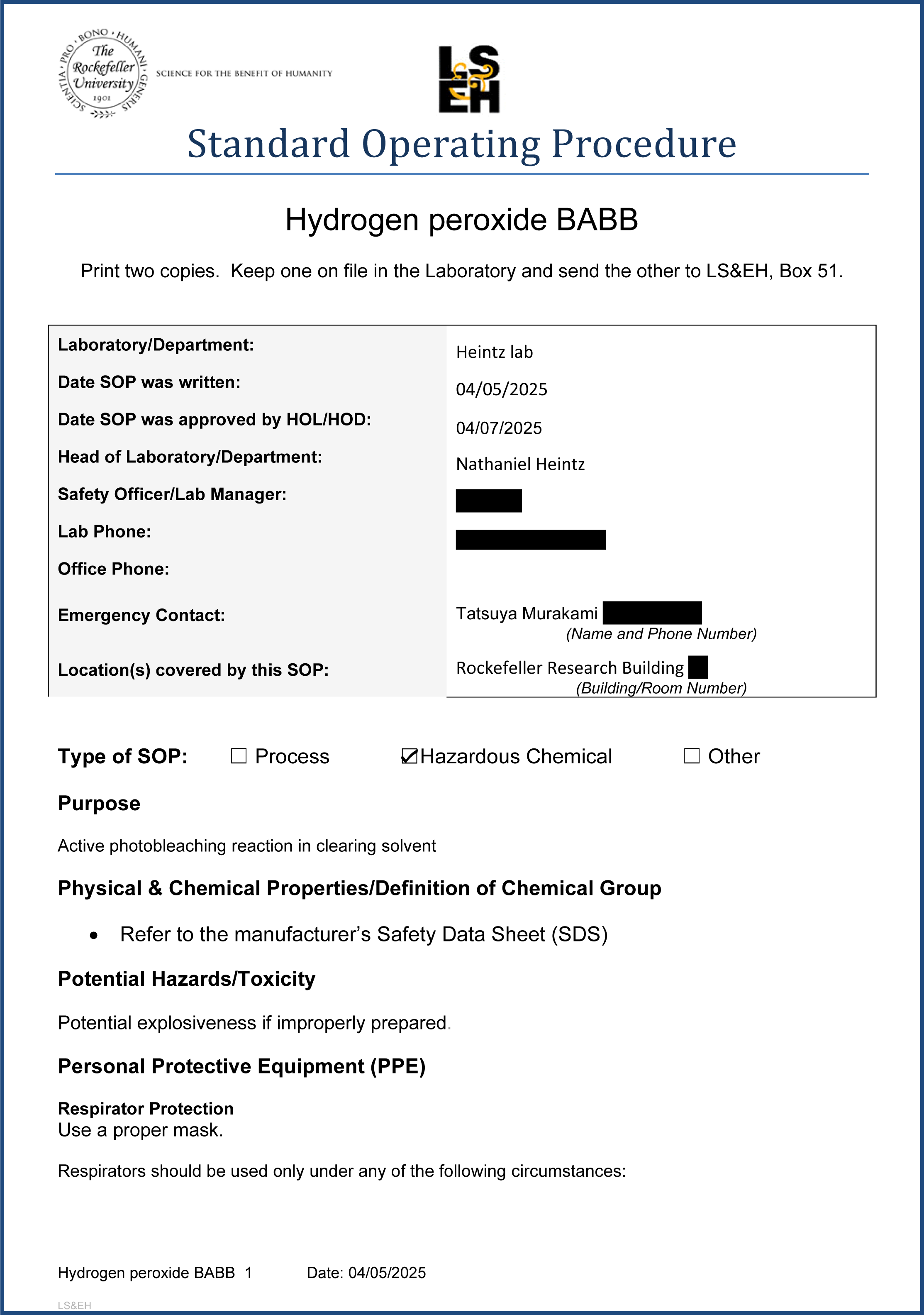
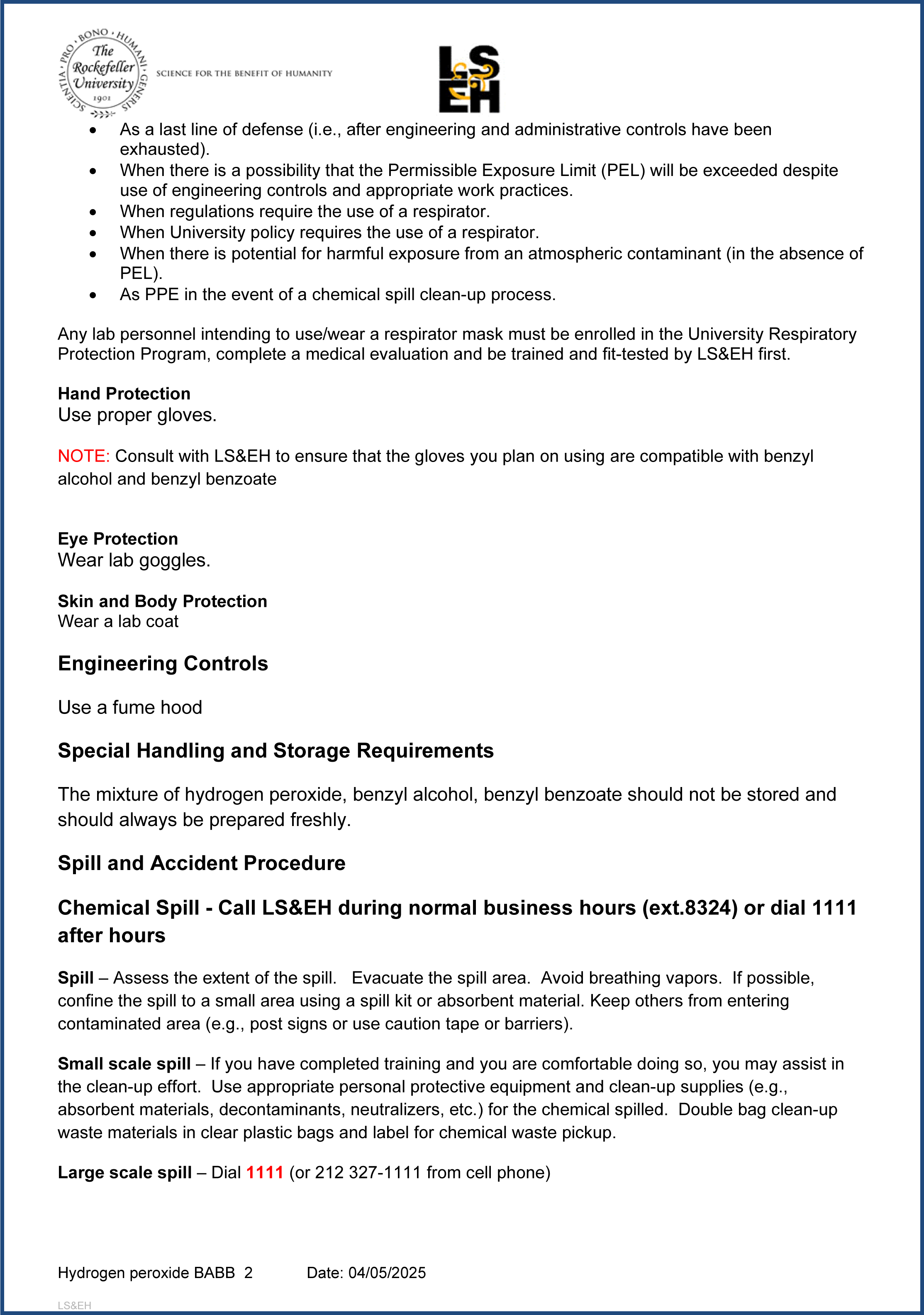
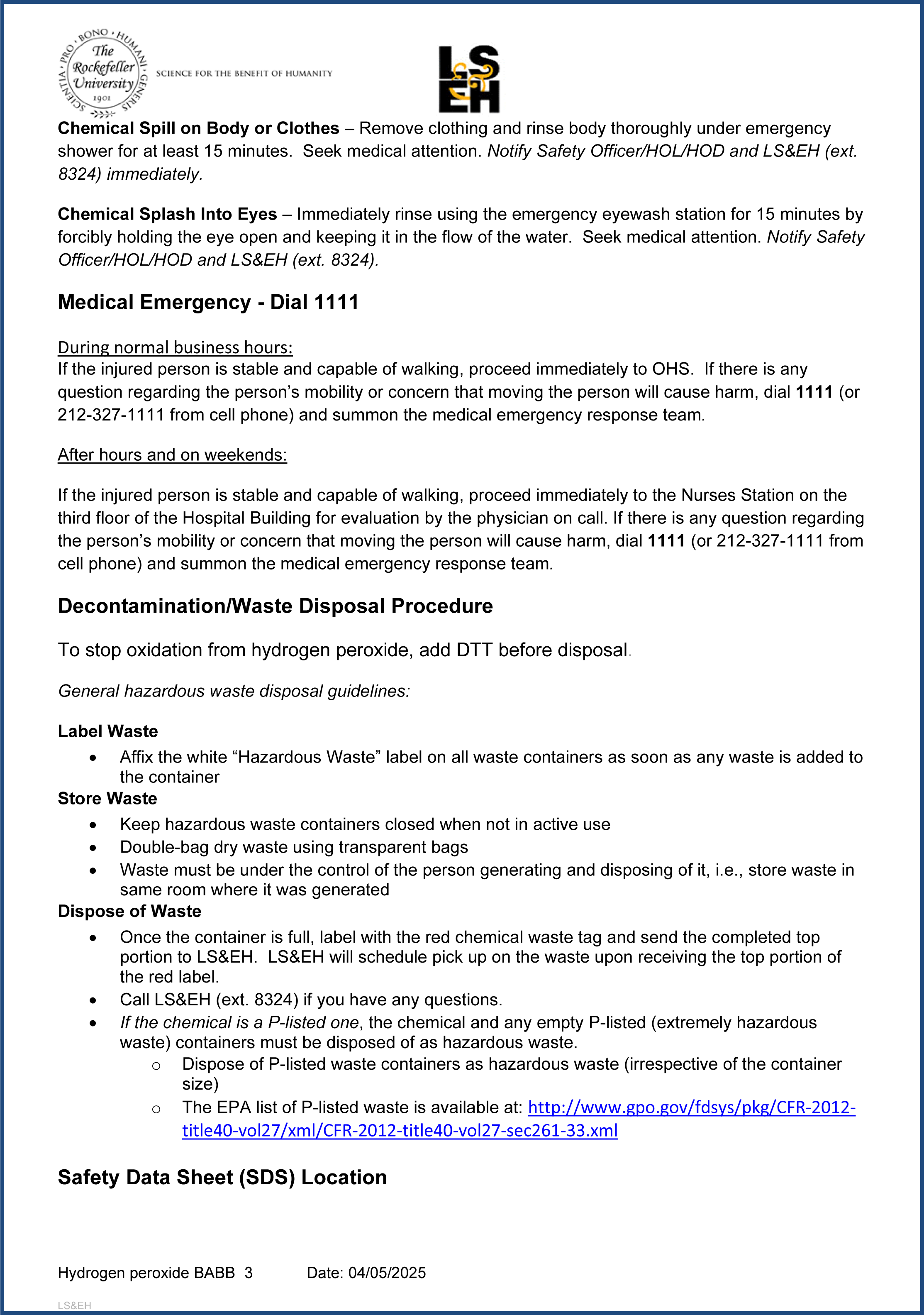
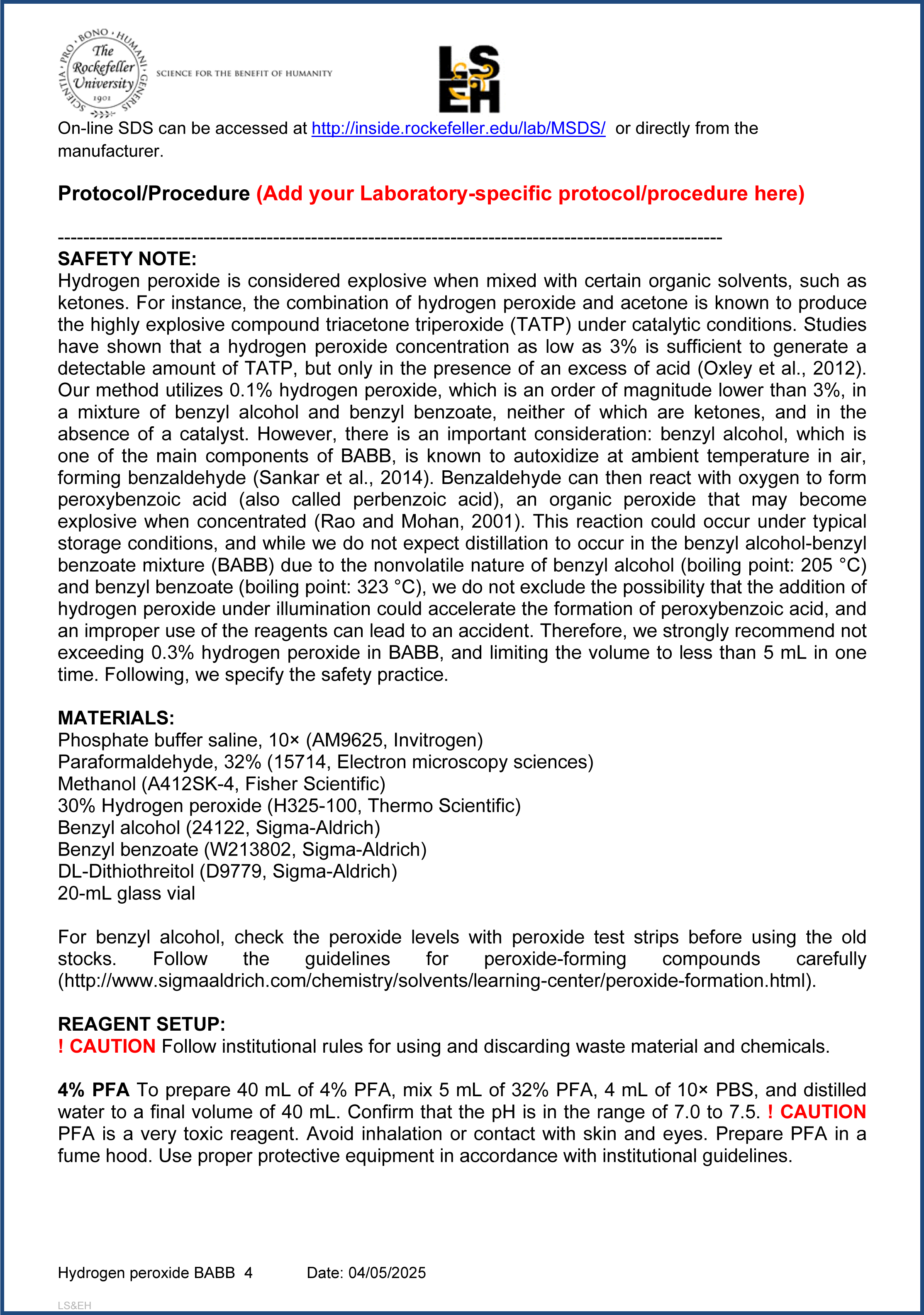
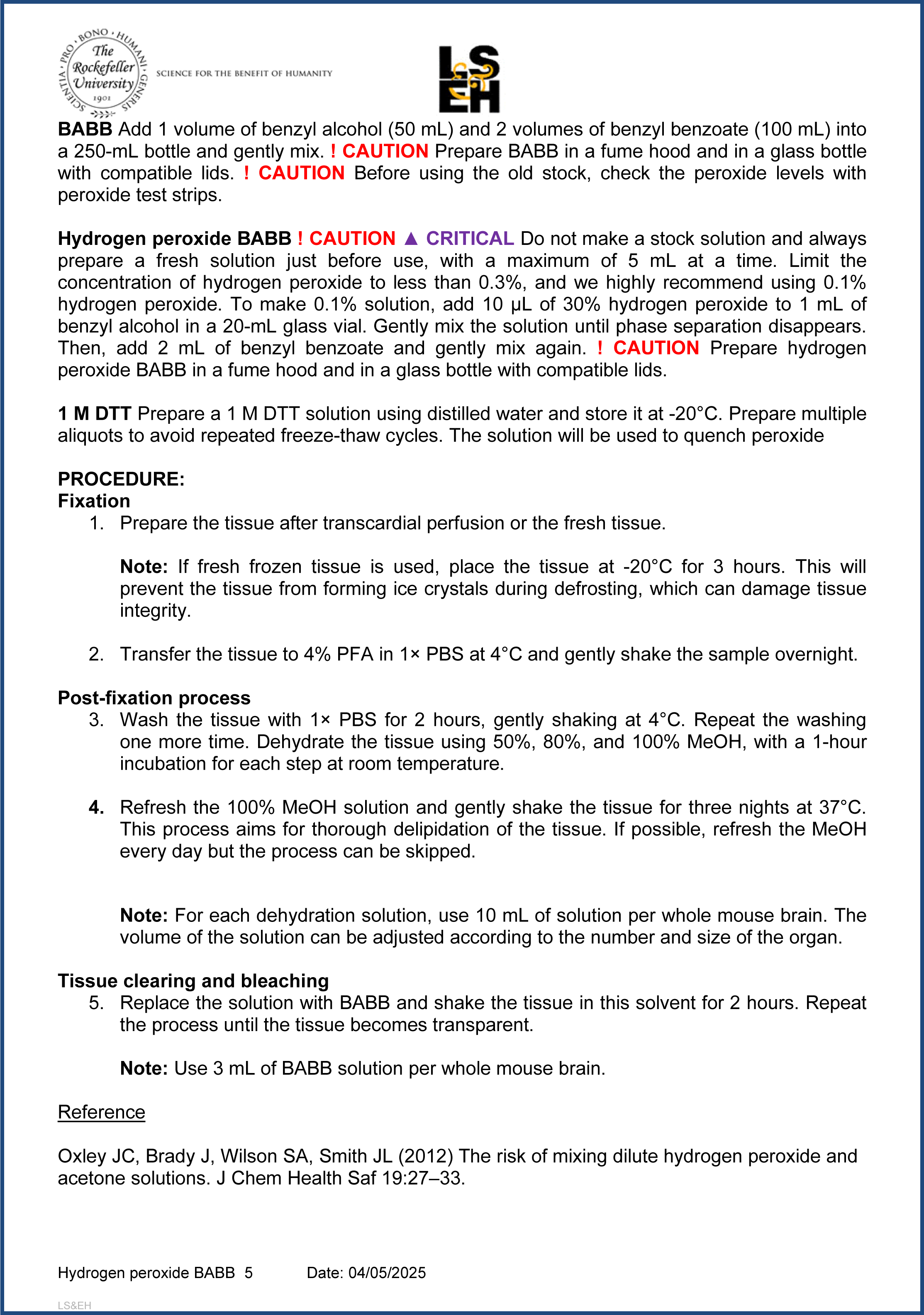
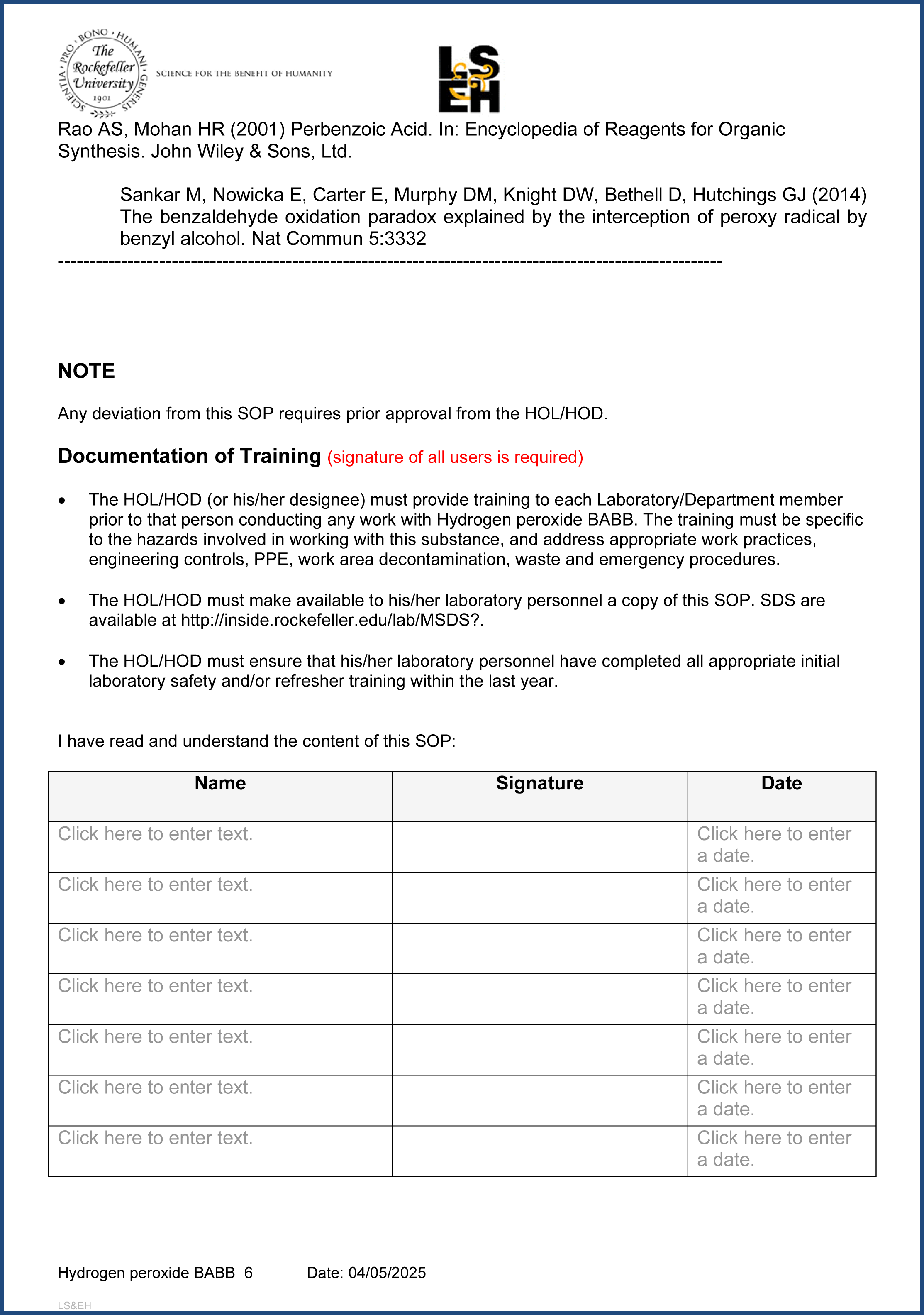
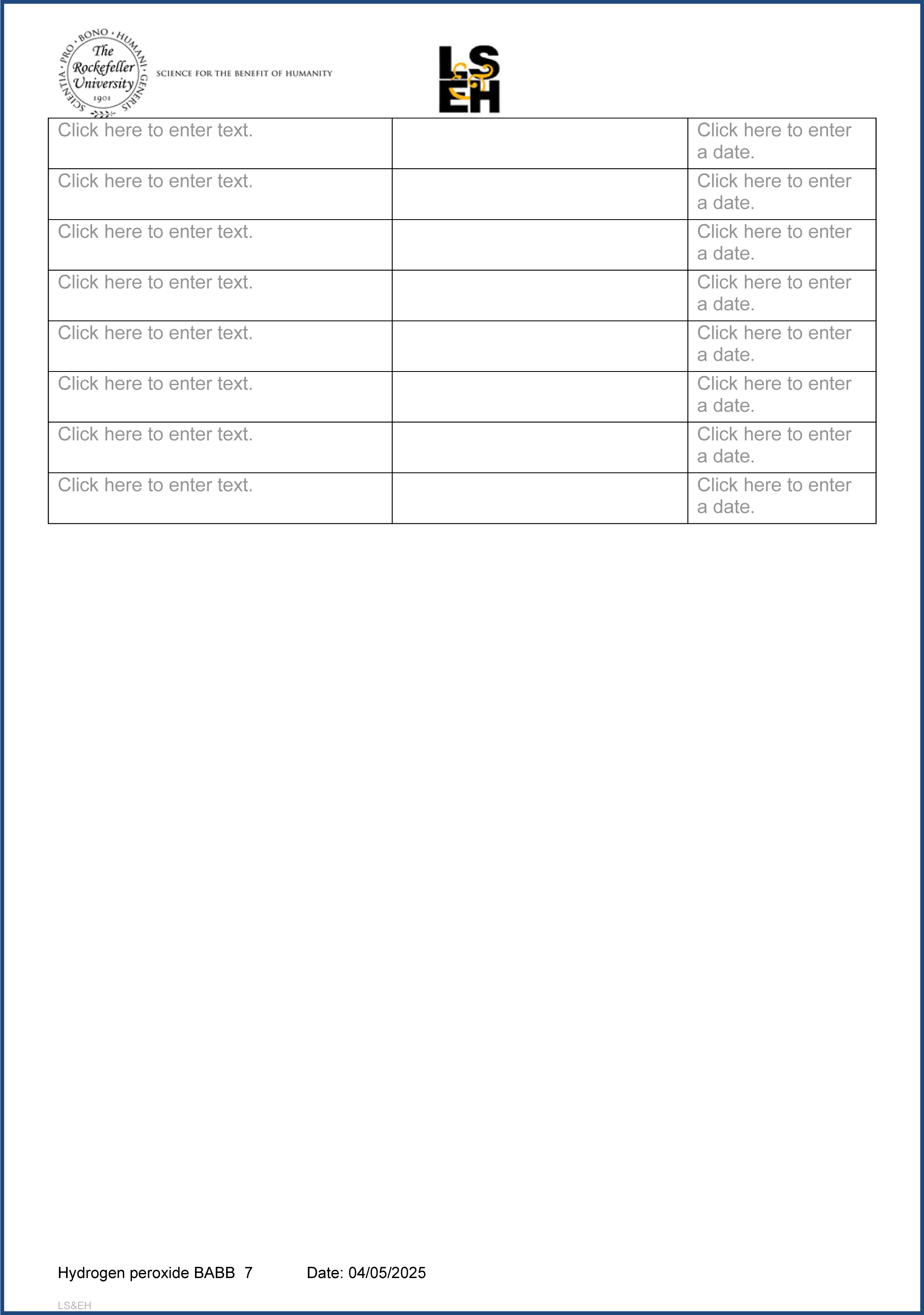

## REFERENCES

1. Baxendale JH, Wilson JA (1957) The photolysis of hydrogen peroxide at high light intensities. Trans Faraday Soc 53:344–356.

2. Chen KH, Boettiger AN, Moffitt JR, Wang S, Zhuang X (2015) Spatially resolved, highly multiplexed RNA profiling in single cells. Science 348:aaa6090.

3. Darche M, Borella Y, Verschueren A, Gantar I, Pagès S, Batti L, Paques M (2023) Light sheet fluorescence microscopy of cleared human eyes. Commun Biol 6:1–7.

4. Deschenes LA, Vanden Bout DA (2002) Single molecule photobleaching: increasing photon yield and survival time through suppression of two-step photolysis. Chemical Physics Letters 365:387–395.

5. Duong H, Han M (2013) A multispectral LED array for the reduction of background autofluorescence in brain tissue. Journal of Neuroscience Methods 220:46–54.

6. Gray DA, Woulfe J (2005) Lipofuscin and Aging: A Matter of Toxic Waste. Science of Aging Knowledge Environment 2005:re1–re1.

7. Hildebrand S, Schueth A, Herrler A, Galuske R, Roebroeck A (2019) Scalable Labeling for Cytoarchitectonic Characterization of Large Optically Cleared Human Neocortex Samples. Sci Rep 9:10880.

8. Ku T, Guan W, Evans NB, Sohn CH, Albanese A, Kim J-G, Frosch MP, Chung K (2020) Elasticizing tissues for reversible shape transformation and accelerated molecular labeling. Nat Methods 17:609–613.

9. Lai HM, Liu AKL, Ng HHM, Goldfinger MH, Chau TW, DeFelice J, Tilley BS, Wong WM, Wu W, Gentleman SM (2018) Next generation histology methods for three- dimensional imaging of fresh and archival human brain tissues. Nat Commun 9:1066.

10. Liu AKL, Hurry MED, Ng O t. W, DeFelice J, Lai HM, Pearce RKB, Wong GT-C, Chang RC-C, Gentleman SM (2016) Bringing CLARITY to the human brain: visualization of Lewy pathology in three dimensions. Neuropathology and Applied Neurobiology 42:573–587.

11. Lubeck E, Coskun AF, Zhiyentayev T, Ahmad M, Cai L (2014) Single-cell in situ RNA profiling by sequential hybridization. Nat Methods 11:360–361.

12. Mai H, Rong Z, Zhao S, Cai R, Steinke H, Bechmann I, Ertürk A (2022) Scalable tissue labeling and clearing of intact human organs. Nat Protoc:1–35.

13. Mann DMA, Yates PO, Stamp JE (1978) The relationship between lipofuscin pigment and ageing in the human nervous system. Journal of the Neurological Sciences 37:83–93.

14. Morawski M, Kirilina E, Scherf N, Jäger C, Reimann K, Trampel R, Gavriilidis F, Geyer S, Biedermann B, Arendt T, Weiskopf N (2018) Developing 3D microscopy with CLARITY on human brain tissue: Towards a tool for informing and validating MRI-based histology. NeuroImage 182:417–428.

15. Murray E, Cho JH, Goodwin D, Ku T, Swaney J, Kim S-Y, Choi H, Park Y-G, Park J- Y, Hubbert A, McCue M, Vassallo S, Bakh N, Frosch MP, Wedeen VJ, Seung HS, Chung K (2015) Simple, Scalable Proteomic Imaging for High-Dimensional Profiling of Intact Systems. Cell 163:1500–1514.

16. Neumann M, Gabel D (2002) Simple Method for Reduction of Autofluorescence in Fluorescence Microscopy. J Histochem Cytochem 50:437–439.

17. Nojima S, Susaki EA, Yoshida K, Takemoto H, Tsujimura N, Iijima S, Takachi K, Nakahara Y, Tahara S, Ohshima K, Kurashige M, Hori Y, Wada N, Ikeda J, Kumanogoh A, Morii E, Ueda HR (2017) CUBIC pathology: three-dimensional imaging for pathological diagnosis. Sci Rep 7:9269.

18. Park J et al. (2024) Integrated platform for multiscale molecular imaging and phenotyping of the human brain. Science 384:eadh9979.

19. Pigoli C, Gibelli LR, Caniatti M, Moretti L, Sironi G, Giudice C (2019) Bleaching melanin in formalin-fixed and paraffin-embedded melanoma specimens using visible light: a pilot study. European Journal of Histochemistry 63.

20. Renier N, Wu Z, Simon DJ, Yang J, Ariel P, Tessier-Lavigne M (2014) iDISCO: A Simple, Rapid Method to Immunolabel Large Tissue Samples for Volume Imaging. Cell 159:896–910.

21. Shinde AV, Humeres C, Frangogiannis NG (2017) The role of α-smooth muscle actin in fibroblast-mediated matrix contraction and remodeling. Biochimica et Biophysica Acta (BBA) - Molecular Basis of Disease 1863:298–309.

22. Sun Y, Chakrabartty A (2016) Cost-effective elimination of lipofuscin fluorescence from formalin-fixed brain tissue by white phosphor light emitting diode array. Biochem Cell Biol 94:545–550.

23. Susaki EA et al. (2020) Versatile whole-organ/body staining and imaging based on electrolyte-gel properties of biological tissues. Nat Commun 11:1982.

24. Tainaka K et al. (2018) Chemical Landscape for Tissue Clearing Based on Hydrophilic Reagents. Cell Reports 24:2196–2210.e9.

25. Tsuneoka Y, Atsumi Y, Makanae A, Yashiro M, Funato H (2022) Fluorescence quenching by high-power LEDs for highly sensitive fluorescence in situ hybridization. Front Mol Neurosci 15.

26. Widengren J, Rigler R (1996) Mechanisms of photobleaching investigated by fluorescence correlation spectroscopy. Bioimaging 4:149–157.

27. Zhao S et al. (2020) Cellular and Molecular Probing of Intact Human Organs. Cell 180:796–812.e19.

28. Zheng J, Wu Y-C, Phillips EH, Cai X, Wang X, Seung-Young Lee S (2024) Increased Multiplexity in Optical Tissue Clearing-Based Three-Dimensional Immunofluorescence Microscopy of the Tumor Microenvironment by Light-Emitting Diode Photobleaching. Laboratory Investigation 104:102072.

